# Antagonistic inhibitory subnetworks control cooperation and competition across cortical space

**DOI:** 10.1101/2021.03.31.437953

**Authors:** Daniel P. Mossing, Julia Veit, Agostina Palmigiano, Kenneth D. Miller, Hillel Adesnik

## Abstract

The cortical microcircuit can dynamically adjust to dramatic changes in the strength, scale, and complexity of its input. In the primary visual cortex (V1), pyramidal cells (PCs) integrate widely across space when signals are weak, but narrowly when signals are strong, a phenomenon known as contrast-dependent surround suppression. Theoretical work has proposed that local interneurons could mediate a shift from cooperation to competition of PCs across cortical space, underlying this computation. We combined calcium imaging and electrophysiology to constrain a stabilized supralinear network model that explains how the four principal cell types in layer 2/3 (L2/3) of mouse V1– somatostatin (SST), parvalbumin (PV), and vasoactive intestinal peptide (VIP) interneurons, and PCs– transform inputs from layer 4 (L4) PCs to encode drifting gratings of varying size and contrast. Using bidirectional optogenetic perturbations, we confirmed key predictions of the model. Our data and modeling showed that recurrent amplification drives a transition from a positive PC→VIP⊣SST⊣PC feedback loop at small size and low contrast to a negative PC→SST⊣PC feedback loop at large size and high contrast to contribute to this flexible computation. This may represent a widespread mechanism for gating competition across cortical space to optimally meet task demands.

## Introduction

The brain must process sensory inputs in their spatial context across a wide range of sensory environments and perceptual goals. When sensory signals are weak, adding activity in ensembles encoding congruent evidence across space (“corroboration”) could optimize detection; when signals are strong, subtraction of congruent signals (“explaining away”) could optimize discrimination or localization (Hawken et al., 2001; Nauhaus et al., 2009; Sceniak et al., 1999). To enable this flexibility, neural ensembles encoding distinct elements of sensory evidence must be able to rapidly change the nature of their interaction, cooperating or competing depending on sensory inputs.

Prior literature suggests that with increasing contrast, interactions across cortical space shift from cooperative to competitive (Nauhaus et al., 2009; Sato et al., 2014). In the primary visual cortex (V1), competition across cortical space likely contributes to surround suppression, in which cortical pyramidal cells (PCs) fire more weakly in response to a large visual stimulus centered on their receptive field than to a small stimulus (Angelucci et al., 2017). Conversely, cooperation across cortical space might contribute to surround facilitation, in which PC responses increase in response to large stimuli. Crucially, surround suppression is dominant at high contrast, while surround facilitation is more prevalent at low contrast (Anderson et al., 2001; Ayaz et al., 2013; Cavanaugh, Bair, & Anthony Movshon, 2002; Cavanaugh, Bair, & Movshon, 2002; Ichida et al., 2007; Nienborg et al., 2013; Olsen et al., 2012; Ozeki et al., 2009; Polat et al., 1998; Sceniak et al., 1999; Schwabe et al., 2010; Sengpiel et al., 1997; Tanaka & Ohzawa, 2009; Vaiceliunaite et al., 2013; Van Den Bergh et al., 2010; C. Wang et al., 2009). Theoretical work has proposed that local inhibition mediates the transition from cooperative (surround facilitating) to competitive (surround suppressive) spatial interactions with increasing contrast, either via a class of interneurons that do not respond to low contrasts (Schwabe, et al. 2010, Angelucci, et al. 2017) or via nonlinear network dynamics that increase the relative role of inhibition with increasing contrast (Rubin et al., 2015). In this study, we determine how a cortical microcircuit of genetically defined interneuron subtypes underlies this shift from cooperation to competition across cortical space in mouse V1.

Three interneuron subtypes, parvalbumin- (PV), somatostatin- (SST) and vasoactive intestinal peptide- (VIP) expressing, comprise 80-85% of cortical interneurons (Pfeffer et al., 2013). All three subtypes have been found to shape the sensory coding properties of PCs in distinct ways to facilitate cortical encoding of sensory features (Adesnik et al., 2012; Ayzenshtat et al., 2016; S.-H. Lee et al., 2012; Rudy et al., 2011). VIP cells specialize in inhibition of other interneurons (S. Lee et al., 2013; Pfeffer et al., 2013; Pi et al., 2013) and may thus play a crucial role in promoting PC cooperation over competition. VIP cells receive feedback excitation from higher areas (Zhang et al., 2014) and strong neuromodulatory input from the midbrain (Fu et al., 2014) which positions them to mediate external control of flexible cooperation and competition in the cortex (X.-J. Wang & Yang, 2018; Yang et al., 2016). Recent work has shown that VIP cells, like PCs and PV cells, are highly surround suppressed (Dipoppa et al., 2018), are sensitive to low contrasts (Millman et al., 2020), and prefer spatially heterogeneous as opposed to homogeneous textures (Keller et al., 2020). VIP cells preferentially innervate SST cells (Pfeffer et al., 2013), which play an important role in spatial competition between PCs (Adesnik et al., 2012). Thus, we hypothesized that VIP cells might contribute to the contrast dependence of cooperation and competition across cortical space, by selectively inhibiting SST cells at low contrast to support cooperation.

To test this idea, we combined neural recording, data-driven mechanistic modeling, and optogenetic perturbations in layer 2/3 (L2/3) of mouse V1. Our model predicted network mechanisms underlying the shift from cooperation to competition across cortical space, and we used optogenetic perturbations to validate the model by testing its predictions. A salient emergent property of the model, borne out by experiment, was that much of the effect of VIP optogenetic perturbations on PC activity could be parsimoniously explained by PC activity level in the absence of perturbation, regardless of the sensory stimulus. Because a linear function (with a multiplicative and additive factor) captured this dependence, we referred to this component of the optogenetic effect as the “linearly predicted” component. The linearly predicted component did not change the preferred size at a given contrast, and thus did not affect contrast dependence of surround suppression. Crucially, our experiments also revealed a component of the impact of these perturbations that depended specifically on the visual stimulus and was not captured by a purely linear effect; we termed this the “residual” component. This residual component changed preferred size and significantly altered the contrast dependence of surround suppression in PCs. Importantly, the mechanistic model predicted an asymmetry between the effects of VIP activation and silencing on PCs: for VIP activation, but not silencing, it predicted large residual changes in PC activity, resulting in an enhancement of contrast-dependent surround suppression. These predictions were likewise borne out by experiment.

Taken together, our experiments and modeling showed that VIP and SST activity outlined disparate regimes of sensory coding in PCs. When the network was weakly activated by stimuli of low contrast and small size, VIP cells dominated. Here, increasing PC activity drove VIP cells, which in turn inhibited SST cells to disinhibit PCs, closing a positive PC→VIP⊣SST⊣PC feedback loop in L2/3. PCs in turn showed surround facilitation and were highly sensitive to changes in contrast. Conversely, in the SST-dominated regime, when the network was strongly activated by stimuli of high contrast and large size, increasing PC activity instead drove SST cells, which in turn inhibited PCs, closing a negative PC→SST⊣PC feedback loop. Positive feedback within the PC-PV network amplified the effects of both feedback loops. Responses to optogenetic perturbations showed signatures of these feedback loops in the form of stimulus-dependent residual effects, after accounting for the component that depended on PC activity in the absence of optogenetic perturbation. Although VIP silencing experiments showed that these feedback loops were not necessary for contrast-dependent surround suppression, VIP activation showed that they could powerfully enhance it. This model-based approach to predicting and interpreting the effects of perturbations may prove to be broadly applicable to understanding network-level interactions in recurrently connected neural circuits.

## Results

### Cell-type specific encoding of size and contrast in the mouse primary visual cortex

To study a paradigmatic example of flexible cooperation and competition across cortical space, we first probed the logic of contrast-dependent surround suppression in layer 2/3 pyramidal cells (L2/3 PCs) of the primary visual cortex (V1). We recorded neural activity in awake, head-fixed mice (fig. 1a) using cell type-specific calcium imaging (fig. 1b, see Methods). After mapping each neuron’s receptive field, we independently varied the size, contrast, and direction of drifting gratings while keeping their location constant. We measured responses using deconvolved event rate (fig. 1c, see Methods), and analyzed all neurons whose cell bodies lay within 10° of retinotopic space (corresponding to roughly one receptive field width) of the stimulus representation center, on trials in which the animal was not locomoting (see Methods). To specifically examine tuning for size and contrast, we averaged responses across directions.

**Figure 1.**
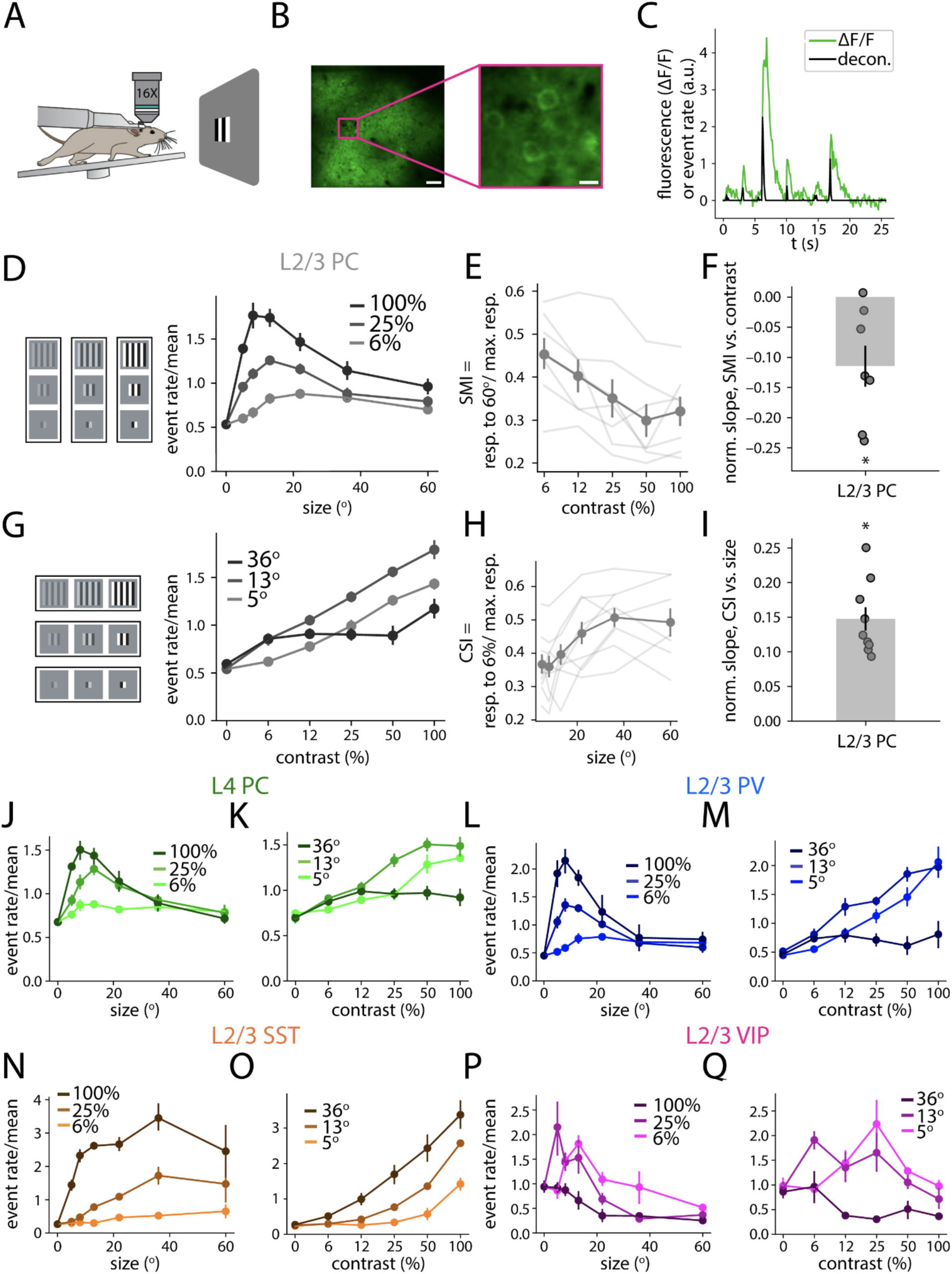
Cell-type specific responses to stimulus size and contrast in mouse V1. A) Experiment schematic: Mice were head-fixed and free to run on a circular treadmill, while passively viewing a monitor showing drifting gratings of varying size, contrast, and direction. B) Example image from a Camk2a-tTA;tetO-GCaMP6s mouse (left), with zoomed in inset (right). Left scale bar corresponds to 50 µm, and right scale bar to 10 µm. C) Example calcium trace (green) and deconvolved event rate (black) which was used to estimate neural activity. D) Left: schematic of the visual stimuli used in this study. Right: averaged deconvolved event rates in L2/3 PCs plotted against drifting grating patch size, with different contrasts plotted separately (*n* = 9 imaging sessions). Event rates are normalized to the mean across all stimuli and all neurons within each imaging session, in this and all following panels. In all size tuning plots, darker color corresponds to higher contrast. E) Surround modulation index (SMI), defined as the evoked (baseline-subtracted) response to 60° gratings divided by the response to the preferred size for each contrast, plotted against grating contrast, for L2/3 PCs. Here and elsewhere, each transparent line corresponds to an imaging session, and the solid line corresponds to the average across sessions (note that SMI could be computed only for the 7 imaging sessions for which 60° size stimuli were shown). F) The slope of SMI vs. contrast, normalized to the slope corresponding to SMI of 0 at the smallest size and 1 at the largest size. The SMI slope corresponds to the magnitude of contrast dependence of surround suppression (slope significantly negative, p<0.05, Wilcoxon signed-rank test). Here and elsewhere, dots indicate imaging sessions, and bar with error bar indicate mean +/- bootstrap SEM across imaging sessions. G) The same tuning curves as in D) are plotted, separated by size, with contrast varying along the x-axis. In all contrast tuning plots, darker color corresponds to larger size. H) Contrast sensitivity index (CSI), defined as the evoked response to 6% contrast gratings divided by the response to the preferred contrast for each size, plotted against grating size, for L2/3 PCs. I) The slope of CSI vs. contrast, normalized to the slope corresponding to CSI of 0 at the smallest size and 1 at the largest size. The CSI slope corresponds to the magnitude of size dependence of contrast sensitivity (slope significantly positive, p < 0.01, Wilcoxon signed-rank test). J-Q) Same as D), G), for L4 PCs (*n* = 6), L2/3 PV cells (*n* = 6), L2/3 SST cells (*n* = 5), and L2/3 VIP cells (*n* = 5), respectively. * significantly different from zero, p < 0.05, Wilcoxon signed rank test. Error bars indicate bootstrap SEM equivalent.

We first asked how size tuning varied as a function of contrast. As in prior electrical recordings from unidentified cell types in monkeys (Cavanaugh, Bair, & Movshon, 2002; Kapadia et al., 1999; Levitt & Lund, 1997; Sceniak et al., 1999), cats (Polat et al., 1998; Sengpiel et al., 1997; Toth et al., 1996), and mice (Ayaz et al., 2013; Nienborg et al., 2013; Vaiceliunaite et al., 2013), we found that at high contrast, L2/3 PC responses showed highly non-monotonic size tuning, peaking sharply at small sizes, while at low contrast, responses peaked more broadly, and at larger sizes (fig. 1d). Because larger stimuli drive activity in distal (surrounding) regions of cortical space, this is consistent with a picture in which surround PCs cooperate with center PCs at low contrast (supporting broad size tuning and preference for larger sizes) and compete with center PCs at high contrast (supporting sharp size tuning and preference for smaller sizes).

To quantify surround suppression as a function of contrast, we defined a surround modulation index (SMI) at a given contrast as the evoked (baseline-subtracted) response to 60° gratings (the largest shown), divided by the response to the preferred size. This metric of surround suppression increased in strength with increasing contrast (fig. 1e; fig. 1f shows slope < 0, p < 0.05, Wilcoxon signed-rank test). Surround facilitation at low contrast has been theorized to enhance detectability of large textures (Hawken et al., 2001; Sceniak et al., 1999). If this were true, we would expect to see sensitivity to low contrast stimuli increase with size. We measured this change in sensitivity directly by taking a complementary view of the same data, examining contrast tuning at each size (fig. 1g). We defined a contrast sensitivity index (CSI) at a given size as the evoked response to 6% contrast (the lowest shown), divided by the response to the preferred contrast. This metric of contrast sensitivity increased with increasing size (fig. 1h; fig. 1i shows slope > 0, p < 0.01, Wilcoxon signed-rank test). These findings were robust across behavioral states (fig. S1) and corroborated by analogous electrophysiological measurements (fig. S2).

To understand the network mechanisms that might control contrast dependence of surround suppression in L2/3 PCs, we next probed the tuning of other cell types in the circuit. We focused first on their primary excitatory inputs, L4 PCs. In fact, L4 PCs on average showed similar size and contrast tuning (fig. 1j,k). Thus, feedforward inputs provided a basis for contrast-dependent surround suppression in L2/3 PCs (the phenomenon has been reported as early as retinal ganglion cells (Nolt et al., 2004)), but this does not preclude inputs within L2/3 also playing a role. Thus, we examined the principal inhibitory inputs to L2/3 PCs, L2/3 inhibitory interneurons. Prior theoretical studies have proposed that local interneurons could support flexible cooperation and competition of PCs, either via an inhibitory cell type that is selectively driven at high contrasts (Angelucci et al., 2017), or via nonlinear dynamical mechanisms (Rubin et al., 2015). We first asked whether L2/3 PV cells, which share inputs with and strongly inhibit L2/3 PCs, might show enhanced selectivity for high contrast relative to L2/3 PCs. However, we found that in keeping with their high degree of shared inputs, tuning of PV cells closely matched that of PCs (fig. 1l,m). Activity measurements of fast spiking (FS) units recorded using extracellular electrophysiology agreed with these calcium imaging measurements (fig. S2; non-locomoting, FS units, R=0.84, p=7×10^-9^, Wald test).

Therefore, we next examined a second major L2/3 interneuron subtype, SST cells. Previous work has implicated SST cells in contributing to surround suppression of PCs at high contrast (Adesnik et al., 2012; Nienborg et al., 2013). Although SST cells are known to prefer larger sizes than the other cortical cell types, how this depends on stimulus contrast is not known. In fact, SST tuning diverged sharply from that of PCs and PV cells. SST cells responded best at large size and high contrast, and weakly or not at all at small size and low contrast (fig. 1n); small size, intermediate contrast stimuli that drove nearly maximal activity in PCs and PV cells only weakly drove SST cells (fig. 1o). Since SST cells receive strong excitation from L2/3 PCs (Adesnik et al., 2012; Hakim et al., 2018; Kapfer et al., 2007; Karnani et al., 2016), this difference in tuning was surprising.

This raised the possibility that strong inhibition from L2/3 VIP cells to SST cells counteracts PC→SST excitation to explain SST cells’ remarkable insensitivity to low contrast. VIP cells, which participate in a powerful inhibitory feedback loop with SST cells (Pfeffer et al., 2013), exhibit the strongest surround suppression of all cell types at high contrast (Dipoppa et al., 2018) and selectively respond to low contrast at large size (Millman et al., 2020). Whether and how VIP contrast responses vary as a function of stimulus size, however, is not known. In opposition to SST cells, VIP cells as a population were selectively responsive at small size and low contrast, and strongly suppressed at large size and high contrast (fig. 1p). Further, VIP cells on average showed non-monotonic contrast tuning, with the preferred contrast decreasing with increasing size (fig. 1q). At a single neuron level, this population average reflected a mixture of behaviors markedly different from other cell types’ (fig. S3a-c), from strict contrast suppression (fig. S3d-f; note contrast-induced decrease in fluorescence in fig. S3e), to non-monotonic size and contrast tuning (fig. S3g-i). These data demonstrated that the relative strengths of SST vs. VIP cells’ activities demarcated disparate regimes of visually-evoked inhibitory circuit activity – a VIP-dominated regime, at small size and low contrast, and an SST-dominated regime, at large size and high contrast.

We next asked whether the tuning of VIP and SST cells, though significantly different from all other cell types measured (fig. 2a), might still be predictive of PC tuning. Because SST and VIP cell activity were strongly anti-correlated (fig. 2b), we restricted our attention to VIP activity, which we first compared with measures of PC surround suppression. We computed the slope of PC tuning with respect to size (fig. 2c) as a measure of surround suppression at each size and contrast. Negative slope corresponded to surround suppression, and positive slope to facilitation. PC size tuning slope was strongly correlated with VIP cell activity (fig. 2d, R=0.51, p=0.001, Wald test). We next asked whether the PC contrast sensitivity was likewise related to VIP cell activity; thus, we measured the slope of PC tuning with respect to contrast as a function of stimulus in a similar way (fig. 2e). Indeed, PC contrast tuning slope was strongly correlated with VIP cell activity and was nearly zero (which would indicate complete insensitivity to contrast) where VIP activity was weakest (fig. 2f, R=0.44, p=7×10^-3^, Wald test). Thus, in the VIP- dominated regime, PCs sensitively encoded contrast and were surround facilitating, and in the SST-dominated regime, PCs were insensitive to contrast, but strongly surround suppressing.

**Figure 2.**
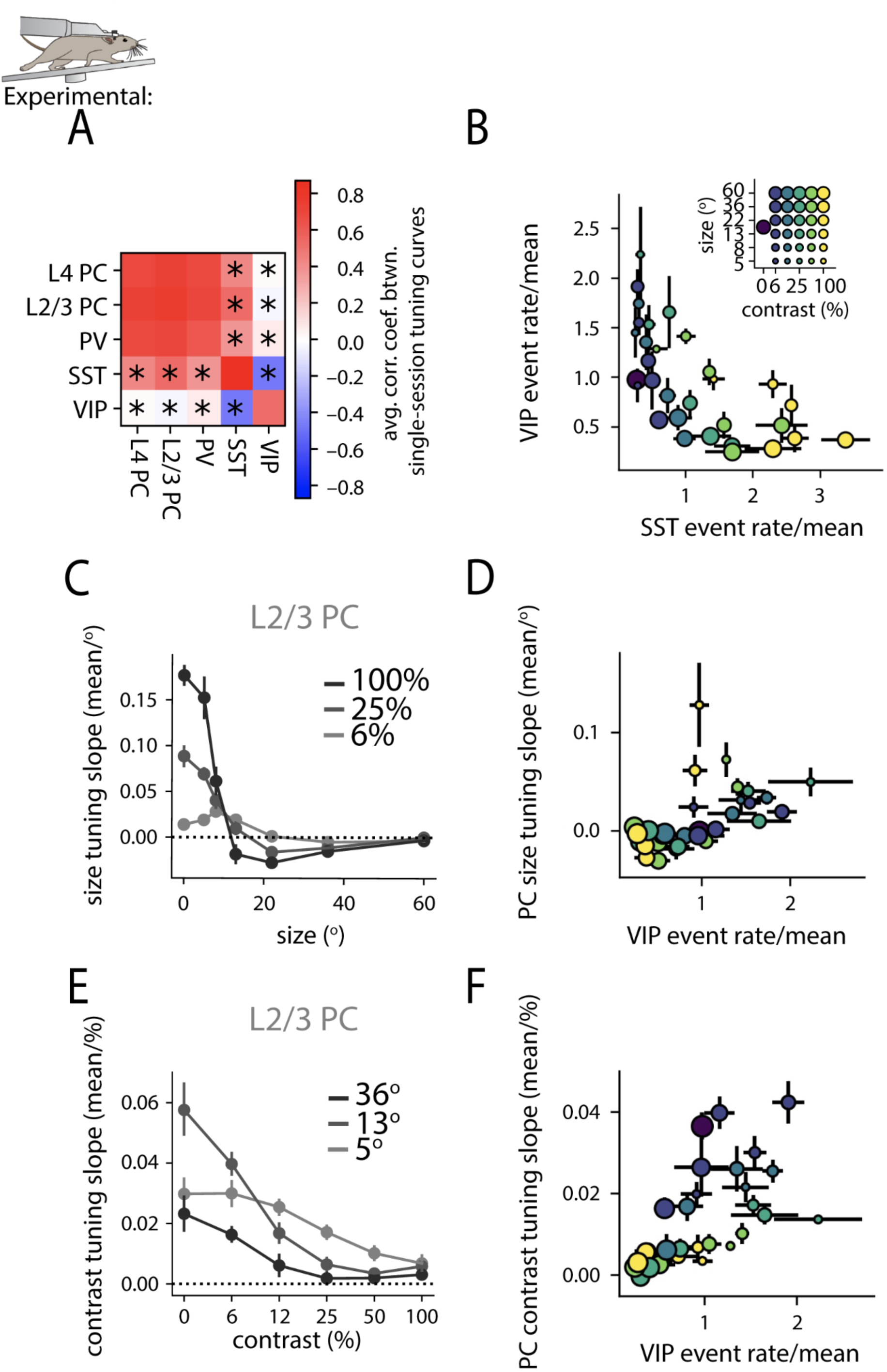
SST and VIP activity outline disparate regimes of PC spatial integration. A) Average Pearson correlation coefficient between population tuning curves estimated from single imaging sessions, either within a cell type (diagonal elements) or between cell types (non-diagonal elements). * across-cell-type correlations for a given pair are significantly less positive than within-cell-type correlations for the cell type corresponding to the row (p < 0.05, Mann-Whitney U-test; *n* = 6 L4 PC, 9 L2/3 PC, 6 L2/3 PV, 5 L2/3 SST, 5 L2/3 VIP imaging sessions). B) Average event rate of VIP cells plotted against the average event rate of SST cells; in this and the following panels, event rates are mean-normalized, or divided by the mean across stimuli and across neurons within each imaging session. Dots indicate stimuli of varying size (dot size) and contrast (dot color—increasing from cold to warm colors) in the model. C) The local slope of PC size tuning curves plotted against stimulus size (see Methods; in units of mean event rate across stimuli and across neurons, per visual degree of size). Where the slope is positive, cells undergo surround facilitation on average, and where the slope is negative, cells experience surround suppression. D) PC size tuning slope plotted against the difference of VIP event rate (R=0.51, p=0.001, Wald test). E) Same as C), but for the local slope of PC contrast tuning curves (in units of mean event rate per percent contrast). F) Same as D), but with PC contrast tuning slope rather than size tuning slope (R=0.44, p=7×10^-3^, Wald test). Error bars indicate bootstrap SEM equivalent.

### A recurrent network model explains size and contrast tuning of the four cell-type circuit

In our physiological data, VIP and SST activity outlined disparate regimes of PC surround suppression and contrast sensitivity. We further hypothesized that VIP-SST competition gave rise to opposing VIP and SST tuning, which in turn supported flexible cooperation and competition among PCs to shape PC tuning. To test these mechanistic hypotheses, we developed a data-driven recurrent network model (Dipoppa et al., 2018; Keller et al., 2020). Using the data on “center” neurons presented above along with the responses of “surround” neurons more than 10° of retinotopic space from the stimulus representation center (fig. S4), we fit a stabilized supralinear network model (Ahmadian et al., 2013; Rubin et al., 2015) (see Methods). In particular, each L2/3 unit had a supralinear (expansive) relationship between input current and output firing rate, so that its gain (its change in output for a given small change in input) increased as its input current increased (Hansel & Van Vreeswijk, 2002; Miller & Troyer, 2002). The model simulated two spatial domains (center and surround) each containing five cell types (fig. 3a): L4 PCs, which served as static feedforward inputs to the network for each stimulus condition, and the four L2/3 cell types. Cell types were modeled as homogeneous populations. In the model, each L2/3 cell type received a linear combination of a stimulus-independent bias current (controlling input current in the absence of inputs from other cell types), stimulus-dependent input from L4 PCs, and inputs from each other population. Connections between two cell types in different spatial domains were given by the within-domain weight, multiplied by a cross-space factor dependent on the postsynaptic cell type. These synaptic inputs were transformed through an expansive nonlinearity to determine the cell type’s firing rate. Along with synaptic currents from within the modeled network, we allowed the fitting procedure to optimize small cell-type- and stimulus-specific “residual” currents not accounted for by the activity of other cell types in the network, which could be interpreted as stimulus-tuned inputs coming from outside L2/3. These were constrained to be small relative to currents coming from the modeled cell types ((std. dev. of residual currents)/(std. dev. of currents) = 13.6% +/- 1.7%, mean +/- std. dev. across model fits).

**Figure 3.**
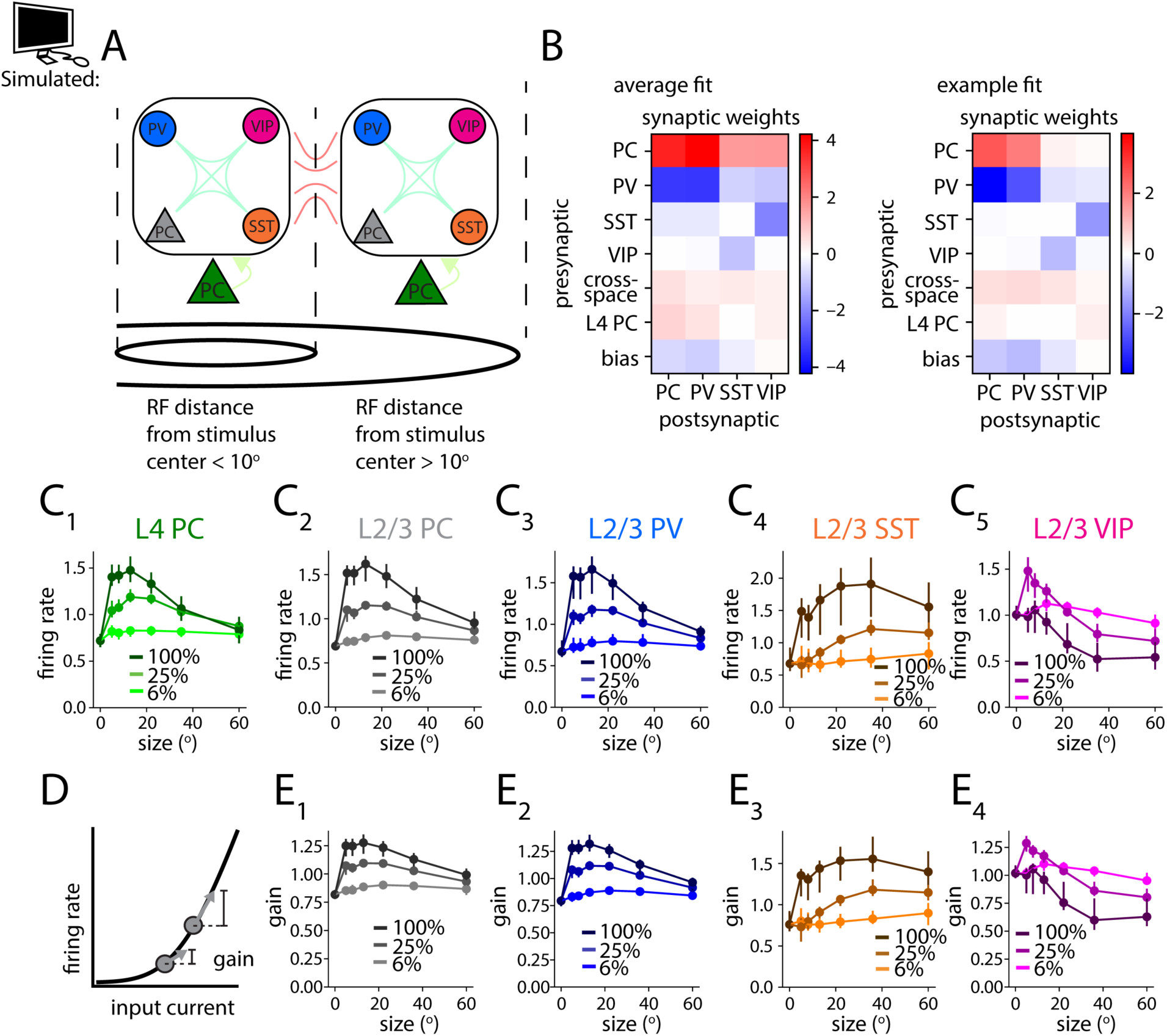
A network model explains the size and contrast tuning properties of layer 2/3 cell types. A) Schematic of the model: the rate-based network model consisted of five cell types, spread across two spatial domains. To fit the calcium imaging data, we optimized a set of weights describing connection strengths among these cell types. Connections between two cell types across spatial domains were attenuated by a cross-space weight that was allowed to vary based on the postsynaptic cell type. B) Synaptic weight matrix between cell types within one spatial position, averaged across *n* = 94 model fits, and of a representative example fit. C) Size tuning of all five modeled cell types, by contrast. D) Schematic of expansive nonlinearity characterizing modeled L2/3 neuron responses as a function of input current. As input current increases, both firing rate and the slope of firing rate with respect to input current (gain) increase. E) Gain of all four modeled L2/3 cell types vs. size, by contrast. Error bars indicate 16^th^ to 84^th^ percentile of successful model fits.

Importantly, the network was constrained to be in an inhibitory stabilized regime across all stimulus conditions, meaning that dynamic changes in inhibition were required to prevent the subnetwork of L2/3 PCs from becoming unstable. Mouse V1 has been shown to operate in this regime (Sanzeni et al., 2020). We analyzed the behavior of the best fitting models, starting from many instantiations of randomly initialized model parameters (see Methods) to identify connection strengths and other model parameters robustly required to explain the calcium imaging data (fig. 3b, fig. S5). Notably, many of the connection strengths in these fits corresponded well to experimentally measured connection strengths from prior studies, including the strong connections among PCs and PV cells, and the strong reciprocal inhibition between SST cells and VIP cells. These models were able to recapitulate key tuning features for each cell type (fig. 3c), including contrast-dependent surround suppression in PCs and PV cells, low contrast sensitivity and large size preference of SST cells, and non-monotonic size and contrast tuning of VIP cells. Importantly, because firing rates of L2/3 cell types were modeled to be supralinear with respect to their input currents, (fig. 3d), each cell type’s sensitivity to changes in input (gain) was highest when it was most active (fig. 3e). Thus, polysynaptic effects mediated by a particular cell type would increase in strength with that cell type’s activity, a key mechanism for network tuning to shape cooperation and competition.

### Determinants of VIP-dominated vs. SST-dominated regimes

Using this network model, we sought to understand the mechanisms underlying the correlations between VIP activity and PC size and contrast tuning slope which we uncovered in the data. Our first goal was to understand how increasing PC activity gave rise to disparate VIP- dominated and SST-dominated regimes. First, we asked why at low contrast, VIP cells were highly active, and increasing PC activity drove VIP cells but not SST cells. We reasoned that high spontaneous drive to VIP cells (perhaps from cholinergic inputs (Fu et al., 2014)), or equivalently, a lower distance from rest to threshold for VIP cells, could explain both phenomena. If spontaneous drive to VIP cells (bias currents in the model) outweighed spontaneous drive to SST cells, then VIP cells could dominate in response to low levels of PC activity. In fact, in every successful model fit (94/94), the bias current to VIP cells was more positive than the bias current to SST cells (fig. 4a).

**Figure 4.**
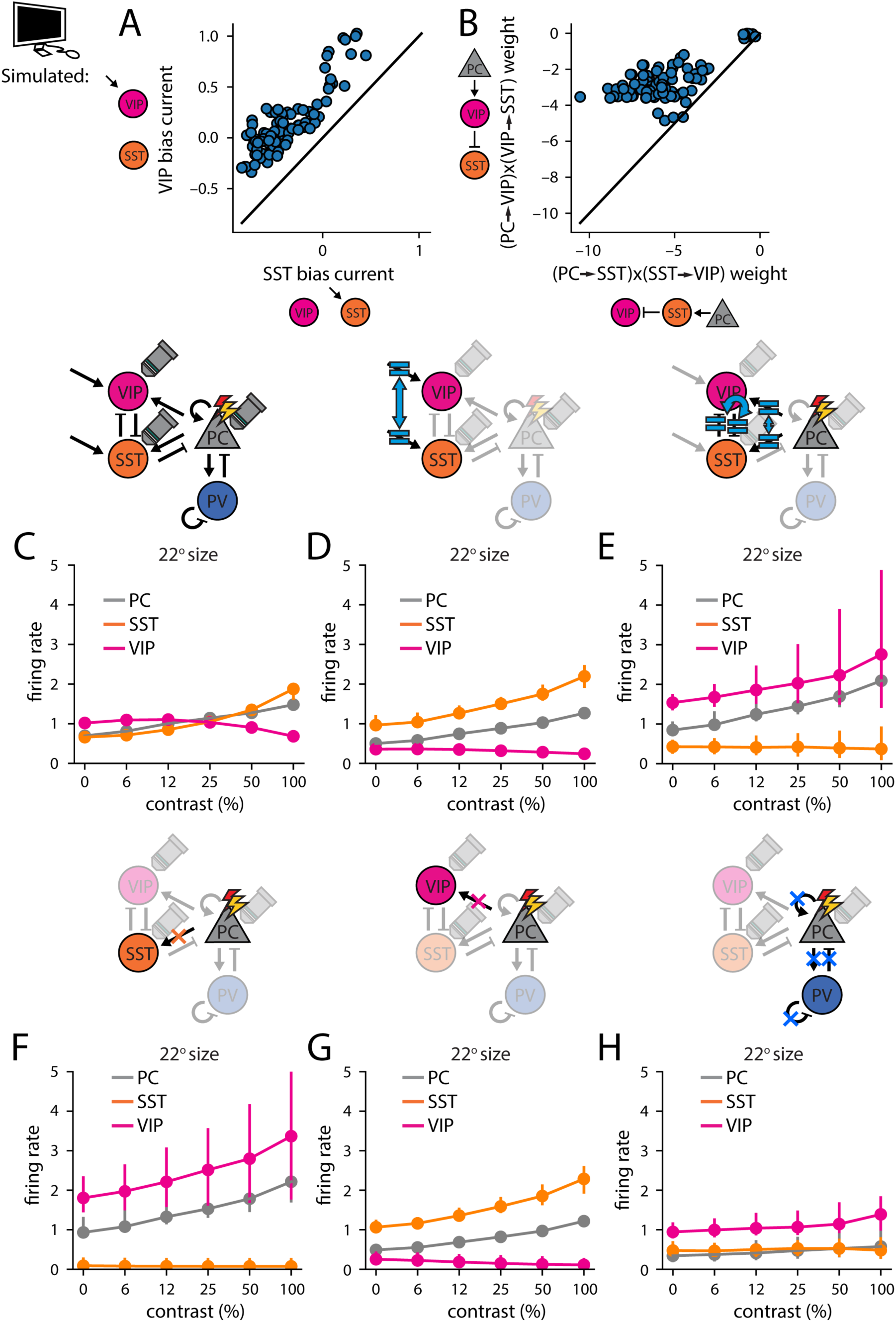
Determinants of VIP-dominated vs. SST-dominated regimes. A) In all models that fit the data well, the bias current, which in the recurrent network model controls firing rate in the absence of activity in the rest of the network, was greater in VIP cells than in SST cells (*n* = 94/94 model fits). B) In 93/94 models that fit the data well, the product of PC→VIP and VIP⊣SST connection weights was smaller than the product of PC→SST and SST⊣VIP connection weights, summing within- and across-spatial-domain weights in each case. C) The firing rate of modeled center PCs (gray), SSTs (orange), and VIPs (magenta) vs. contrast at 22° size (*n* = 89 model fits; 5 model fits showed divergent responses of at least one cell type to at least one of the manipulations in the following panels). D) Same as C), but setting VIP bias current equal to SST bias current. E) Same as C), but setting PC→SST connection weights to equal PC→VIP connection weights, and SST⊣VIP connection weights to equal VIP⊣SST connection weights (all summing within- and across-spatial-domain weights). F-H) Same as C), but deleting all PC→SST, PC→VIP, or all PC→PC, PC→PV, PV⊣PC, and PV⊣PV connections, respectively. In D-H, the stated manipulations were performed both in center and surround circuits. Error bars indicate 16^th^ to 84^th^ percentile of successful model fits.

Next, we aimed to understand what shifted the network to an SST-dominated regime in response to high contrast gratings. We reasoned that when the PC network was highly active, VIP-SST competition would favor whichever of the two cell types could more effectively convert increasing PC activity to increasing disynaptic inhibition of the other cell type. The strength of such disynaptic effects in the model depended on the product of the two synaptic weights as well as the gains (*i.e.*, activity levels) of the cell types involved. Thus, if the product of PC→SST weights and SST⊣VIP weights were larger than the product of PC→VIP weights and VIP⊣SST weights, we hypothesized that growing levels of PC activity (and thus PC gain) could allow SST cells to overcome VIP cells’ initial advantage in spontaneous input current and ultimately outcompete them. Indeed, in ∼99% of successful model fits (93/94), the product of PC→SST⊣VIP weights was stronger than the product of PC→VIP⊣SST weights (fig. 4b). Together with the previous finding, this suggested that differential bias current supported the VIP- dominated regime at low contrast, while SST cells mediated powerful disynaptic inhibition of VIP cells to support the SST-dominated regime at high contrast. These differences in SST and VIP connectivity in the model offered a possible explanation for how increasing feedforward drive to PCs could shift the network from a VIP-dominated to an SST-dominated regime with increasing contrast.

We next tested the hypothesis that these differences in connectivity were necessary to explain the shift from a VIP-dominated to an SST-dominated regime. We first examined modeled PC, SST, and VIP responses as a function of contrast, at 22° size, which showed a shift from VIP-dominated to SST-dominated at the highest contrasts (fig. 4c). Keeping L4 PC activity constant, we then observed how L2/3 network activity changed as a function of the model parameters we identified. Reducing the VIP bias current to be equal to the SST bias current allowed SST cells to dominate across contrasts (fig. 4d), while setting the PC→SST and SST⊣VIP connection weights to be equal to the PC→VIP and VIP⊣SST connection weights, respectively (fig. 4e), allowed VIP cells to dominate across contrasts. These results supported the idea that a large stimulus-independent bias current to VIP cells competed with strong stimulus-dependent disynaptic inhibition via the PC→SST⊣VIP circuit to support opposing regimes of VIP and SST activity depending on contrast.

We hypothesized that excitatory connections within L2/3, in particular PC→SST and PC→VIP, might flexibly support SST or VIP activity depending on stimulus. To test this idea, we next specifically deleted these connections. In the absence of PC→SST connections, VIP cells dominated across contrasts (fig. 4f), while in the absence of PC→VIP connections, SST cells dominated across contrasts (fig. 4g). Next, we wondered whether recurrent connections between PCs and PV cells, the strongest in the network fits, might play a role in amplifying the degree of competition between SST and VIP. In fact, deleting all connections among PCs and PV cells (PC→PC, PC→PV, PV⊣PC, and PV⊣PV) caused PC activity to vary only shallowly with contrast, resulting in VIP cells dominating across contrasts (fig. 4h). We repeated these manipulations of the network, replacing increasing contrast with increasing current injection to PCs starting from the 0% contrast condition, and observed similar results (fig. S8), showing that these connections were likewise sufficient to explain the shift from a VIP-dominated to an SST-dominated regime with increasing PC activity. Thus, the greater spontaneous drive to VIP cells and the stronger disynaptic inhibition onto VIP, along with excitatory connections within L2/3, appeared to explain the shift from a VIP-dominated to an SST-dominated regime with increasing contrast.

### Determinants of PC cooperation vs. competition

We hypothesized that contrast-dependent PC cooperation and competition could shape contrast-dependent surround suppression. To better understand how PC cooperation and competition changed going from the VIP- to the SST-dominated regime, we next looked at the effects of injecting current in either center PCs or surround PCs alone, while varying visual stimulus contrast, again at 22° size. We hypothesized that, in the VIP-dominated regime, the increased gain of VIP cells would support positive PC→VIP⊣SST⊣PC feedback over negative PC→SST⊣PC feedback, and that the opposite would be true in the SST-dominated regime. As a first test, we injected excitatory current in center or surround PCs, and measured responses of center PCs, VIP cells, and SST cells, starting from the steady state for visual stimuli of varying contrasts (fig. 5a—see Methods). Consistent with our core hypothesis, as contrast increased, moving from the VIP-dominated to the SST-dominated regime, current to center PCs increasingly drove center SST cells, suppressed center VIP cells, and in turn more weakly drove center PCs (fig. 5a). Similarly, excitatory current injection to surround PCs facilitated center PCs on average at low contrast, corresponding to cooperation across space, but suppressed center PCs on average at high contrast, corresponding to competition (fig. 5b). Thus, as VIP activity decreased with contrast, additional drive to PCs increasingly drove SST and suppressed VIP activity (fig. 5a,b), and correspondingly, PC spatial interactions changed from cooperative to competitive (fig. 5c, R=0.17, p=4×10^-24^, Wald test). Within the model, the cooperation and competition of L2/3 PCs across space was correlated with the slopes of PC size and contrast tuning (fig. 5d,e, R=0.11, p=2×10^-11^; R=0.35, p=6×10^-98^, Wald test), in keeping with the idea that cooperation and competition of PCs across space could shape their stimulus tuning.

**Figure 5.**
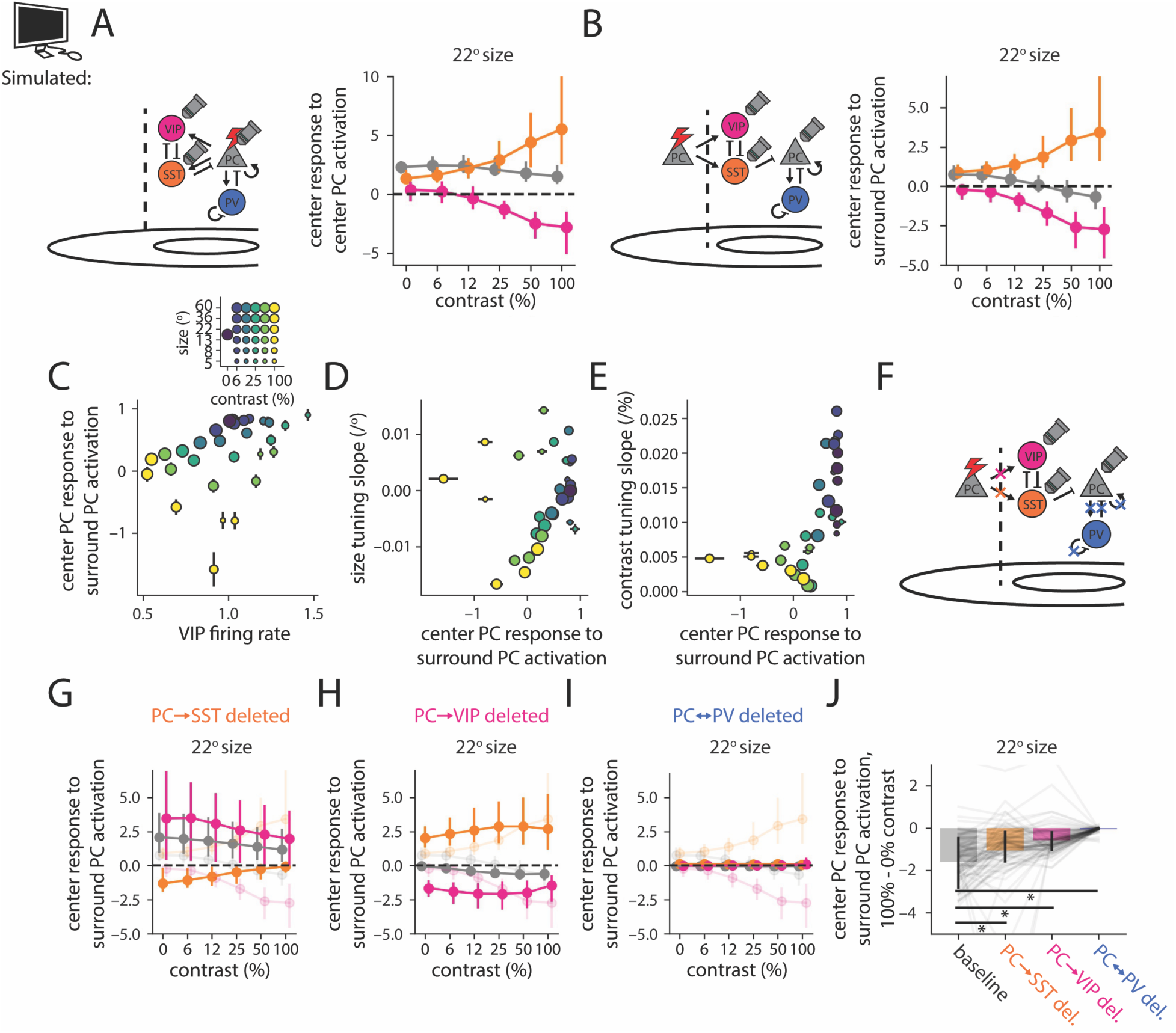
Determinants of modeled VIP-SST competition control PC-PC cooperation and competition. A) Left: schematic showing perturbations to center L2/3 PCs in the model, and recording of responses in PCs, SST cells, and VIP cells. Right: Plot of the response of PCs (gray), SST cells (orange) and VIP cells (magenta) in the center of the visual field to the injection of infinitesimal excitatory current into PCs in the center domain, starting from the steady state for visual stimuli of varying contrasts and 22° size (*n* = 89 model fits; excluding 5 fits that showed divergent response to at least one connection deletion manipulation). B) Same as A), but for current injected into L2/3 PCs in the surround domain. C) Response of center PCs to infinitesimal current injection into surround PCs vs. VIP firing rate for varying size and contrast (see legend; R=0.17, p=4×10^-24^, Wald test). D) Slope of center PC size tuning curve vs. the response of center PCs to infinitesimal current injection into surround PCs for varying size and contrast (R=0.11, p=2×10^-11^, Wald test). E) Same as D), but for slope of center PC contrast tuning curve (R=0.35, p=6×10^-98^, Wald test). F) Schematic: same perturbation as in B), but after deleting specified connections and allowing the network to settle to a new steady state. G) Same as B), but after deleting all PC→SST connections. H,I) Same as B), but after deleting PC→VIP connections (H), or all PC→PC, PC→PV, PV⊣PC, and PV⊣PV connections (I). In G-I, transparent lines show baseline responses from B) for ease of comparison. J) Difference between response of center PCs to infinitesimal current injection in surround PCs, between 100% and 0% contrast at 22° size, for the conditions shown in B,G,H,I), respectively (difference significantly less for the baseline condition; p=1×10^-4^, p=4×10^-16^, p=1×10^-13^, respectively, for comparisons labeled with *, Wilcoxon signed-rank test.). Error bars indicate 16th to 84th percentile of successful model fits. In all panels, Y response to infinitesimal X activation corresponds to the derivative, dY/dX.

We hypothesized that an interplay between PC→VIP⊣SST⊣PC and PC→SST⊣PC feedback was necessary for the contrast-dependent shift from PC cooperation to competition we observed. To test this, we deleted individual L2/3 excitatory connections within the model, as before, to selectively disrupt either the PC→SST⊣PC or the PC→VIP⊣SST⊣PC feedback loop. To test the importance of PC→SST⊣PC feedback, we deleted PC→SST connections, both within and across spatial domains, allowed the rest of the network to settle to a new steady state, and then repeated the activation of surround L2/3 PCs as before (fig. 5f). Deleting this connection promoted cooperation at all contrasts and weakened the contrast dependence of this competition overall (fig. 5g). Deleting PC→VIP connections in a similar way promoted competition across all contrasts and weakened the contrast dependence of this competition overall (fig. 5h). This demonstrated that in the model, a balance of negative PC→SST⊣PC feedback and positive PC→VIP⊣SST⊣PC feedback was necessary for the contrast-dependent shift from PC cooperation to competition between PCs across space. Likewise, deleting recurrent connections among PCs and PV cells, which we observed to disrupt the contrast dependence of VIP-SST competition, also disrupted the contrast dependence of PC competition across space in the model (fig. 5i). Thus, the feedback loops supporting VIP-SST competition were also necessary for the contrast-driven shift from PC cooperation to competition across space in the model (fig. 5j).

### Formulation of model predictions

These manipulations within our computational model implicated several network mechanisms in mediating a shift from PC cooperation to competition with increasing contrast. We reasoned that optogenetically perturbing VIP cells *in vivo* would allow us to empirically probe the biological system for signatures of these mechanisms. First, we used our computational model to predict the impact of injecting either excitatory or inhibitory current into VIP cells. In the absence of a visual stimulus, VIP perturbations strongly altered network activity, but showed a clear asymmetry: excitatory current injections caused robust facilitation of VIP cells and PCs and suppression of SST cells, while inhibitory current injections caused weaker (and opposite) effects (fig. 6a_1_). In the presence of a visual stimulus (e.g. at 22° size, 100% contrast), network activity showed a strongly sigmoidal relationship with injected VIP current: PCs were highly sensitive to currents within a narrow range, but insensitive elsewhere (fig. 6a_2_). However, deleting the excitatory connections within L2/3 necessary for VIP-SST competition (PC→SST, PC→VIP, or PC-PV, as above) abolished the asymmetry between inhibitory and excitatory current injection, as well as the sigmoidal behavior (fig. 6b-d). Thus, the computational model predicted an asymmetry between the effects of inhibitory and excitatory current to VIP cells, supported by the same feedback loops implicated in VIP-SST competition.

**Figure 6.**
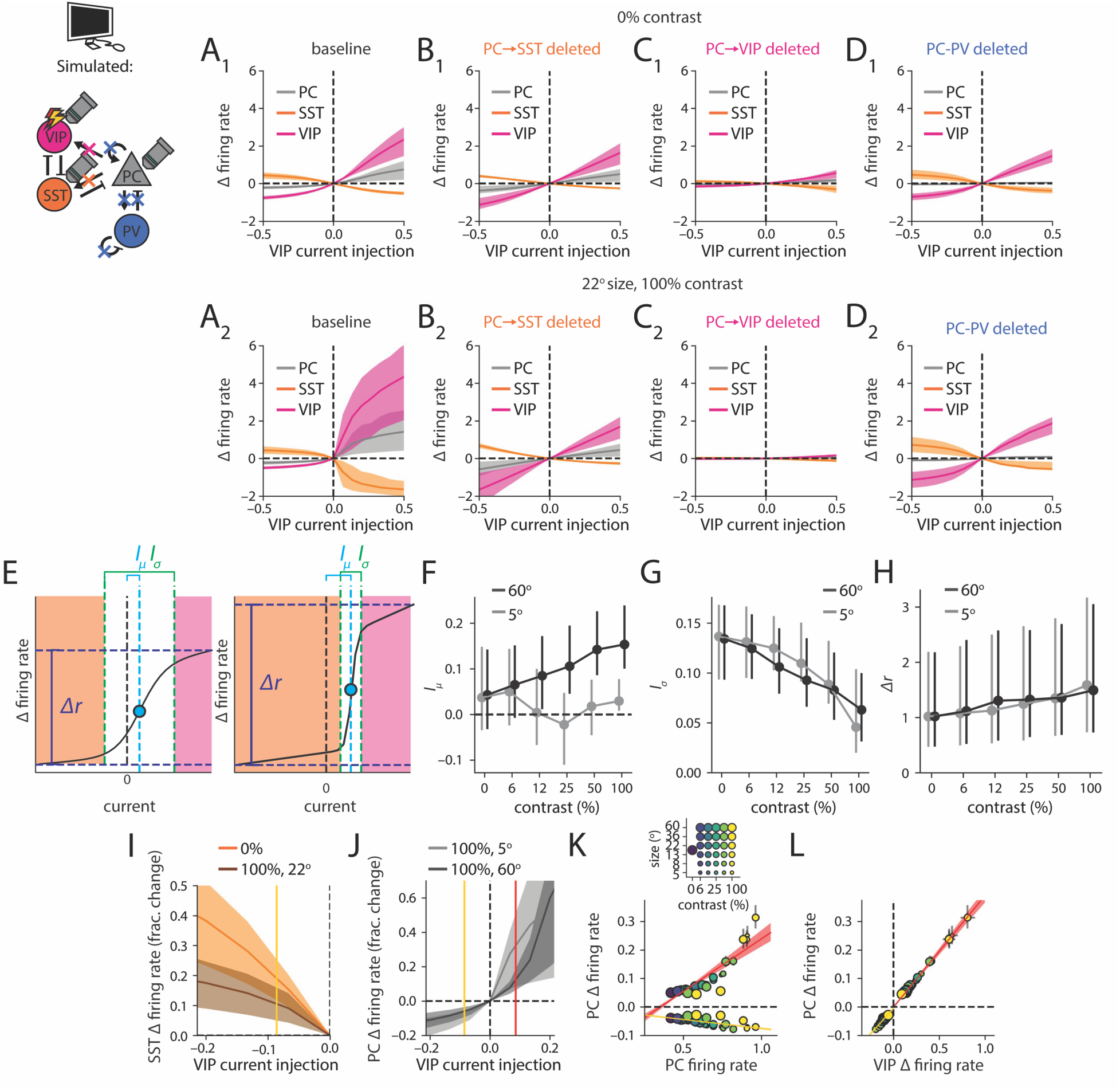
Sign and magnitude of modeled VIP current injection qualitatively shape the effect of VIP cells on the network. A_1_) Change in firing rate vs. current injection to VIP cells for PCs, SST cells, and VIP cells, in the 0% contrast condition, with no connections deleted (*n* = 89 model fits; excluding 5 fits that showed divergent response to at least one connection deletion manipulation). A_2_) Same as A_1_), but in the 22° size, 100% contrast condition. B-D) Same as A), but with PC→SST, PC→VIP, and PC-PV connections deleted, respectively. E) Effects of VIP perturbation on PCs are parameterized as sigmoids with six parameters: initial slope and offset, final slope and offset, transition current *I_μ_* (midpoint of sigmoidal transition between initial and final asymptote), and transition scale *I_σ_* (width of sigmoidal transition between initial and final asymptote) (see Methods). We focus on the last two parameters, as well as the amplitude *Δr*, defined as the difference between firing for the most negative and most positive currents, respectively. To the left, a sigmoid fit is shown with small *I_μ_*, large *I_σ_*, and small *Δr*, and to the right, a sigmoid fit with larger *I_μ_*, smaller *I_σ_*, and larger *Δr*. Orange denotes the SST-dominated, and magenta denotes the VIP-dominated regime. Note that the network in the absence of injected VIP current is in the “transition regime” (shown in white) if |*I_μ_*|<*I_σ_*. F) *I_μ_* is plotted vs. contrast for 5° and 60° size. G) Same as F), but for *I_σ_*. Note that the network in the absence of injected VIP current is in the transition regime at low contrast, but not at large size and high contrast. H) Same as F), but for *Δr*. I) Overlay of SST Δ firing rate from A_1_) and A_2_), expressed as fractional change (i.e., change divided by original value) from firing rate in the absence of VIP current, vs. inhibitory injected VIP current. Yellow vertical line indicates “intermediate” current injection magnitude, defined as the mean |*I_μ_*| across all sizes at 100% contrast. J) Overlay of PC Δ firing rate, expressed as a fractional change, vs. injected VIP current for 5° and 60° size, respectively, at 100% contrast. Note that due to varying values of *I_μ_* for these conditions, intermediate excitatory current injection (red) is sufficient to drive the network into the transition regime at 5° but not at 60° size. K) Scatter plot of PC Δ firing rate in response to intermediate current injection shown in I) and J), vs. PC firing rate in the absence of current injection. Points with y-value above zero correspond to excitatory current injection (red line, linear fit with shaded bootstrapped confidence intervals) and with y-value below zero, inhibitory current injection (yellow line, linear fit). Inset: legend for the dots shown in this and the subsequent figure. L) Same as K), for PC Δ firing rate vs. VIP Δ firing rate. Error bars indicate 16th to 84th percentile of successful model fits.

To quantify the response of the network as a function of injected VIP current, we fit sigmoid functions to PC activity for each stimulus condition (fig. 6e). We defined the regions where PCs were insensitive to VIP current to be the “fully SST-dominated” and “fully VIP- dominated” regimes, respectively, and the region where PCs were sensitive to be a “transition” regime. We defined the “transition current” as the current needed to shift the network to the midpoint of the transition regime. The transition current was positive for most stimuli, indicating that the unperturbed network was closer to the fully SST-dominated than to the fully VIP-dominated regime (fig. 6f). At large size, the transition current increased with increasing contrast, consistent with an increasingly SST-dominated regime for increasingly large size and high contrast; the transition current remained close to zero at small size (fig. 6f). With increasing contrast, the transition regime of the sigmoid narrowed (fig. 6g; indicating a more switch-like transition between the fully VIP-dominated and fully SST-dominated regimes). Whereas at low contrast, the unperturbed network was in the transition regime (see fig. 6e,f,g), at large size and high contrast, it was well outside it. Further, with increasing contrast, the overall effect of current injection increased (fig. 6h).

Through its prediction that the majority of the effect of VIP perturbations would be limited to currents around the transition regime, our model generated three key predictions. First, VIP inhibitory current injections would have the greatest effect on SST activity at low contrast, where the network was within the transition regime in the absence of VIP current (fig. 6i). Second, the effects of excitatory and inhibitory current injections would be asymmetric, since the unperturbed network was not far from the fully SST-dominated regime for most stimuli, and transition currents were positive and stimulus-dependent. Therefore, unlike VIP inhibitory currents, excitatory current injections which were intermediate in strength should be sufficient to push the network into the transition regime specifically at small size or low contrast, but not at large size and high contrast, where the unperturbed network was far from the transition regime (fig. 6j). This would result in a larger and more strongly contrast-dependent effect of VIP excitatory current injection compared with VIP inhibitory current injection. Third, the difference in PC activity between the SST- and VIP-dominated regimes was largest, and the transition steepest, at high contrast, when PCs were highly active in the unperturbed network. Thus, the effects of VIP excitatory and inhibitory current injections on PCs should in general increase in strength as unperturbed PC activity increased (fig. 6k). A strong linear (additive and multiplicative) relationship would exist between unperturbed PC activity and effect of perturbations. Because PC and VIP activity tended to be tightly coupled, the same would be true of effects on VIP cells as well (fig. 6l). Note that these predictions are not necessarily obvious from the simple fact of VIP-SST competition. For example, (Millman et al., 2020), based on VIP cells being tuned for low contrast, predicted that the effects of VIP perturbations on PCs would be much larger at low contrast than at high contrast. Our model predicted that the effect of VIP perturbations would in general increase with increasing contrast, particularly at smaller sizes.

### Experimental validation of recurrent network model predictions

As explained above, our network model made specific predictions for the impacts of excitatory or inhibitory current injection to VIP cells. We set out to test these predictions with new optogenetic experiments; importantly no optogenetic data were used to fit the model. First, we analyzed more closely within the model the effects of VIP perturbation in a regime of intermediate current magnitudes, such that VIP excitatory current injection was sufficient to push the network into the fully VIP-dominated regime only for some stimuli, as described above (see Methods). The modeling results above might not be directly comparable with experimental results, as experimentally silencing or activating VIP cells might affect other inputs to the network, beyond the recurrently connected populations modeled. Because our network model took L4 PC activity as feedforward input, it was also necessary to measure the direct effects of bidirectional VIP perturbations on L4 PCs and incorporate those effects into the model (see fig. S9a,b and Methods). To simulate the effects of optogenetically silencing VIP cells, we then simulated inhibitory current injections to VIP cells, taking as given the empirically measured effects on L4 PCs. Residual currents from outside the modeled cell types were assumed to remain constant.

Given that positive PC→VIP⊣SST⊣PC feedback was predicted to shape network activity selectively at low contrast, we predicted that silencing VIP cells would strongly disinhibit SST cells in response to these stimuli. Indeed, in simulation, silencing VIP cells increased SST responses most strongly at low contrast (fig. 7a,b,c,d,e). As a result, the contrast sensitivity index of SST cells significantly increased in response to simulated VIP silencing (fig. 7f, p < 10^-10^, Wilcoxon signed-rank test). To test this empirically, we optogenetically silenced VIP cells using Cre-dependent eNpHR3.0, while simultaneously imaging VIP cells, SST cells, and PCs (see Methods and fig. S6). We first confirmed the efficacy of optogenetic silencing; red illumination through the microscope robustly suppressed eNpHR3.0-expressing VIP cell activity (fig. S7). Consistent with model predictions, experimentally silencing VIP cells potently facilitated SST cells across all sizes and contrasts (fig. 7g), but preferentially facilitated SST cells at low contrast (fig. 7h), and only more weakly at high contrast (fig. 7i; however, note that experimentally, facilitation of SST cells at high contrast was stronger than predicted by the model). Thus, across sizes (fig. 7j,k), silencing VIP cells significantly enhanced the sensitivity of SST cells to contrast (fig. 7f,l, p < 0.05, Wilcoxon signed-rank test). This demonstrated that VIP cells shaped the contrast sensitivity of SST cells by strongly suppressing their responses at low contrast.

**Figure 7.**
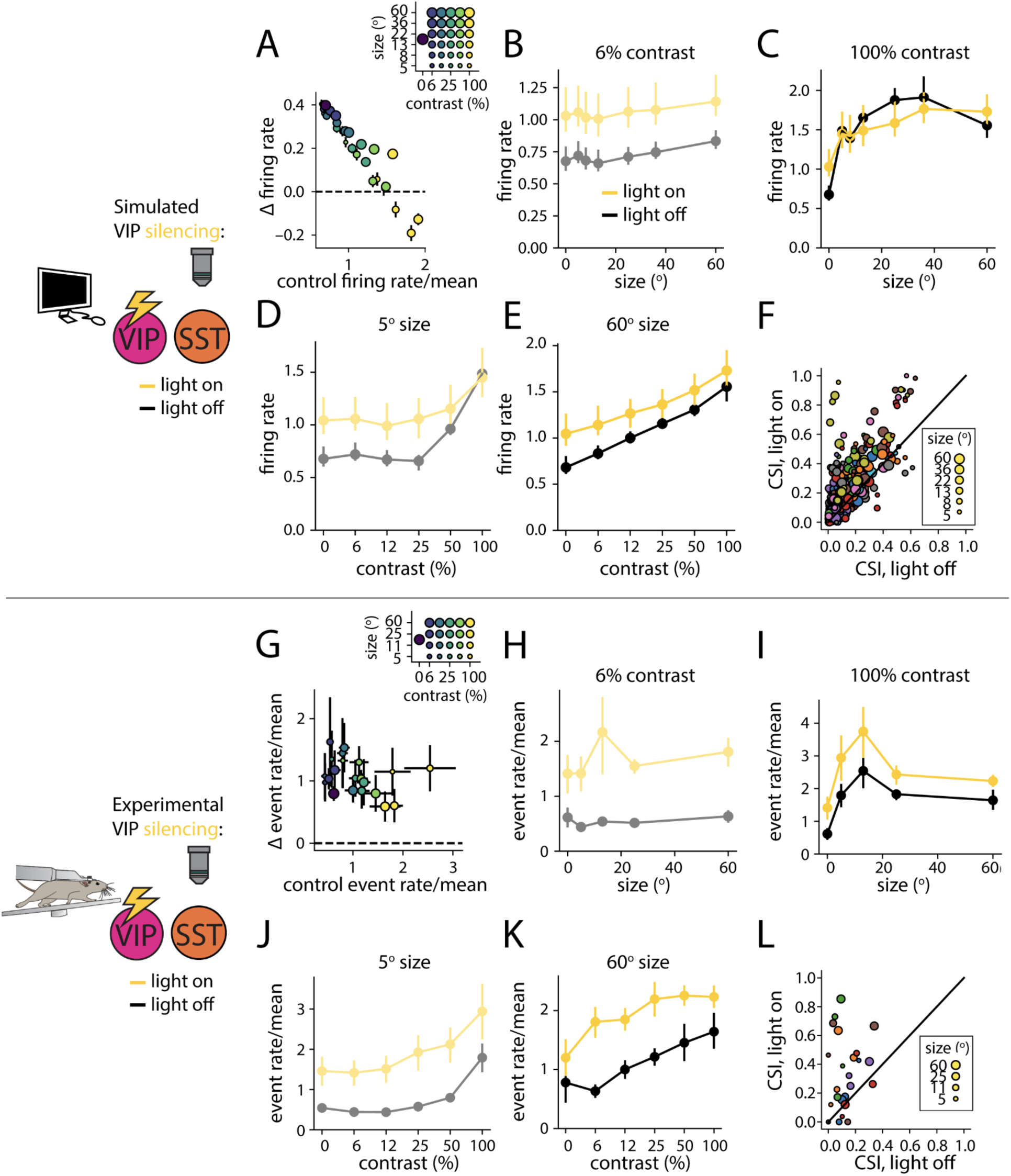
VIP suppression preferentially enhances responses of SST cells, and does so most strongly at low contrast. A) In response to simulated optogenetic silencing of VIP cells, scatter plot of difference between SST event rate with light on and light off vs. SST event rate with light off. Inset: legend for the dots in the scatter plot (*n* = 89 model fits; excluding 5 fits that showed divergent response to at least one connection deletion manipulation). B) Modeled SST event rate as a function of size at 6% contrast in the light off (black) and light on (yellow) condition. C) Same as B) but for 100% contrast. D) SST event rate as a function of contrast at 5° size. E) Same as D) but for 60° size. F) Scatter plot of contrast sensitivity index (CSI, defined as in fig. 1) at varying size in the light on vs. light off condition (significantly higher in the light on condition, p < 10^-10^, Wilcoxon signed-rank test). Size represents stimulus size as in A), and colors indicate individual model fits. G-L) Same as A-F), but for experimental silencing of VIP cells (*n* = 6 imaging sessions; SMI significantly higher in the light on condition, p < 0.05, Wilcoxon signed-rank test). Colors in L) correspond to individual experiments. Error bars in A-F) indicate 16th to 84th percentile of successful model fits, while error bars in G-L) indicate bootstrap SEM equivalent across individual experiments.

We next asked how this effect would propagate through the network polysynaptically to PCs, which only receive weak monosynaptic inhibition from VIP cells on average (Pfeffer et al., 2013). In simulation, we found that VIP silencing significantly suppressed visual responses across most stimuli (fig. 8a,b), but it increased the surround modulation index (SMI) to a similar degree across contrasts (fig. 8c). Thus, SMI slope with respect to contrast showed an inconsistent, but statistically significant, increase (fig. 8d, p < 10^-8^, Wilcoxon signed-rank test; 64/89 model fits increasing). On the other hand, simulated VIP activation produced the largest relative change in PC responses at low contrast (fig. 8e) and at small size and high contrast (fig. 8f), resulting in a significant decrease in SMI slope with respect to contrast (fig. 8g,h, p < 10^-11^, Wilcoxon signed-rank test; 75/89 model fits decreasing). VIP silencing showed that VIP activity was not entirely necessary for, but did weakly enhance, contrast dependence of surround suppression in PCs, while VIP activation showed that adding VIP activity more strongly enhanced contrast dependence of surround suppression in PCs. This result was consistent with the idea that much or all contrast-dependent surround suppression observed under baseline conditions originates in L4, but that contrast-dependent surround suppression could be enhanced by circuitry within L2/3 in a VIP-dependent manner. It was also consistent with the idea that VIP activation would more strongly facilitate PCs at small size or low contrast, where the network was in or near the transition regime, than at large size and high contrast, where the network was in the fully SST-dominated regime.

**Figure 8.**
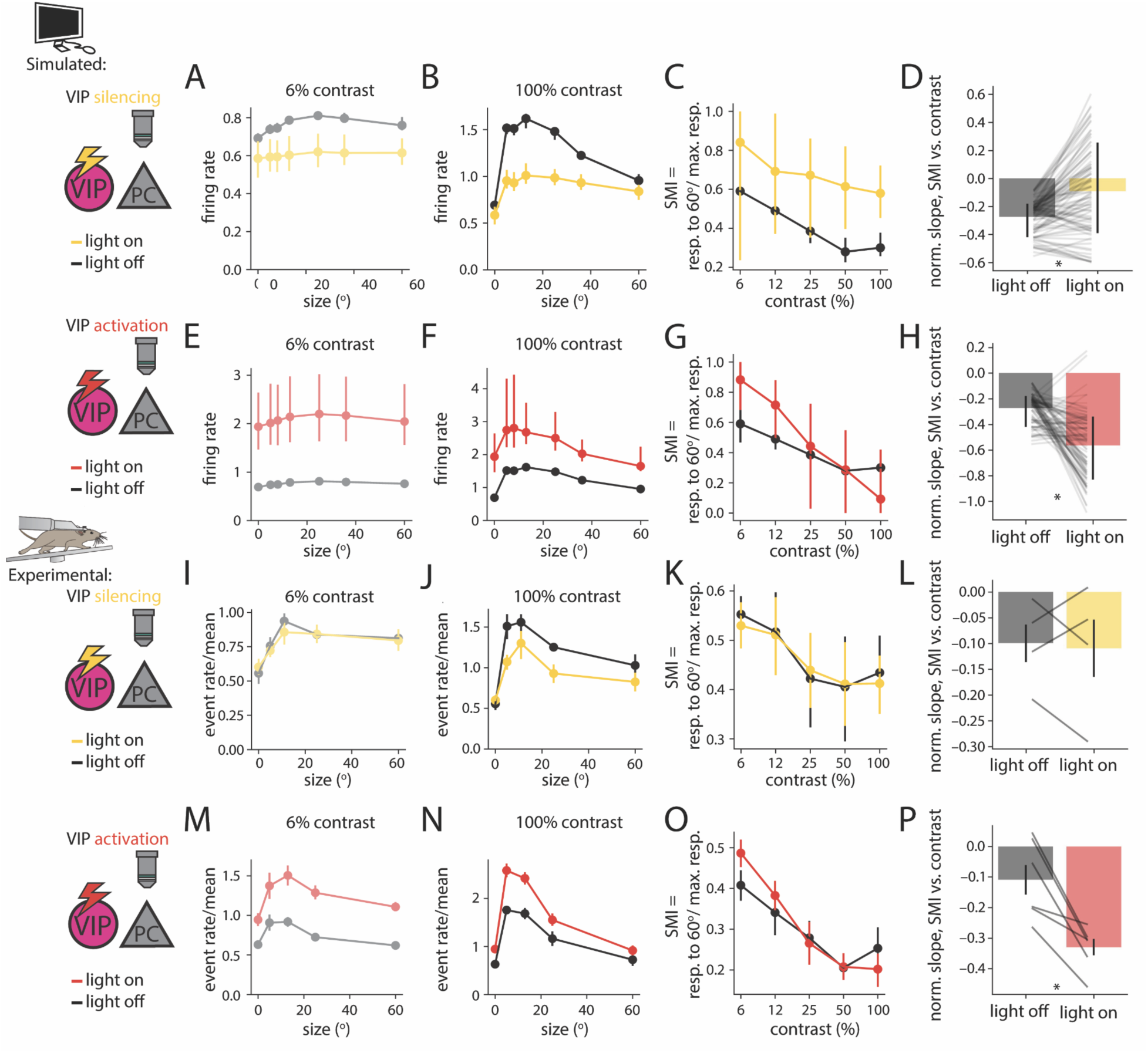
VIP activation strongly enhances contrast dependence of surround suppression. A) Plot of simulated PC event rate versus grating size for 6% contrast in the control, light off condition (light off, black; *n* = 89 model fits; excluding 5 fits that showed divergent response to at least one connection deletion manipulation) and during simulated optogenetic suppression of VIP cells (light on, yellow). B) Same as A) for 100% contrast. C) Surround modulation index (SMI) is plotted as a function of contrast in the light on and light off conditions. D) Change in SMI vs. contrast slope between the light off and light on condition (SMI slope significantly higher in light on condition, p < 10^-8^, Wilcoxon signed-rank test). Each horizontal line represents an individual model fit. Bar indicates mean across model fits, and error bar indicates 16th to 84th percentile across model fits. E-H) Same as A-D), but for simulated optogenetic activation of VIP cells (SMI slope significantly higher in light on condition, p < 10^-11^, Wilcoxon signed-rank test). I-P) Same as A-H), but for experimental silencing and activation and silencing of VIP cells (*n* = 4 imaging sessions, silencing, and *n* = 6 imaging sessions, activation; SMI slope significantly higher in light on condition for VIP activation, p < 0.05, Wilcoxon signed-rank test). Horizontal lines in L,P) correspond to individual imaging sessions, and bars correspond to mean across imaging sessions. Error bars in A-H) indicate 16th to 84th percentile of successful model fits, while error bars in I-P) indicate bootstrap SEM equivalent across individual experiments.

To test these predictions experimentally, we used VIP silencing, as before, but also performed VIP activation. We virally expressed Cre-dependent eNphR3.0-mRuby3 or ChrimsonR-tdTomato in VIP cells in separate mice, and GCaMP6s in all L2/3 neurons using the hSynapsin promoter (see Methods). As with VIP silencing, we confirmed the efficacy of optogenetic activation of VIP cells with ChrimsonR. Red illumination through the microscope objective drove robust activity in ChrimsonR-expressing VIP cells across stimulus conditions (fig. S10a,b), although optogenetic facilitation was somewhat stronger for small and low contrast stimuli (fig. S10c-h). Importantly, in control mice expressing no opsin, the optogenetic stimulation light alone produced no systematic facilitation or suppression of visual responses (fig. S11). This confirmed that the observed effects were due to the optogenetic perturbation *per se*. Next, we analyzed the impact of perturbing VIP cells on the activity of all GCaMP6s-expressing neurons not labeled with a red fluorophore (primarily PCs). In both VIP silencing and VIP activation experiments, PC responses with the optogenetic light off agreed with responses we measured previously using calcium imaging in transgenic GCaMP-expressing mice (fig. S12; VIP silencing, R=0.95, p=3×10^-12^, Wald test; VIP activation, R=0.84, p=3×10^-7^, Wald test).

We then tested our predictions of the effects of VIP perturbations on the contrast dependence of PC surround suppression. We first tested the effects of VIP silencing. Although we found that VIP silencing significantly suppressed PC visual responses at higher contrasts (fig. 8i,j; two-way ANOVA, with factors sensory stimulus, and light; main effect of stimulus, p < 10^-20^; main effect of light, p < 10^-3^; interaction, p > 0.05), it did not significantly affect the PC surround modulation index (SMI) at any contrast (fig. 8k), and thus there was no significant change in PC SMI slope with respect to contrast (fig. 8l). Similarly, there was no significant change in PC contrast sensitivity at any size (fig. S13i-l). To ensure this lack of an effect was not an artifact of calcium imaging, we corroborated these results with extracellular multi-electrode array electrophysiology in a separate cohort of mice, both in the locomoting and the non-locomoting condition, and both in RS units (putative PCs) and FS units (putative PV cells) (fig. S14). This lack of an effect of optogenetic VIP silencing on PC SMI, along with its lack of effect on visual responses at low contrast, disagreed with our model. However, the lack of effect on the contrast dependence of SMI was closer to model predictions, and was consistent with the idea that much of contrast-dependent surround suppression might be independent of VIP activity, and instead largely inherited from feedforward inputs.

Our simulations predicted that activation should produce more substantial effects on PC tuning. Indeed, we found a similar asymmetry in real experiments: VIP activation disproportionately facilitated PCs in response to low contrast stimuli (fig. 8m), but less so for large, high contrast stimuli (fig. 8n; two-way ANOVA, with factors sensory stimulus, and light; main effect of stimulus, p < 10^-59^; main effect of light, p < 10^-34^; interaction, p > 0.05). As a consequence, VIP activation enhanced the contrast dependence of surround suppression, which was evident as an increase in SMI specifically at low contrast (fig. 8o,p) and a decrease in the slope of SMI with respect to contrast. Correspondingly, VIP activation also enhanced the size dependence of contrast sensitivity (fig. S13m-p). Although the magnitude of both measured VIP silencing and activation was weaker than that predicted by the model assuming an “intermediate strength” optogenetic perturbation, the effects on the slope of SMI vs. size were in agreement. Thus, experimental evidence was consistent with our model’s prediction that VIP activation could enhance the existing contrast-dependent surround suppression in L2/3 PCs, but VIP silencing could not fully abolish it.

We next investigated what features of VIP silencing and activation were related to their differential effect on the shape of PC tuning. The network model predicted that much of the impact of these perturbations would be a linear function (with both an additive and a multiplicative term) of the baseline firing rate before current injection (fig. 9a; compare with fig. 6k, which did not include effects on L4 PCs). Across stimuli, higher firing rates in control conditions predicted stronger suppressive effects of VIP silencing (R=-0.55, p < 10^-10^, Wald test) and stronger facilitating effects of VIP activation (R=0.40, p=3×10^-8^, Wald test). The component of perturbations that could be linearly predicted from baseline activity notably did not affect the rank ordering of stimuli: to the extent that the perturbation was entirely linear, the identity of the preferred size and contrast remained the same. As before, simulated VIP activation produced larger effects on both PCs and VIP cells than did VIP silencing, and changes in PC and VIP activity were strongly correlated (fig. 9b). This was in keeping with the finding that the network at baseline was closer to the SST-dominated than to the VIP-dominated regime, and thus that VIP activation could in general result in larger changes to PC activity before it saturated outside the transition regime (fig. 6).

**Figure 9.**
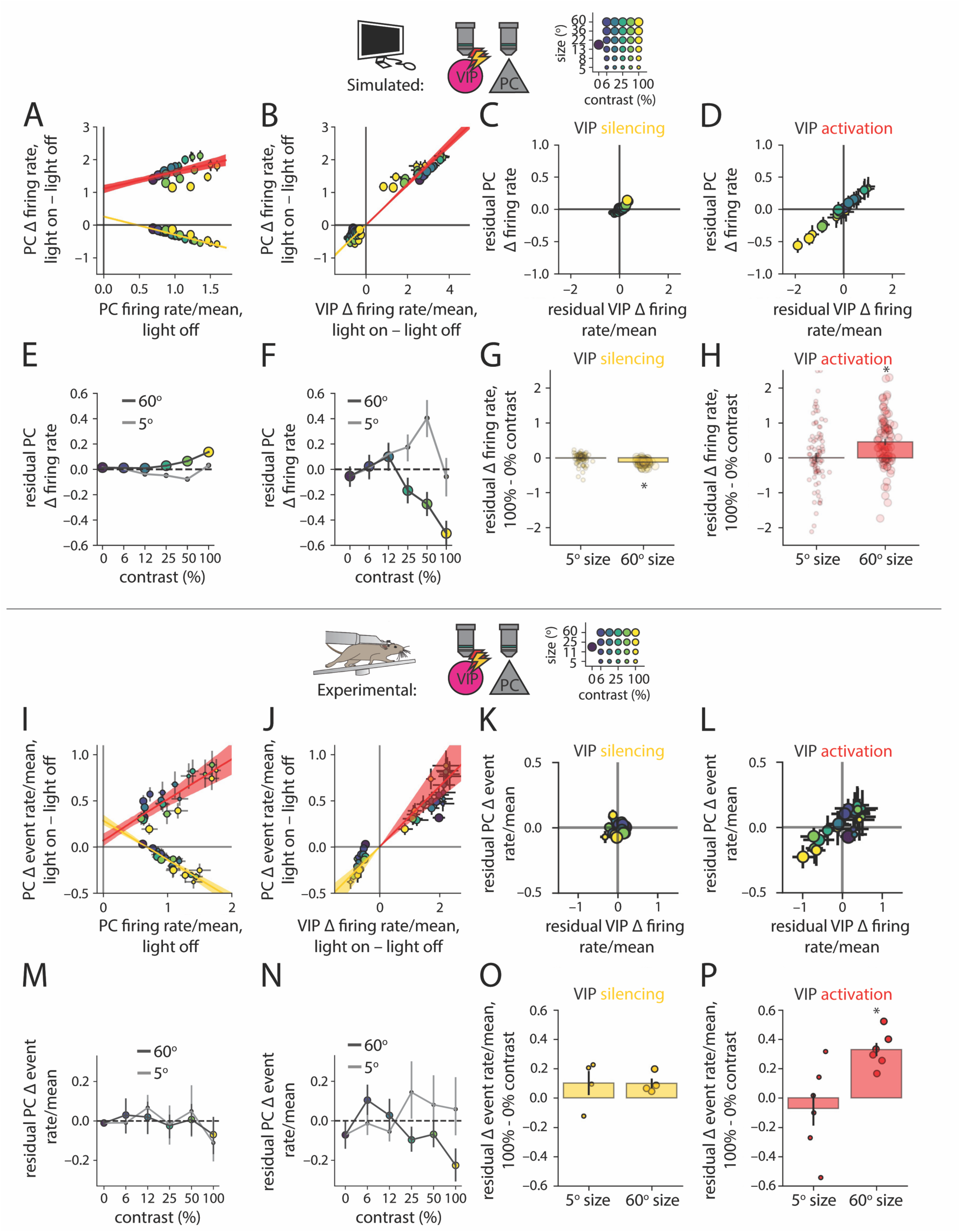
Network supralinearity explains experimental and simulated effects of bidirectional VIP perturbations, in simulation and experiment. A) Scatter plot of PC Δ firing rate vs. control firing rate for simulated finite excitatory and inhibitory current injection to VIP cells. Yellow and red lines represent the least squares linear fit for inhibitory and excitatory current, respectively, with shading indicating bootstrap 68% confidence intervals (*n* = 89 model fits; excluding 5 fits that showed divergent response to at least one connection deletion manipulation. VIP silencing, R=-0.55, p < 10^-10^, and VIP activation, R=0.40, p=3×10^-8^, Wald test). B) Same as A), but plotted against VIP Δ firing rate. C) Residual effect of simulated VIP inhibitory current injection in A) on PCs, not captured by the best fit line, vs. the difference in VIP Δ firing rate from mean across stimuli, for finite inhibitory current injection to VIP cells. Legend for the dots in the scatter plot is given at the top of the figure. D) Same as C), for finite excitatory current injection to VIP cells. E,F) Residual PC Δ firing rate plotted against contrast for 5° and 60° size, for inhibitory (E) and excitatory (F) current injection to VIP cells. G,H) Residual effect at 100% - 0% contrast, for 5° and 60° size, for data in E,F), respectively. Each dot indicates a single successful model fit. Difference of residuals significantly negative (E); p=4×10^-16^, Wilcoxon signed-rank test) and positive (F); p=9×10^-7^, Wilcoxon signed-rank test), respectively, for 60° size. I-P) Same as A-H), but for experimental, rather than simulated, perturbations (I); VIP silencing, R=-0.64, p=2×10^-12^, and VIP activation, R=0.45, p=2×10^-8^, Wald test). Each dot in O,P) indicates a single imaging session (*n* = 4 imaging sessions, silencing, and *n* = 6 imaging sessions, activation). Difference of residuals significantly positive (F); p=0.03, Wilcoxon signed-rank test), for 60° size. In A-F), dot locations indicate mean across model fits, and error bars indicate 16th to 84th percentile of model fits. In I-N), dot locations indicate mean across imaging sessions, and error bars indicate bootstrap SEM equivalent across imaging sessions.

Although much of the effects of VIP perturbations on PC activity could be linearly predicted from the firing rates of neurons in the control condition, a substantial component could not be. Thus, we partitioned the optogenetic effects on PCs into two components: a “linearly predicted” component, and a “residual component” that could not be linearly predicted. Importantly, this residual component might change stimulus preferences, and thus explain the change in contrast dependence of surround suppression we observed. Likewise, we defined residual effects on VIP cells themselves as the difference between the change in VIP cell event rate for a particular stimulus, and the mean change in VIP cell event rate (defined to be constant across stimuli). First, in simulation, we found that small residual effects of VIP cell silencing on PCs were correlated with small residual effects on VIP cells (fig. 9c), and large asymmetric residual effects of VIP activation on PCs were correlated with large residual effects of VIP activation on VIP cells (fig. 9d). Thus, residual effects of VIP activation on PCs were consistent with a shift to a fully VIP-dominated regime at small size and low contrast. Importantly, residual effects depended on both size and contrast, but much more strongly in the case of VIP activation (fig. 9e-h). Thus, the stimulus-dependent shift to a VIP-dominated regime, via residual effects, could explain the greater effect of simulated VIP activation on PC tuning, compared with VIP silencing.

Notably, our model predicted that the network would differentially amplify the network impacts of perturbations to VIP cells depending on the strength of PC activity in control conditions. For example, despite VIP cells’ preference for low contrast, the model predicted a large overall impact of VIP perturbations (including both linearly predicted and residual effects) at the highest contrast and particularly up to intermediate size, where PC activity was high in control conditions. On the other hand, the model predicted smaller effects at low contrast, where PC activity was low in control conditions (fig. 9a). This explained a difference between our model predictions and the predictions of a previous model of this circuit ((Millman et al., 2020); fig. S13b) in which VIP perturbations were predicted to selectively affect PC activity at low contrast.

Our experimental data was remarkably consistent with the predictions of the full recurrent network model. The data showed that higher PC activity in control conditions predicted stronger suppressive effects of VIP silencing (R=-0.64, p=2×10^-12^, Wald test), and stronger facilitating effects of VIP activation (R=0.45, p=2×10^-8^, Wald test) across stimulus conditions (fig. 9i). Further in line with the model, the effects of VIP optogenetics on VIP cells were strongly correlated with the effects on PCs, and all effects were substantially stronger in the case of VIP activation than in the case of VIP silencing (fig. 9j). As in the model, there was a large asymmetric residual effect of VIP optogenetics on PCs (stronger in the case of VIP activation than silencing), correlated with an asymmetric residual effect on VIP cells themselves (fig. 9k,l), and this asymmetric residual effect depended on size and contrast (fig. 9m-p). Artificially subtracting these asymmetric residual components from the effects both of experimental VIP activation and silencing abolished the effects of VIP activation on contrast dependence of surround suppression (fig. S15).

These results empirically confirmed signatures of VIP-SST competition which our network model predicted. Because we found the changes in PC contrast dependence of surround suppression to be consequences of residual effects of VIP perturbations, we next tested whether these residual effects indeed depended on the hypothesized interplay of PC→VIP⊣SST⊣PC and PC→SST⊣PC feedback loops. We selectively deleted excitatory connections within L2/3 necessary for these feedback loops, as before. While deleting PC→SST, PC→VIP, or PC-PV connections, respectively, only weakly affected simulated responses to VIP silencing (fig. S16), they dramatically reduced simulated responses of PCs (fig. 10a,e,i) and VIP cells (fig. 10b,f,j) to VIP activation. More importantly, these manipulations significantly and selectively reduced the residual effects of VIP activation (fig. 10c,g,k), and their contrast dependence (fig. 10d,h,l), but not those of VIP silencing (fig. S16). Thus, the asymmetric contrast-dependent residual effect of VIP activation, which enhanced contrast-dependent surround suppression, was a signature of contrast-dependent VIP-SST competition.

**Figure 10.**
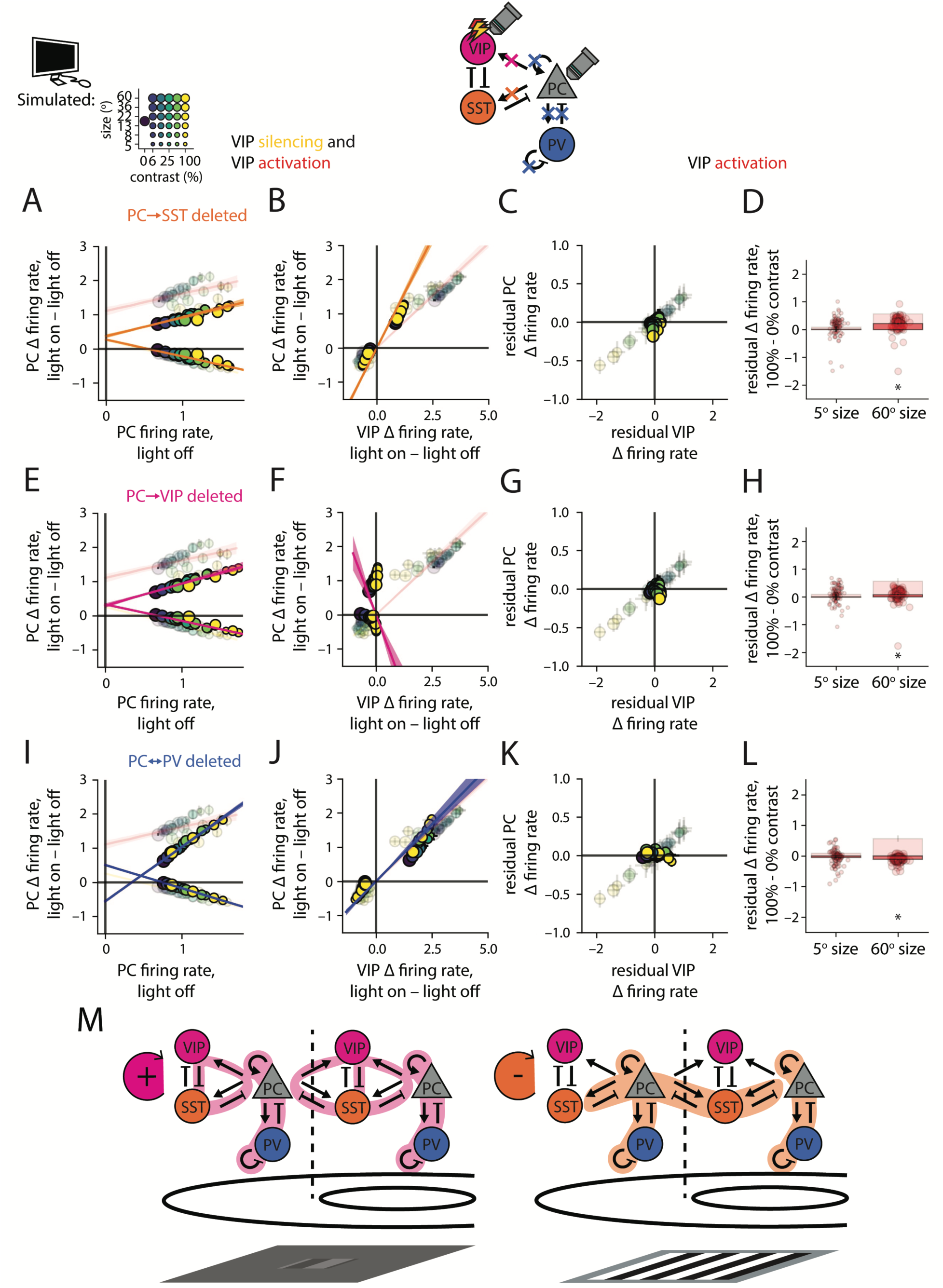
Excitatory connections within L2/3 controlling VIP-SST competition control residual effects of VIP activation. A) Change in firing rate plotted against control (light off) firing rate, both normalized to mean control firing rate across stimuli, in response to finite excitatory or inhibitory current injection to VIP cells after deleting all PC→SST connections. Transparent dots and lines in all panels correspond to the baseline condition, without deletion of any connections (*n* = 89 model fits; excluding 5 fits that showed divergent response to at least one connection deletion manipulation). B) Change in PC activity plotted against the change in VIP activity for the same perturbations as in A). C) Residual change in PC activity, not explained by control firing rate, plotted against the difference in the change in VIP activity, from the mean change in VIP activity for the case of finite excitatory current injection to VIP cells. D) Residual change in PC activity at 100% - 0% contrast, for 5° and 60° size, for the data in C) (difference in residuals significantly reduced by connection deletion, p=6×10^-4^, Wilcoxon signed-rank test). E-H) Same as A-D), but after deleting all PC→VIP connections (difference in residuals significantly reduced, p=7×10^-5^, Wilcoxon signed-rank test). I-L) Same as A-D), but after deleting all PC→PC, PC→PV, PV⊣PC, and PV⊣PV connections (difference in residuals significantly reduced, p=1×10^-7^, Wilcoxon signed-rank test). E-H) Residual effect of VIP silencing and activation in A-D), respectively, defined as the difference between the effect, and the linear prediction based on the control (light off) firing rate. Error bars indicate 16th to 84th percentile of successful model fits.

## Discussion

We combined comprehensive calcium imaging, optogenetic experiments, and computational modeling to address the circuit mechanisms that gate cooperation and competition across cortical space. We examined contrast-dependent surround suppression in the primary visual cortex as a paradigmatic example of such flexible cooperation and competition. Although discovered more than two decades ago (Anderson et al., 2001; Ayaz et al., 2013; Cavanaugh, Bair, & Anthony Movshon, 2002; Cavanaugh, Bair, & Movshon, 2002; Ichida et al., 2007; Nauhaus et al., 2009; Nienborg et al., 2013; Olsen et al., 2012; Ozeki et al., 2009; Polat et al., 1998; Sato et al., 2014; Sceniak et al., 1999; Schwabe et al., 2010; Sengpiel et al., 1997; Tanaka & Ohzawa, 2009; Vaiceliunaite et al., 2013; Van Den Bergh et al., 2010; C. Wang et al., 2009), a circuit mechanism for how the same cortical circuit dynamically shifts between a cooperative and competitive operating regime based on the size and contrast of the visual stimulus has been lacking. As the same cell types and circuits exist throughout cortical areas, the findings of this study may generalize beyond the visual cortex to more broadly explain how cortical circuits adjust cooperation and competition across cortical space to meet task demands.

Our work showed that a transition from positive PC→VIP⊣SST⊣PC feedback at low contrast to negative PC→SST⊣PPC feedback at high contrast was able to modulate the contrast dependence of surround suppression, by supporting contrast-dependent PC competition across space. The PC-PV subnetwork amplified the strength of both feedback loops. Future experiments using SST optogenetic manipulations will be necessary to fully quantify the contribution of these inhibitory circuits to contrast-dependent surround suppression and distinguish competing models (Angelucci, et al. 2017; Rubin et al., 2015). However, it is clear that L2/3 interneuron-mediated competition allows sensitive tuning of contrast-dependent surround suppression, on top of what is inherited from feedforward inputs.

We tested core predictions of this model experimentally by optogenetically perturbing VIP cells. First, in keeping with the supralinearity of the network, both VIP silencing and activation had the strongest effects when PC activity was highest, namely at small size and high contrast. This simple interpretation was qualitatively consistent with the results of other optogenetic perturbations of this circuit (Keller et al., 2020). More generally, it may be helpful to parsimoniously interpret optogenetic effects in terms of their activity dependence, in addition to their stimulus dependence. On top of this activity-dependent effect, VIP activation had a large, asymmetric residual effect that was unexplained by PC activity level, positive at low contrast and negative at large size and high contrast. Our network model explained that this residual effect was a consequence of feedback loops supporting contrast-dependent VIP-SST competition. We found that partitioning optogenetic effects into a component that depended purely on activity level in control conditions, and a residual component that did not, was critical to our interpretation of the optogenetic results. The asymmetric effects of VIP activation and silencing on pyramidal cell tuning were reminiscent of a recent study demonstrating asymmetric effects of activation and inactivation of PV and SST cells on auditory gain and tuning in A1 (Phillips & Hasenstaub, 2016). Bidirectional perturbations, generally, are likely to provide rigorous tests of a predictive understanding of network dynamics and need not conform to our simplest intuitions, particularly in highly recurrent neural networks such as the neocortex.

The data and modeling presented above highlight that understanding strongly recurrently connected networks (within and beyond the cortex), requires a combination of causal perturbations and computational modeling. Without causal perturbations, computational models may be underconstrained. In contrast to the theoretical predictions of a recent study (Millman et al., 2020), we found that the effect of removing VIP activity from the network was not selective for low contrast stimuli, despite the fact that VIP activity was highest for those stimuli in control conditions. On the other hand, inferring recurrent network mechanisms from causal perturbations is likely to be difficult when unsupported by a computational model. The contrast dependence of PC coupling to the VIP and SST subnetworks, which our perturbations revealed, challenged the simplest linear intuitions of how the network might operate. Iteratively developing new intuitions instructed by recurrent network models, such as feedback loops and stabilization, may be crucial for developing a predictive understanding of neural function. Because one study cannot explore all possible causal perturbations, constraining models based on the results of multiple studies will likely become increasingly important. Incorporating the constraint that our modeled network occupy an inhibitory stabilized regime, established by causal perturbations in previous studies (Sanzeni et al., 2020), constrained our model to have strong feedback loops among PCs and PV cells, which in turn powerfully shaped network dynamics in the model. Importantly, some discrepancies remained between model predictions and experimental optogenetic results in this work, particularly in the effects of VIP silencing at low contrast. These discrepancies could perhaps be reconciled in future work by incorporating more experimental measurements to further constrain model fits.

Further, in this study we considered PCs, PV cells, SST cells, and VIP cells as homogenous populations, and restricted most of our analysis to population averages. However, it is already well appreciated that each of the cell types breaks down into distinct sub-classes with unique physiological properties, anatomy, and circuit connectivity. Future work may uncover subpopulations of the interneurons we studied playing dissociable roles. For example, VIP cells are transcriptionally diverse, and molecular markers such as cholecystokinin and calretinin (He et al., 2016), could define functional subclasses of VIP cells. Tuning diversity of VIP cells, like transcriptional diversity, has also been found to correlate with cortical depth (Dipoppa et al., 2018). Diversity in tuning properties may align with diversity of functional roles in the cortical microcircuit. A recent study in the somatosensory barrel cortex found that a subpopulation of VIP cells co-expressing ChAT exert an inhibitory influence on PCs specifically in response to weak stimuli (Dudai et al., 2020). A combination of intersectional genetic techniques and optogenetics, or two-photon optogenetics, could yield further insight in the functional roles of distinct VIP populations in sensory coding. Furthermore, other 5-HT3aR+ interneurons not expressing VIP, particularly in layer 1, have been shown to have similar tuning for sensory signals as VIP cells (Abs et al., 2018; Cohen-Kashi Malina et al., 2021; Mesik et al., 2015). Future work could clarify the extent to which these cell types act in concert with VIP cells or play disparate roles to control sensory coding. In addition to discrete variability between sub-classes of each cell type, continuous variability could be important, both in synaptic connections and in tuning properties. Future work could seek to model this variability (Beiran et al., 2021), which might generate further hypotheses to be tested with more sophisticated optogenetic perturbations.

Translaminar inhibitory as well as excitatory connections are also likely to be important, particularly as VIP perturbations affected L4 as well as L2/3 activity. Future work expanding the scope of these experiments could clarify the extent to which translaminar feedforward and feedback connections shape the response properties of these cell types. Further, these cell types might differentially integrate feedback signals from higher areas (Keller et al., 2020; Zhang et al., 2014), which could be important for their encoding of homogeneous textures as well as richer stimuli.

Although we argue that the influence of VIP cells is likely to be important for network regimes optimizing detection, this study has examined only passively viewing mice, not animals motivated to detect weak signals. Behavioral context and salience of sensory inputs has been found to powerfully shape the tuning properties of these interneuron types (Kuchibhotla et al., 2017), possibly to enhance coding properties of pyramidal cells. Future work combining two-photon imaging of these cell types with behavior and network modeling could suggest possible dynamical mechanisms for this enhancement.

Despite these limitations, there is growing evidence that the circuit motifs examined in this study may generalize to other areas of the brain. In auditory cortex, a series of recent studies have uncovered analogous tuning properties and functional roles of the same interneuron types. VIP cells show non-monotonic tuning for sound amplitude (analogous to contrast) (Mesik et al., 2015), while SST cells are insensitive to all but high amplitude tones (L. Li et al., 2015). Whereas PCs commonly prefer narrow spectral bandwidth (analogous to size) (H. Li et al., 2019), SST cells prefer broader spectral bandwidth (Lakunina et al., 2020). Silencing SST cells results in a reduction of lateral inhibition (Kato et al., 2017) and a facilitation of PC responses to broad spectral bandwidth sounds (Lakunina et al., 2020). Future work could examine the extent to which the L2/3 microcircuit in V1 and A1 occupies a similar dynamical regime, and examine tuning properties of interneurons in S1 for space and intensity. How these insights extend to non-sensory areas, such as frontal or parietal cortices (Pi et al., 2013), is an exciting area for future study.

---

**Figure S1.**
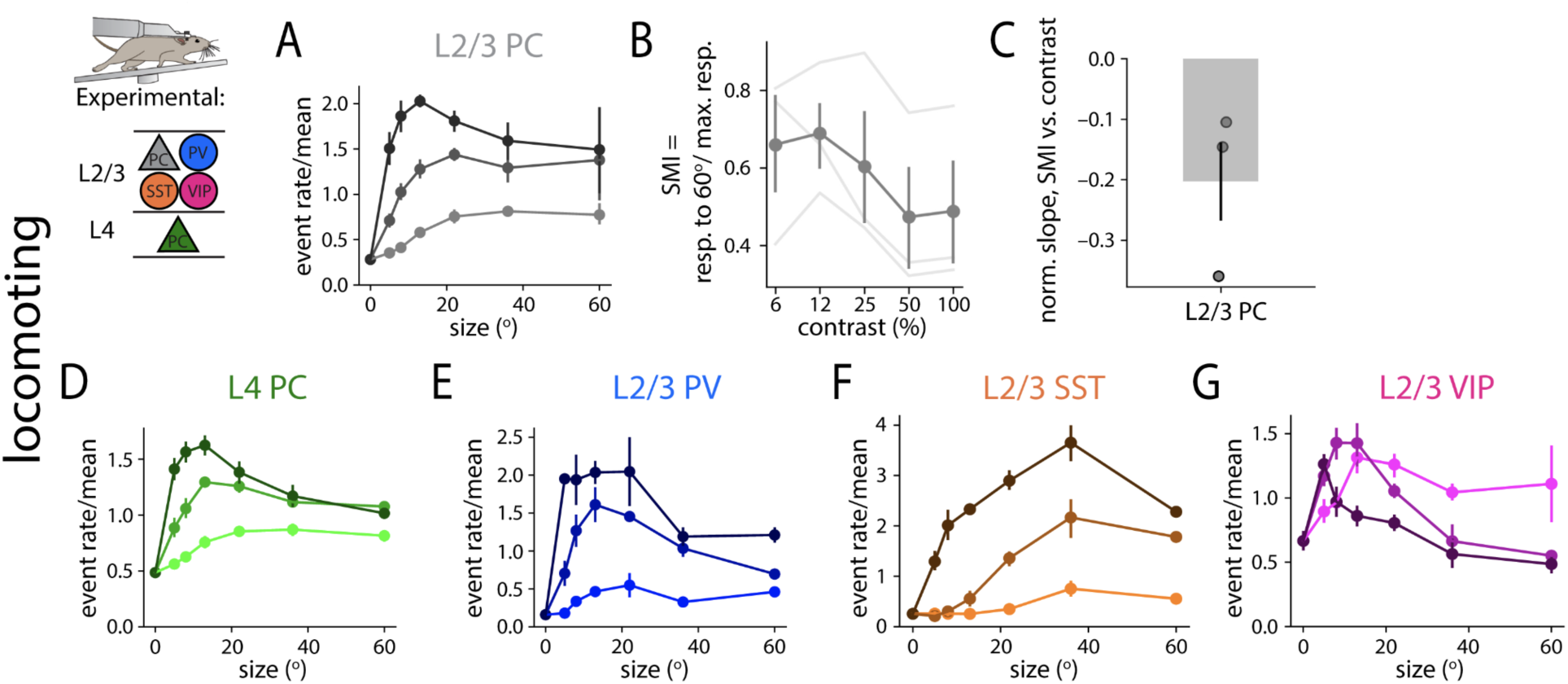
Similar cell-type specific responses to stimulus size and contrast during locomotion. A) Size tuning curves of L2/3 PCs in the locomoting condition, averaged across experiments. B) Surround modulation index (SMI; response to 60° size divided by response to preferred size) of L2/3 PCs as a function of contrast in the locomoting condition. C) Change in SMI between 6% and 100% contrast for L2/3 PCs. D-G) Same as A), but for L4 PCs, L2/3 PV cells, L2/3 SST cells, and L2/3 VIP cells, respectively. (*n* = 8, 6, 2, 4, 6 imaging sessions, respectively, for A-E. Note that SMI could be computed only for the subset of 3 L2/3 PC imaging sessions at which the 60° size stimulus was shown) Error bars indicate bootstrap SEM equivalent.

**Figure S2.**
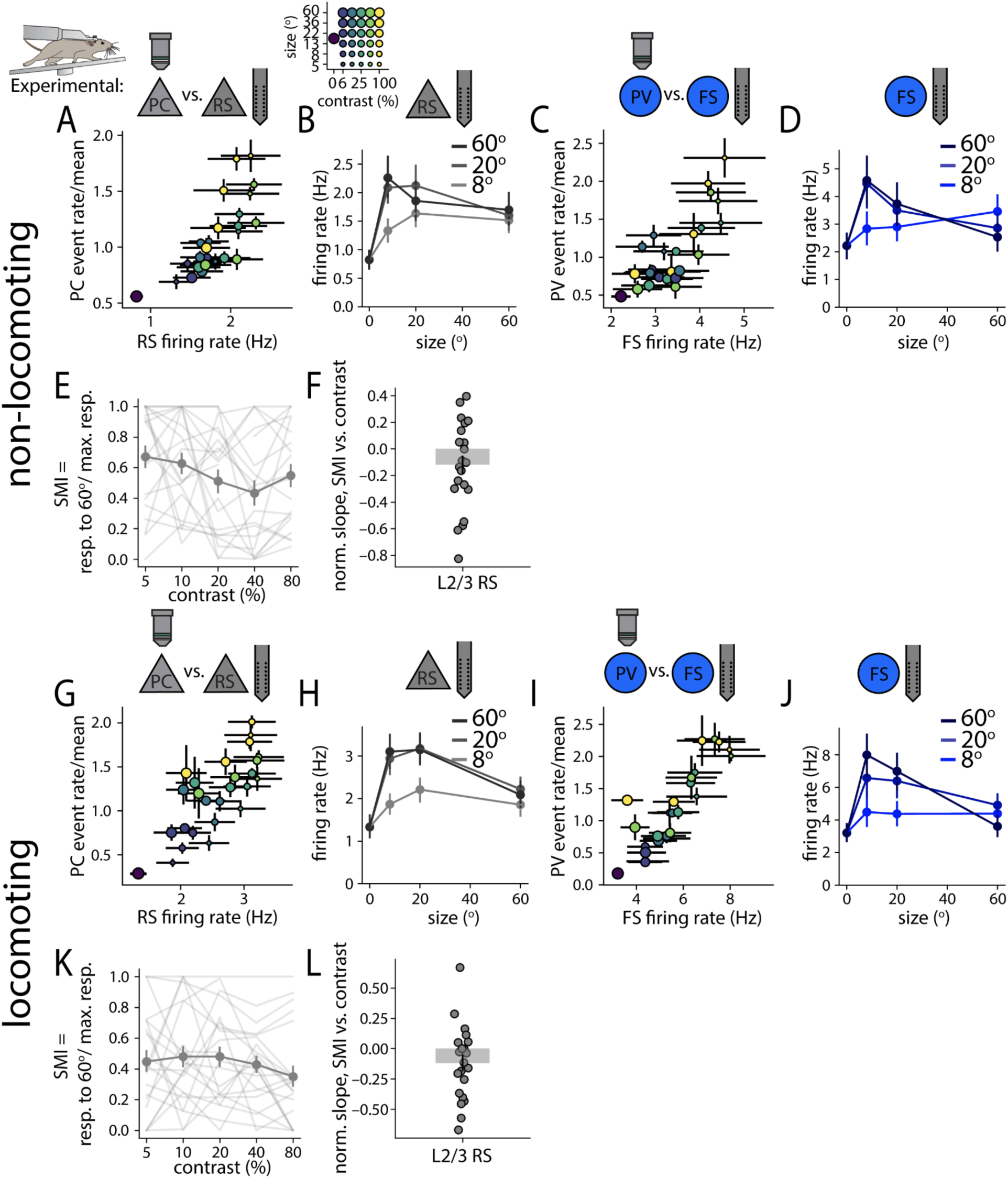
Extracellular electrophysiology reveals qualitatively similar tuning properties to calcium imaging. A) Scatter plot of firing rate of regular spiking (RS) units recorded using extracellular electrophysiology, vs. PC event rate/mean across stimuli measured using calcium imaging for the equivalent stimuli, in the non-running condition (significantly correlated, R=0.84, p=7×10^-9^, Wald test). Dot size and color are as in fig. 2B. B) Size tuning curves, separated by contrast, for RS units plotted in A). C,D) Same as A,B), but for fast spiking (FS) units recorded using extracellular electrophysiology, vs. PV event rate/mean measured using calcium imaging. E) Surround modulation index (SMI), defined as the response to 60° gratings divided by response to the preferred size gratings, as a function of contrast, for L2/3 RS units. Each line is a single recording session, and the bold line corresponds to average across recording sessions. Note that due to the small numbers of neurons sampled in electrophysiological recordings relative to calcium imaging recordings, within-session measurement errors are larger here and in following panels than e.g. in fig. 1E. F) Difference of SMI between 6% and 100% contrast. Each dot corresponds to one recording session. G-M) Same as A-F), for the locomoting condition.

**Figure S3.**
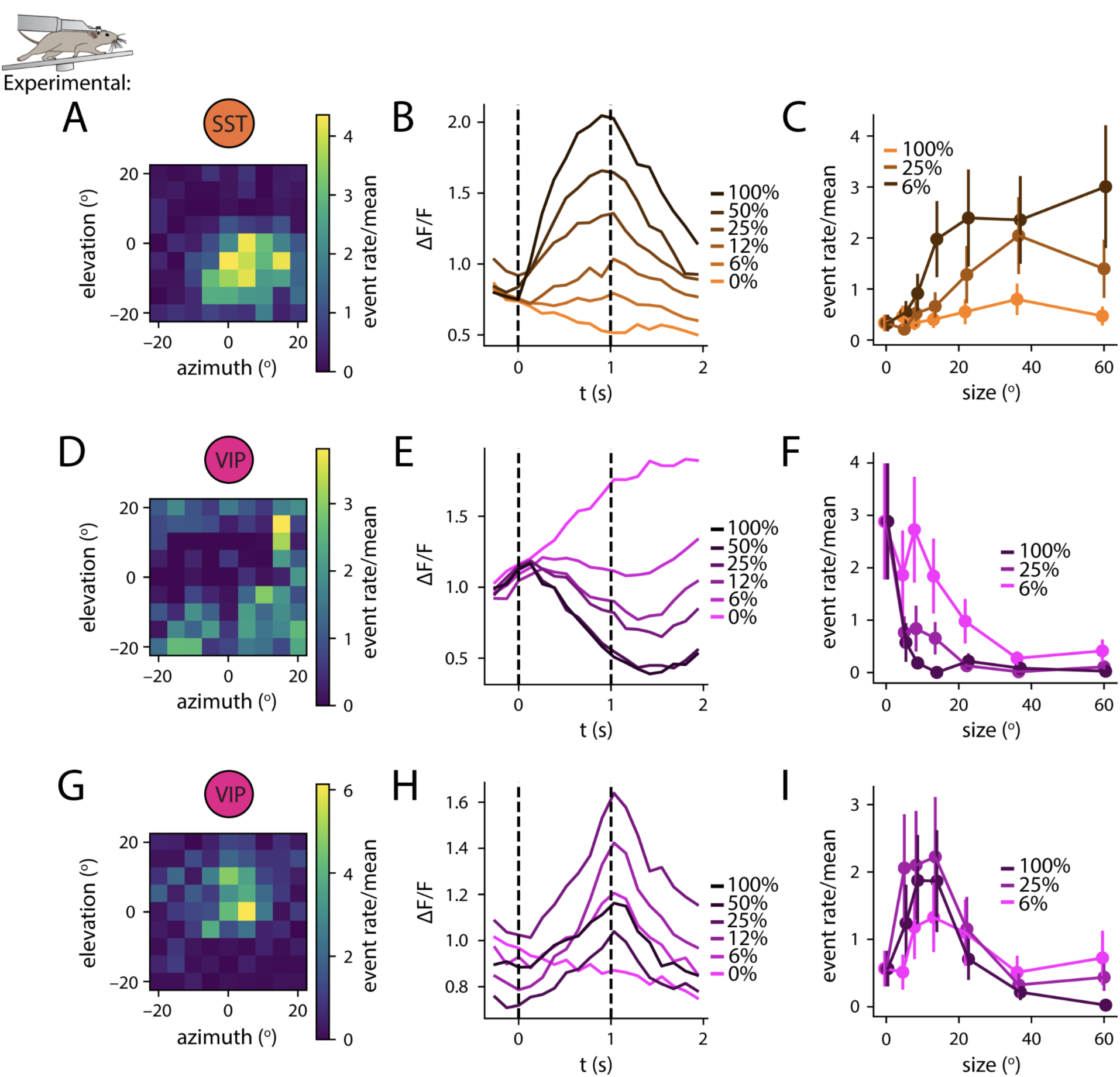
Example tuning curves show divergence of VIP cell size-contrast tuning from SST cells, and between-VIP-cell diversity. A) Receptive field of an example SST cell (see Methods). Mapped using 10° drifting gratings of time-varying direction, shown for 1.5 seconds each. B) ΔF/F time course of size-contrast responses for the neuron shown in A; each curve represents the response to gratings of a given contrast, averaged across sizes and orientations. C) Size tuning of the same cell, split up by contrast. D-F) Same as A-C), for an example contrast-suppressed VIP cell. G-I) Same as A-C), for an example contrast-facilitated VIP cell. Error bars indicate bootstrap SEM equivalent.

**Figure S4.**
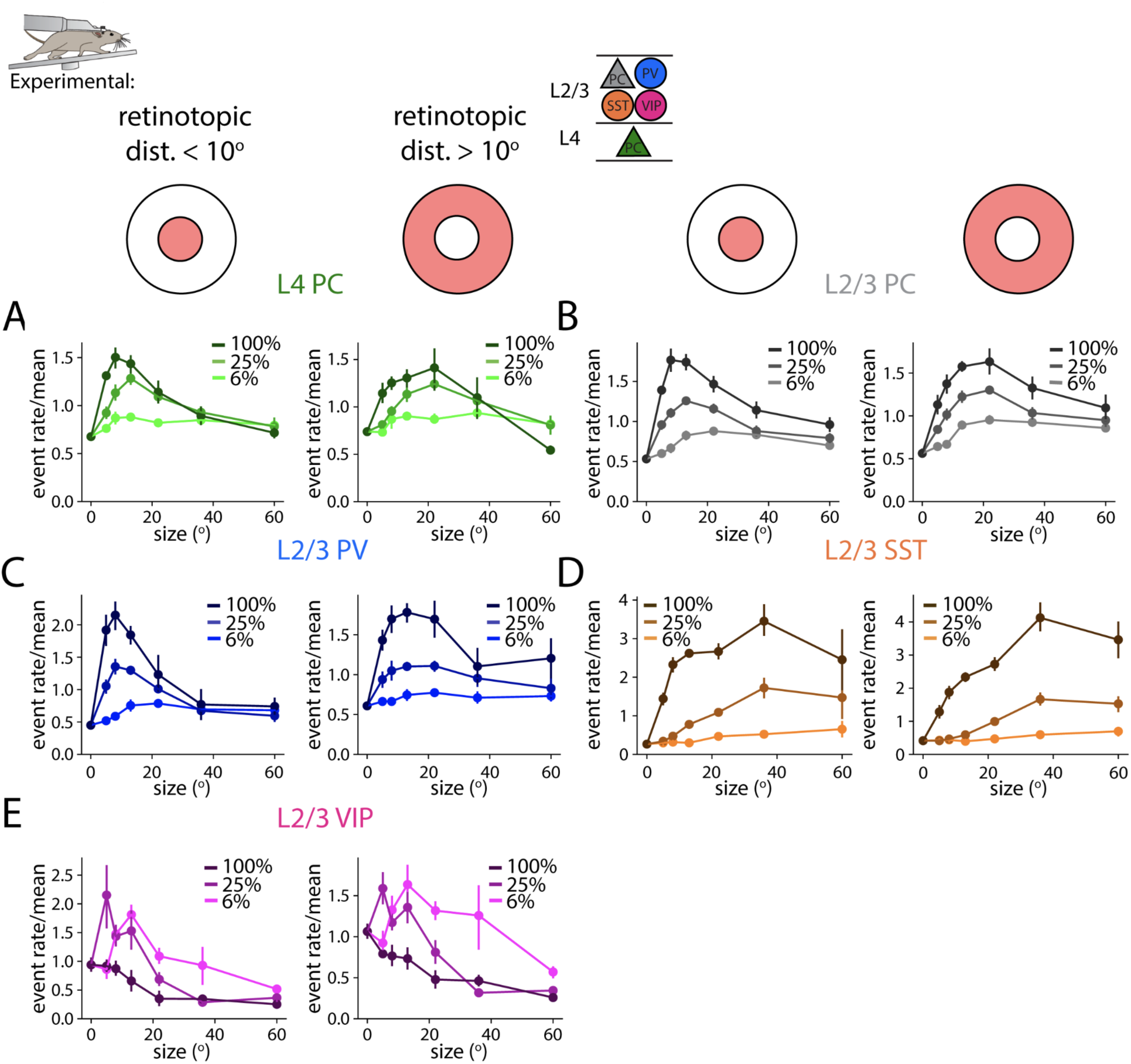
Neurons with receptive fields not aligned with the stimulus center show preference for larger sizes. A) Size tuning of L4 PCs, including only those located within 10 visual degrees of retinotopic space of the center of the stimulus representation (left). Beyond 10 visual degrees (right). B-E) Same as A), for L2/3 PCs, PV cells, SST cells, and VIP cells, respectively. (Aligned: *n* = 6, 9, 6, 5, 5 imaging sessions, respectively, for A-E. Not aligned: *n* = 4, 11, 7, 8, 6 imaging sessions, respectively, for A-E.) Error bars indicate bootstrap SEM equivalent.

**Figure S5.**
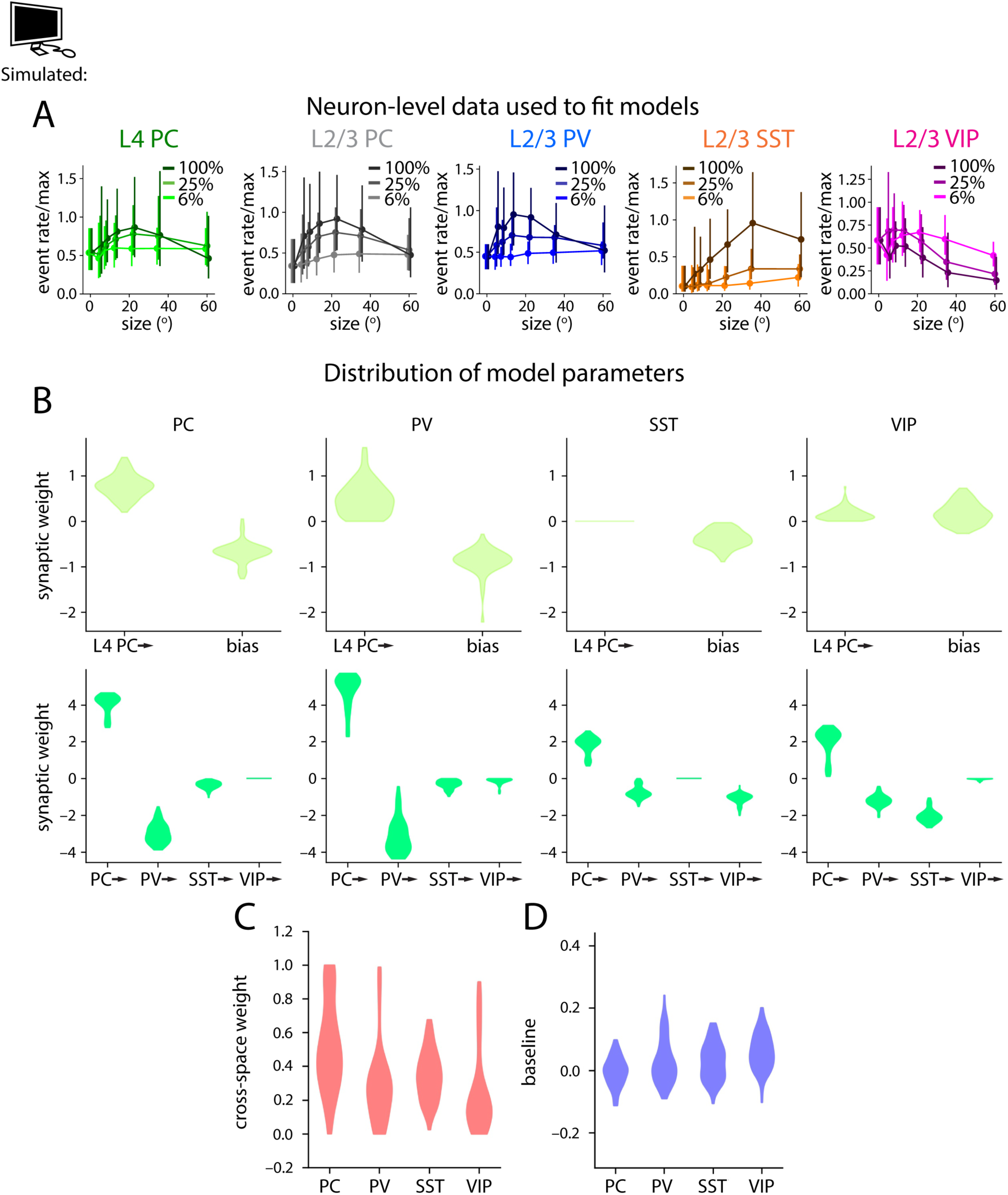
Well-performing model fits show variability in fit parameters. A) Model fits were performed on neuron-level, data, rather than averages within imaging sessions as displayed elsewhere in the paper (see Supplementary Note). Shown are size tuning data used to fit models for each of the five cell types, with error bars represented as standard deviations, rather than SEM, as elsewhere where experimental data is presented, to illustrate the degree of diversity between neurons within a cell type. Each neuron’s responses are first individually normalized to the mean response across stimuli, and then all neurons’ responses are normalized together such that the maximum neuron-averaged value across stimuli is 1. B) Violin plots showing variability in synaptic input connection weights among model fits for each of the four L2/3 cell types modeled. C) Cross-space weight for each cell type. D) Baseline for each cell type.

**Figure S6.**
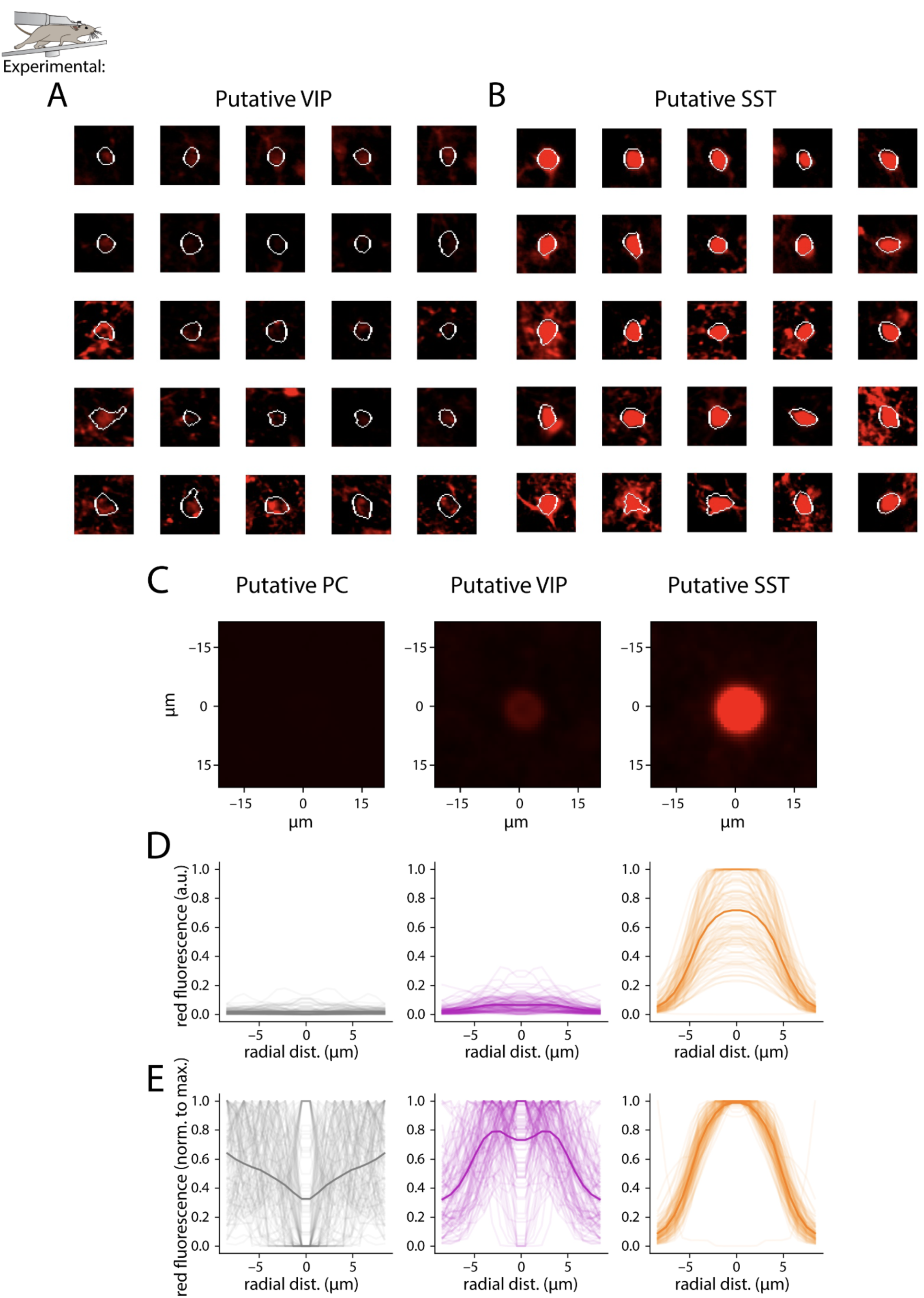
SST-tdTomato and VIP-mRuby3 cells can be distinguished by brightness and spatial profile. A) Outlines of putative VIP (membrane-bound mRuby3 expressing) cells, with red channel images. B) Analogous for putative SST (cytosolic tdTomato expressing) cells. Candidate labeled cells were identified based on having (1) high mean and (2) high coefficient of variation of red intensity (thus, identifying cells with bright red labeling, mostly cytosolic, or labeling that was highly variable across the ROI, mostly membrane-bound). Among these candidates, true labeled cells were manually identified. Putative VIP and SST cells were then separated based on a manually selected cutoff of mean red intensity (SST cells higher red intensity, and VIP cells lower red intensity). C) Average red channel images for unlabeled cells (putative PCs; left), VIP cells (middle), and SST cells (right), aligned to the center of each cell. D) Red intensity (a.u.) as a function of radial distance from the center of the ROI. Each transparent line is one cell, and the bold line is the average. On the left, in gray, are unlabeled cells (putative PCs), in the center, in magenta, are putative VIP cells, and on the right, in orange, are putative SST cells. E) Same plots as D), but normalized to the maximum value.

**Figure S7.**
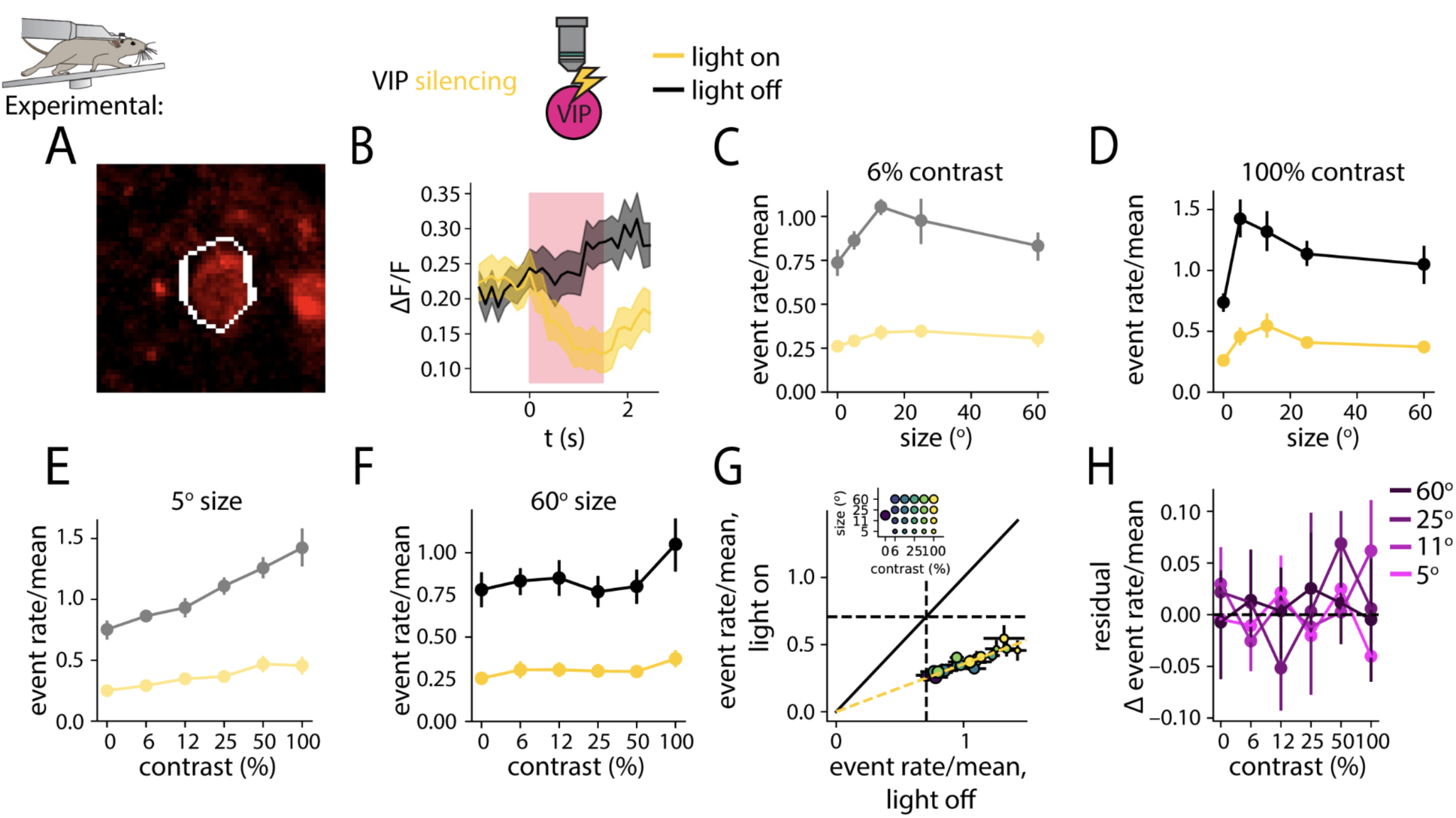
Optogenetic silencing of VIP cells. A) eNpHR3.0-mRuby3-labeled VIP cell from a VIP silencing experiment. B) ΔF/F averaged across all VIP cells, for gray screen trials following gray screen trials with no optogenetic stimulation. In yellow is the average trace for trials where the optogenetic stimulation light was on, and in black is the average trace for trials where the optogenetic stimulation light was off. In pink is the duration of optogenetic stimulation, in the case of the yellow trace. C) Response of a VIP cell to gratings of varying size, at 6% contrast. In black is the light off response, and in yellow is the light on response. D) Same as C), for 100% contrast. E,F) Same as C), D), but for varying contrast, at 5° and 60° size, respectively. G) Scatter plot of light on vs. light off event rate for putative VIP cells at varying size and contrast. In yellow is the best fit line, with the intercept constrained to be 0. Dot size and color are as in fig. 3A. H) Residual effect of optogenetic perturbation not captured by the best fit line in G), for varying contrast, and at varying sizes. Shading and error bars represent bootstrap SEM equivalent.

**Figure S8.**
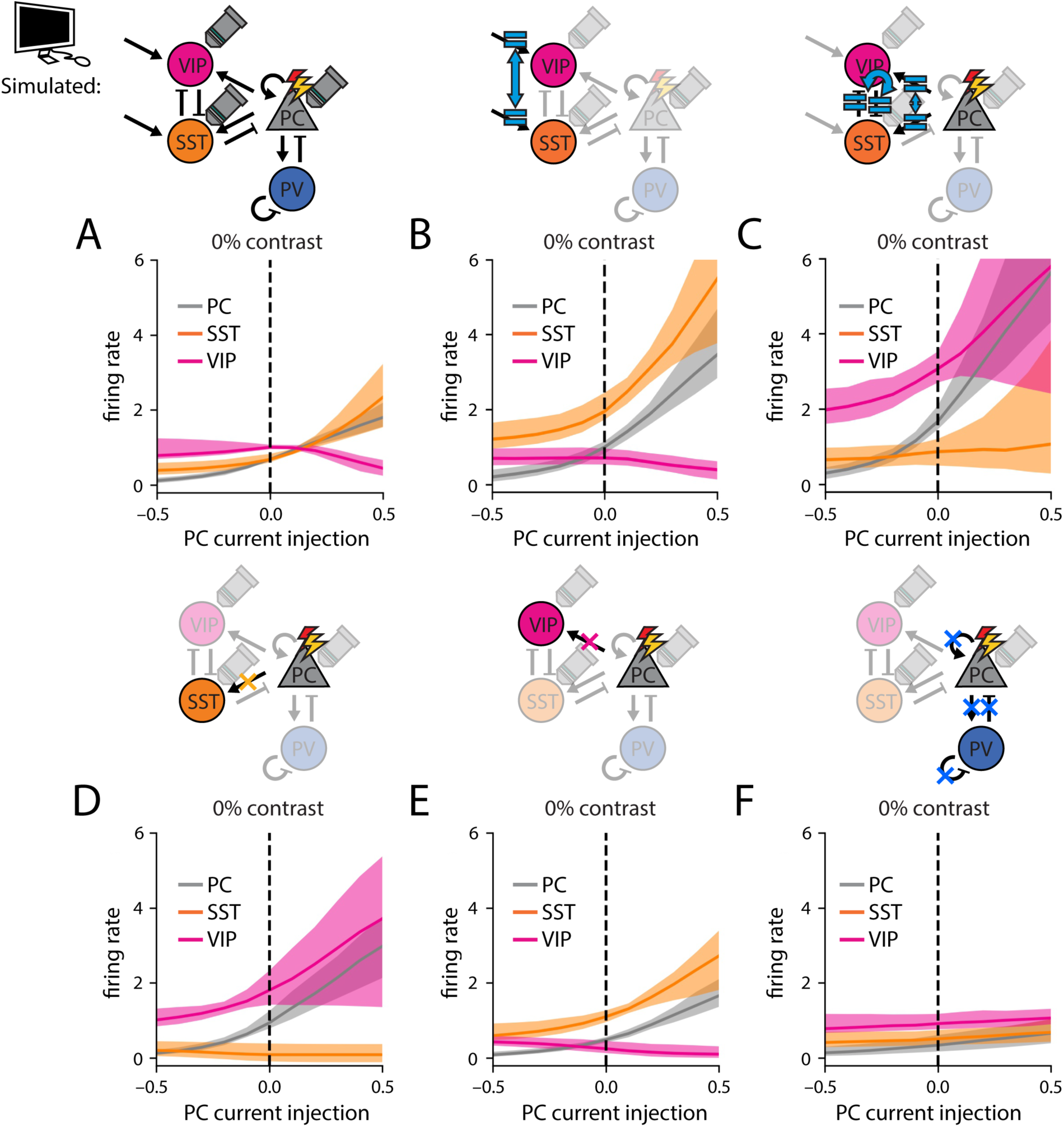
Excitatory connections within L2/3 shape VIP-SST competition. A) The firing rate of center PCs (gray), SSTs (orange), and VIPs (magenta) in response to the injection of finite positive or negative current to center PCs, plotted as firing rate vs. injected current, starting from the 0% contrast condition (*n* = 89 model fits). B) Same as A), but setting VIP bias current equal to SST bias current. C) Same as A), but setting PC→SST connection weights to equal PC→VIP connection weights, and SST⊣VIP connection weights to equal VIP⊣SST connection weights. D-F) Same as A), but deleting all PC→SST, PC→VIP, or all PC←→PC, PV⊢→PV, and PV⊢⊣PV connections, respectively. In B-F), the stated manipulations were performed both on center and surround circuits. Shading indicates 16th to 84th percentile of successful model fits.

**Figure S9.**
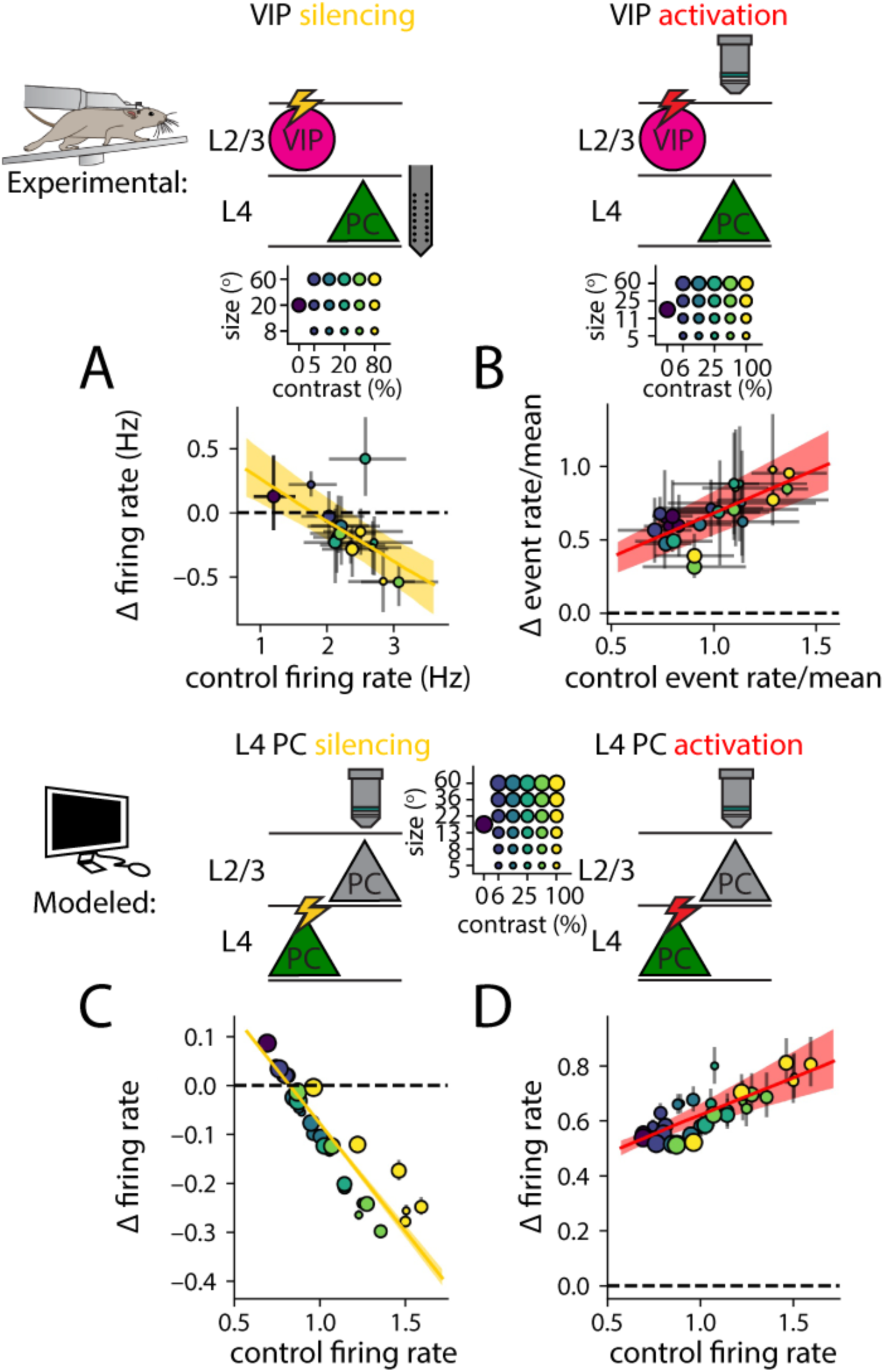
VIP perturbation modulates feedforward inputs to L2/3. A) Effect of VIP silencing on L4 regular spiking units measured using extracellular electrophysiology. Change in measured firing rate, normalized to mean light off firing rate, is plotted against light off firing rate, normalized to its mean, averaged across mice. Dot size and color are as in fig. 2B. B) Effect of VIP activation on L4 PCs measured using calcium imaging. Change in measured event rate, normalized to mean light off firing rate, is plotted against light off firing rate, normalized to its mean, averaged across mice. C) Response of L2/3 PCs to a modeled DC decrease in L4 PC firing rate across stimulus conditions. Change in firing rate, normalized to mean, is plotted against baseline firing rate. D) Same as C), but for a DC increase in L4 PC firing rate, across stimulus conditions. Error bars indicate bootstrap SEM equivalent.

**Figure S10.**
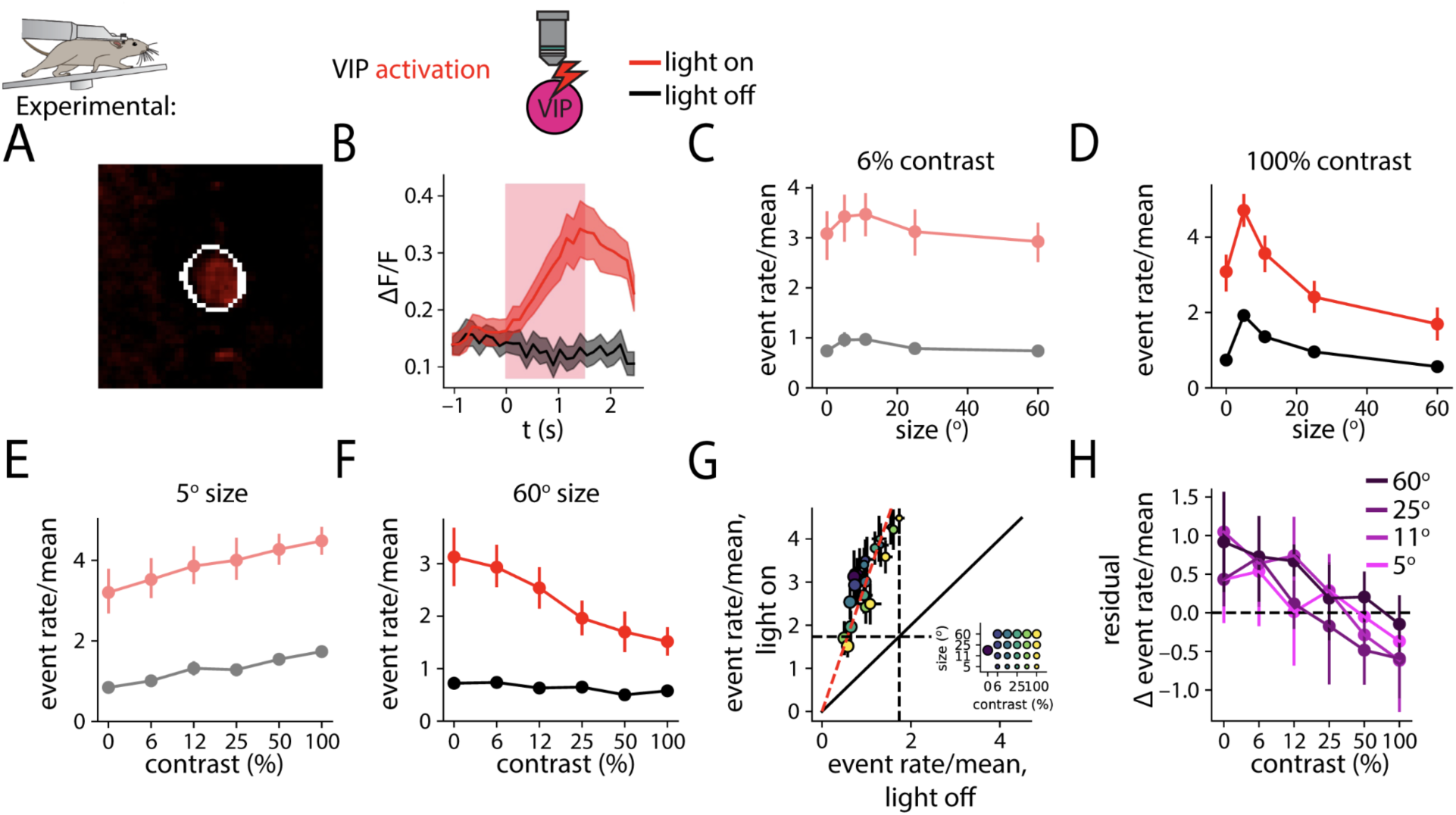
Optogenetic activation of VIP cells. A) ChrimsonR-tdTomato-labeled VIP cell from a VIP activation experiment. B) ΔF/F averaged across all VIP cells, for gray screen trials following gray screen trials with no optogenetic stimulation. In red is the average trace for trials where the optogenetic stimulation light was on, and in black is the average trace for trials where the optogenetic stimulation light was off. In pink is the duration of optogenetic stimulation, in the case of the red trace. C) Response of a VIP cell to gratings of varying size, at 6% contrast. In black is the light off response, and in red is the light on response. D) Same as B, for 100% contrast. E,F) Same as C), D), but for varying contrast, at 5° and 60° size, respectively. G) Scatter plot of light on vs. light off event rate for putative VIP cells at varying size and contrast. In red is the best fit line, with the intercept constrained to be 0. Dot size and color are as in fig. 2B. H) Residual effect of optogenetic manipulation not captured by the best fit line in G), for varying contrast, and at varying sizes. Shading and error bars indicate bootstrap SEM equivalent.

**Figure S11.**
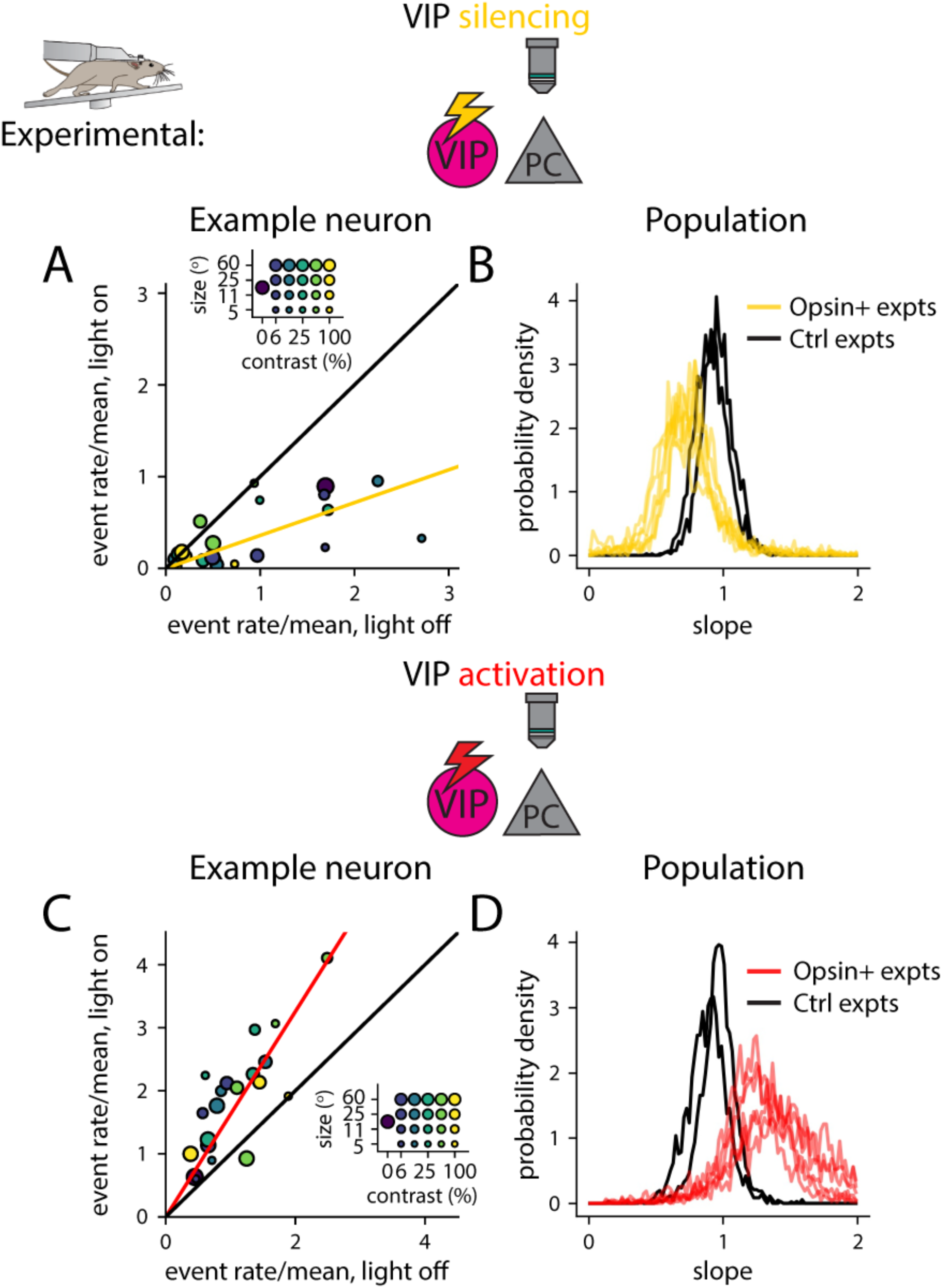
VIP perturbations bidirectionally control responses of non-opsin-expressing cells. A) Scatter plot of size-contrast tuning responses for an example unlabeled neuron from a VIP-eNpHR3.0 expressing mouse. Light on responses are plotted vs. light off responses. In yellow is the best-fit slope transforming light-off to light-on responses, and in black is the unity line (slope = 1). Dot size and color are as in fig. 2B. B) Histogram of best-fit slopes for optogenetic experiments (yellow) and control experiments (black) in which identical illumination parameters were used. Each line corresponds to one imaging session. C,D) Analogous to A), B) for VIP-ChrimsonR experiments, with optogenetic experiments represented in red, and control experiments in black.

**Figure S12.**
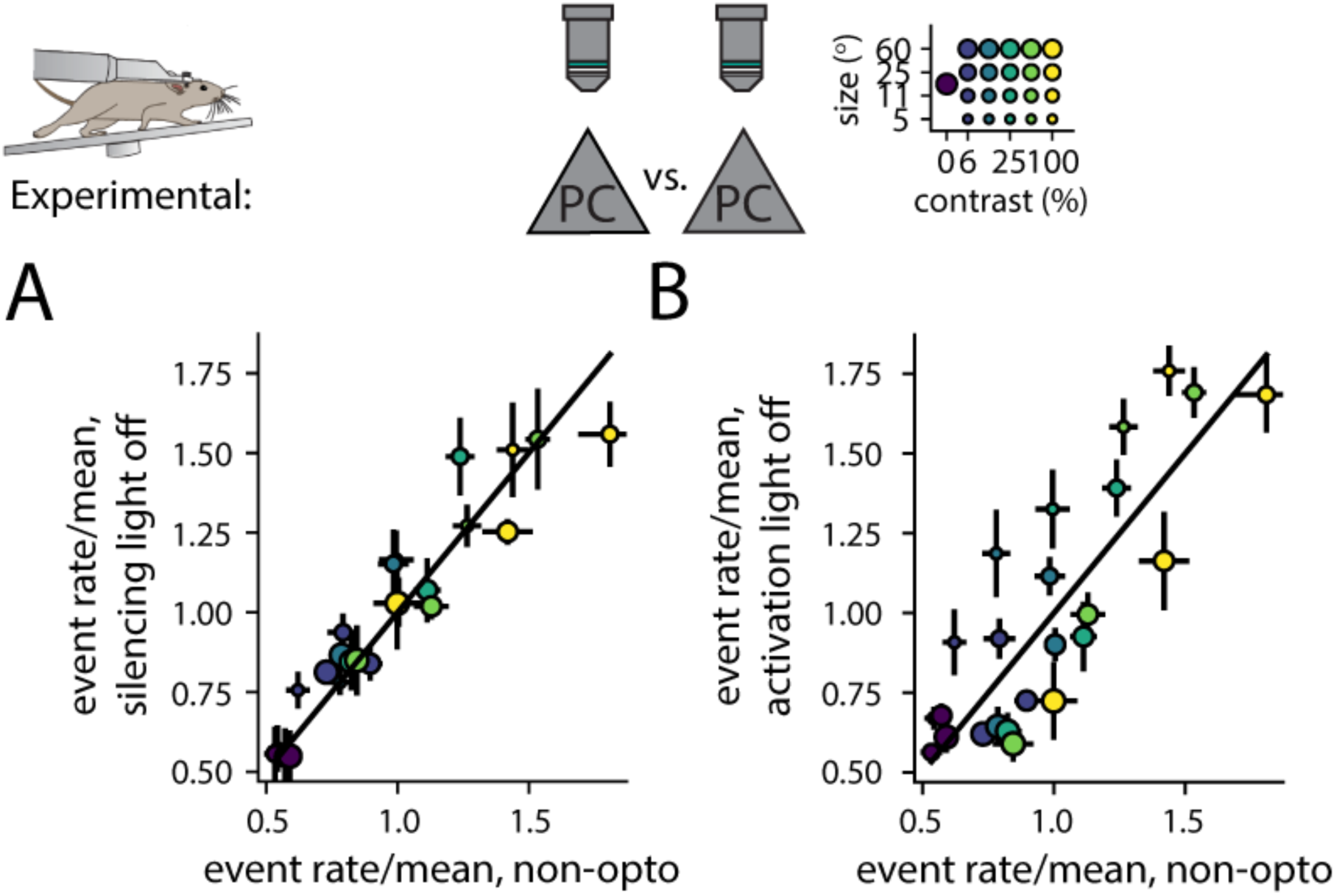
Measured control condition activity in calcium imaging and optogenetic experiments matches activity measured in only-calcium imaging experiments. A) Scatter plot of event rate, normalized to mean, for synapsin-GCaMP6s responses in VIP silencing experiments, with the optogenetic stimulation light off, vs. transgenic GCaMP6s responses of pyramidal cells in non-optogenetic experiments, for the analogous stimuli (R=0.95, p=3×10^-12^, Wald test). Dot size and color are as in fig. 3A. B) Same as A), but for responses in VIP activation, rather than silencing, experiments (R=0.84, p=3×10^-7^, Wald test). Error bars indicate bootstrap SEM equivalent.

**Figure S13.**
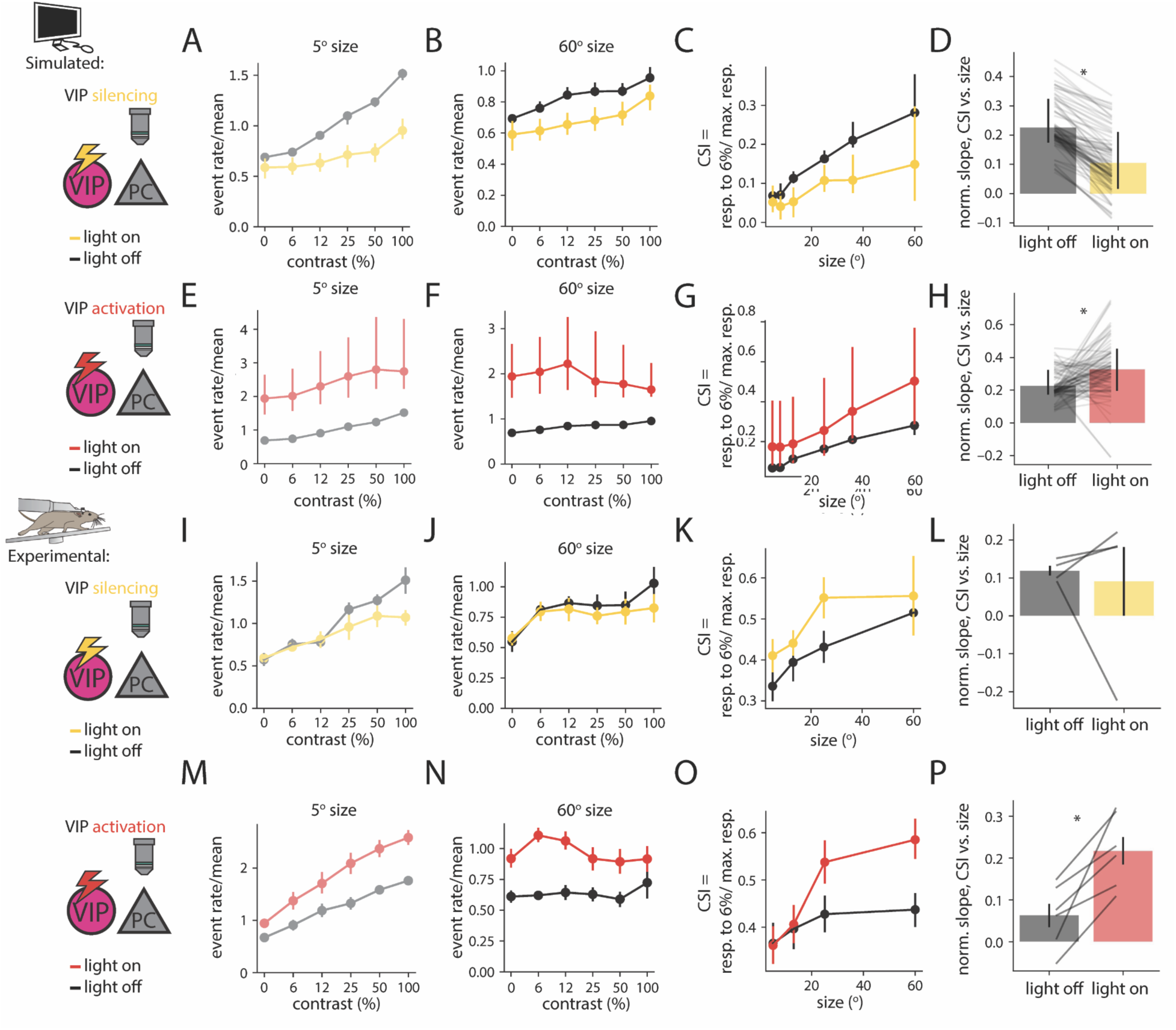
VIP activation enhances size dependence of contrast sensitivity. A) Plot of simulated PC event rate versus grating contrast for 5° size in the control, light off condition (light off, black) and during simulated optogenetic suppression of VIP cells (light on, yellow). B) Same as A) for 60° size. C) Contrast sensitivity index (CSI) is plotted as a function of size in the light on and light off conditions. D) Change in CSI vs. size slope between the light off and light on condition. Each horizontal line represents an individual model fit. E-H) Same as A-D), but for simulated optogenetic activation of VIP cells. I-P) Same as A-H), but for experimental silencing and activation and silencing of VIP cells. Horizontal lines in L,P) correspond to individual experiments. Error bars in A-H) indicate 16th to 84th percentile of successful model fits, while error bars in I-P) indicate bootstrap SEM equivalent across individual experiments.

**Figure S14.**
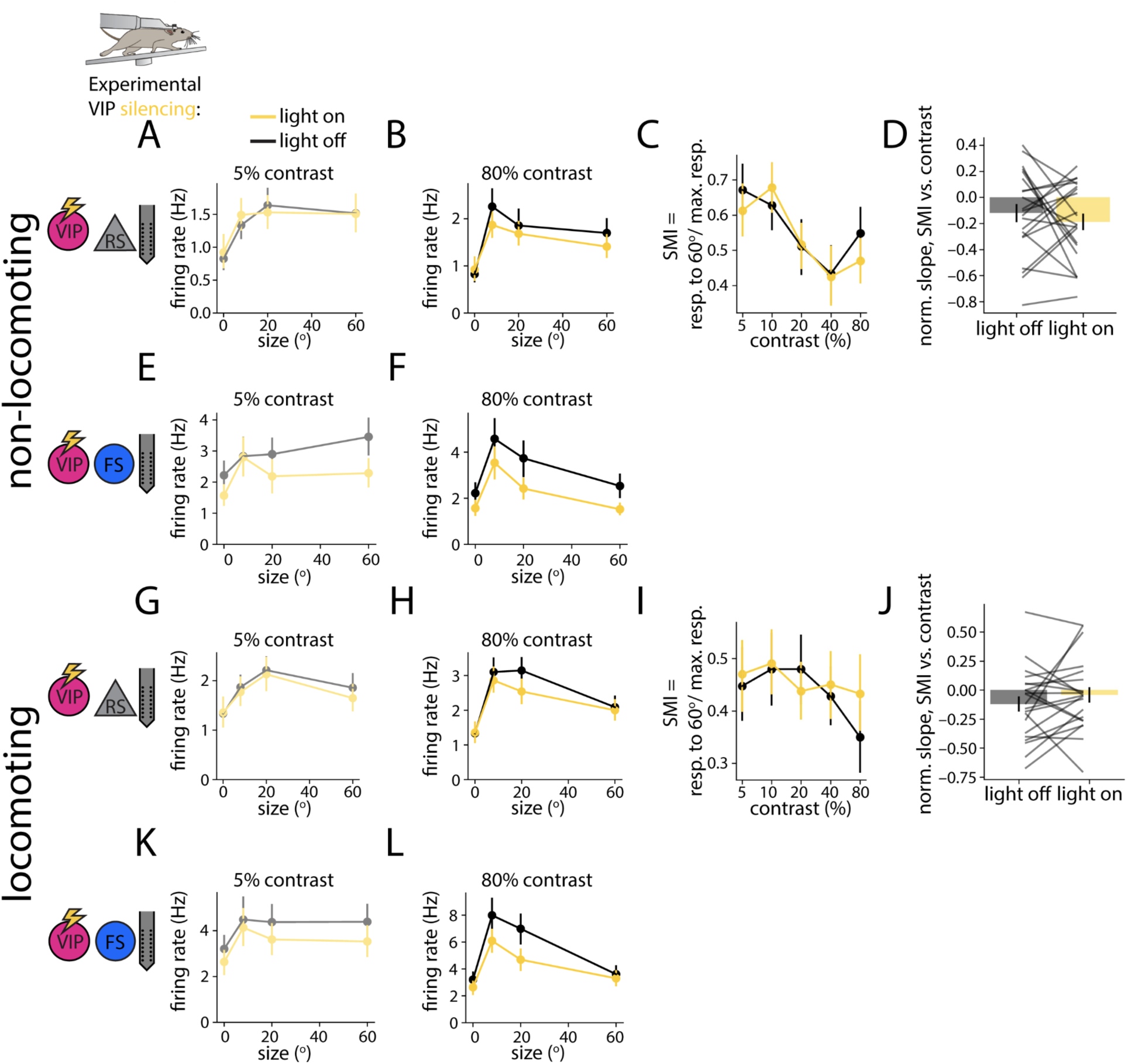
Extracellular electrophysiology confirms VIP silencing does not affect contrast dependence of surround suppression. A) Response of regular spiking (RS) units recorded using electrophysiology to drifting gratings of varying size, at 5% contrast, in the non-running condition. B) Same as A, but at 80% contrast. C) Surround modulation index is plotted as a function of contrast, in the light on and light off case. D) Change in SMI between 6% and 100% contrast is plotted, for the light on and light off cases. Each line represents the average of units recorded over one experiment. E-H) Same as A-D), but for fast-spiking (FS) units. I-P) Same as A-H), but for the running condition. The light on and light off conditions are not significantly different in D), H), L), or P), p > 0.05. Error bars indicate bootstrap SEM equivalent.

**Figure S15.**
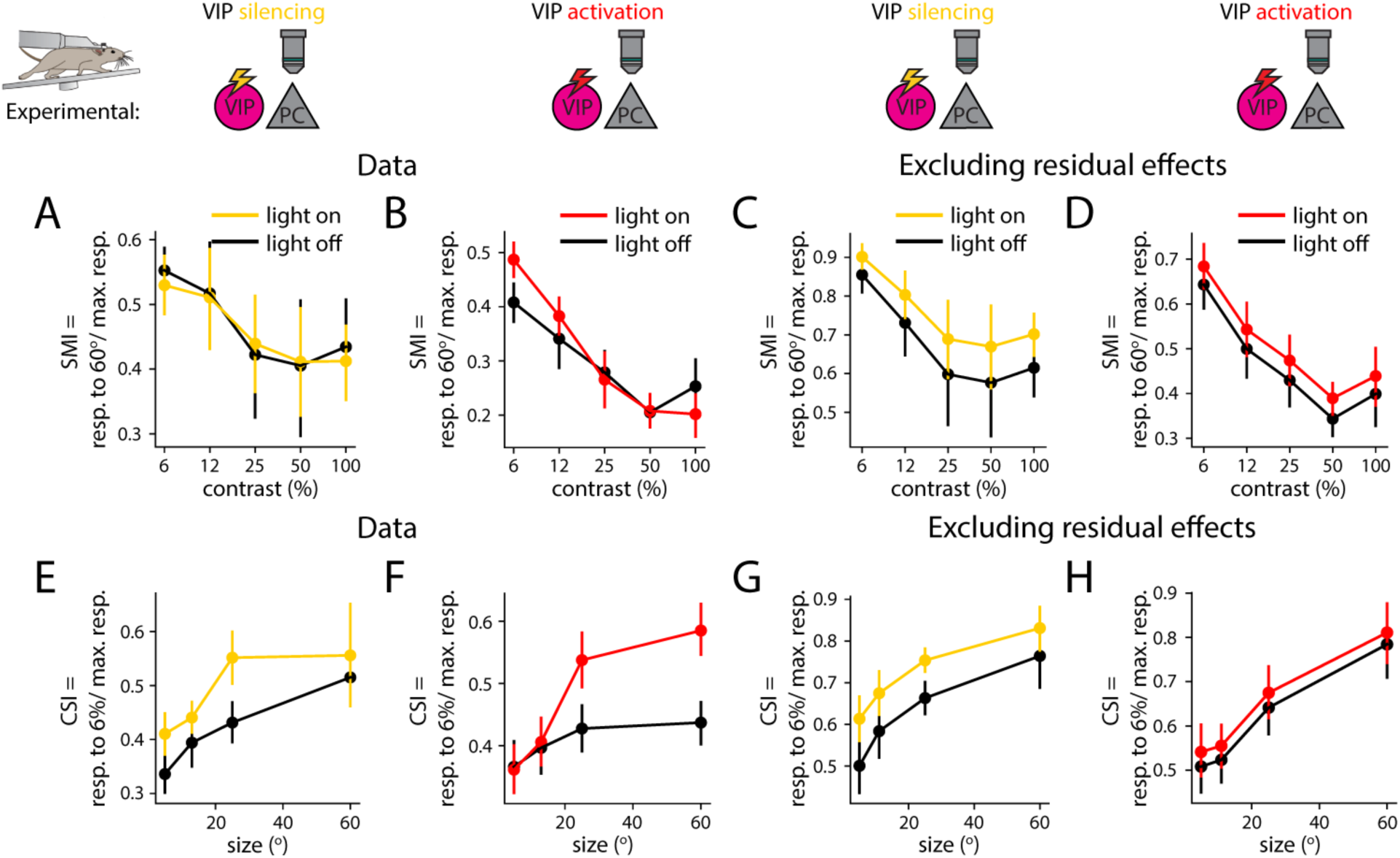
Linear firing rate transformations do not explain enhancement of contrast dependence of surround suppression. A,B) Fig. 5C,K are reproduced here for clarity. C) As in A), surround modulation index is plotted as a function of contrast, for the VIP silencing data. In black is the light off condition. In red is plotted, rather than the true light on condition, the prediction of a linear fit to the light on condition, based on the light off condition. D) Same as C), for the VIP silencing case. E-H) Same as A-D), for contrast sensitivity index. Error bars indicate bootstrap SEM equivalent.

**Figure S16.**
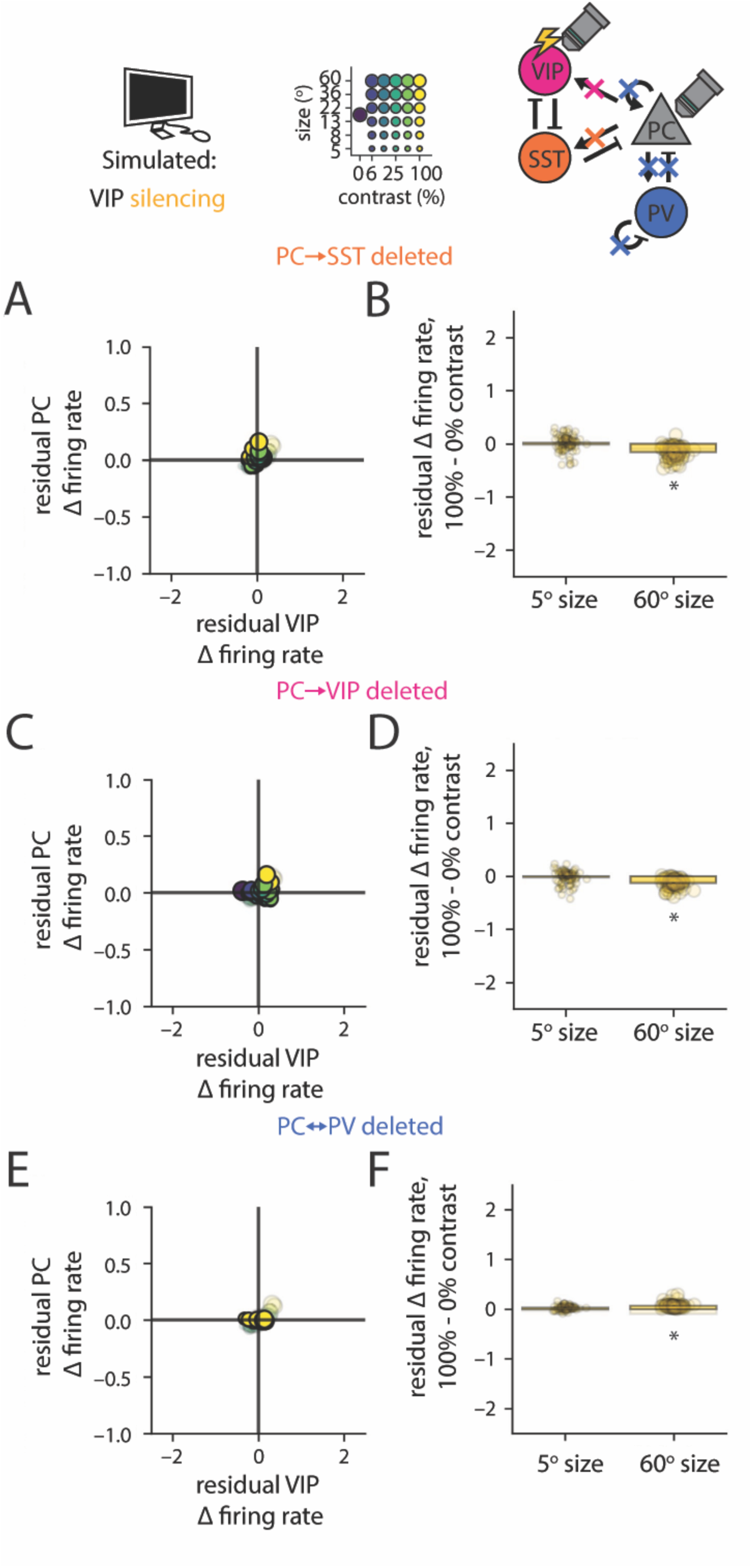
Determinants of VIP-SST competition weakly affect residual effects of VIP silencing. A) Residual change in PC activity, not explained by control firing rate, plotted against the difference in the change in VIP activity from the mean change in VIP activity, for the case of finite excitatory current injection to VIP cells, with PC→SST connections deleted. B) Residual change in PC activity at 100% - 0% contrast, for 5° and 60° size, for the data in B) (difference in residuals significantly reduced by connection deletion, p=2×10^-9^, Wilcoxon signed-rank test). C,D) Same as A,B), but after deleting all PC→VIP connections (difference in residuals significantly reduced, p=6×10^-10^, Wilcoxon signed-rank test). E,F) Same as A,B), but after deleting all PC→PC, PC→PV, PV⊣PC, and PV⊣PV connections. Dots indicate individual model fits (difference in residuals significantly reduced, p=6×10^-7^, Wilcoxon signed-rank test). Note that axes are chosen to match those in fig. 10.

**Figure S17.**
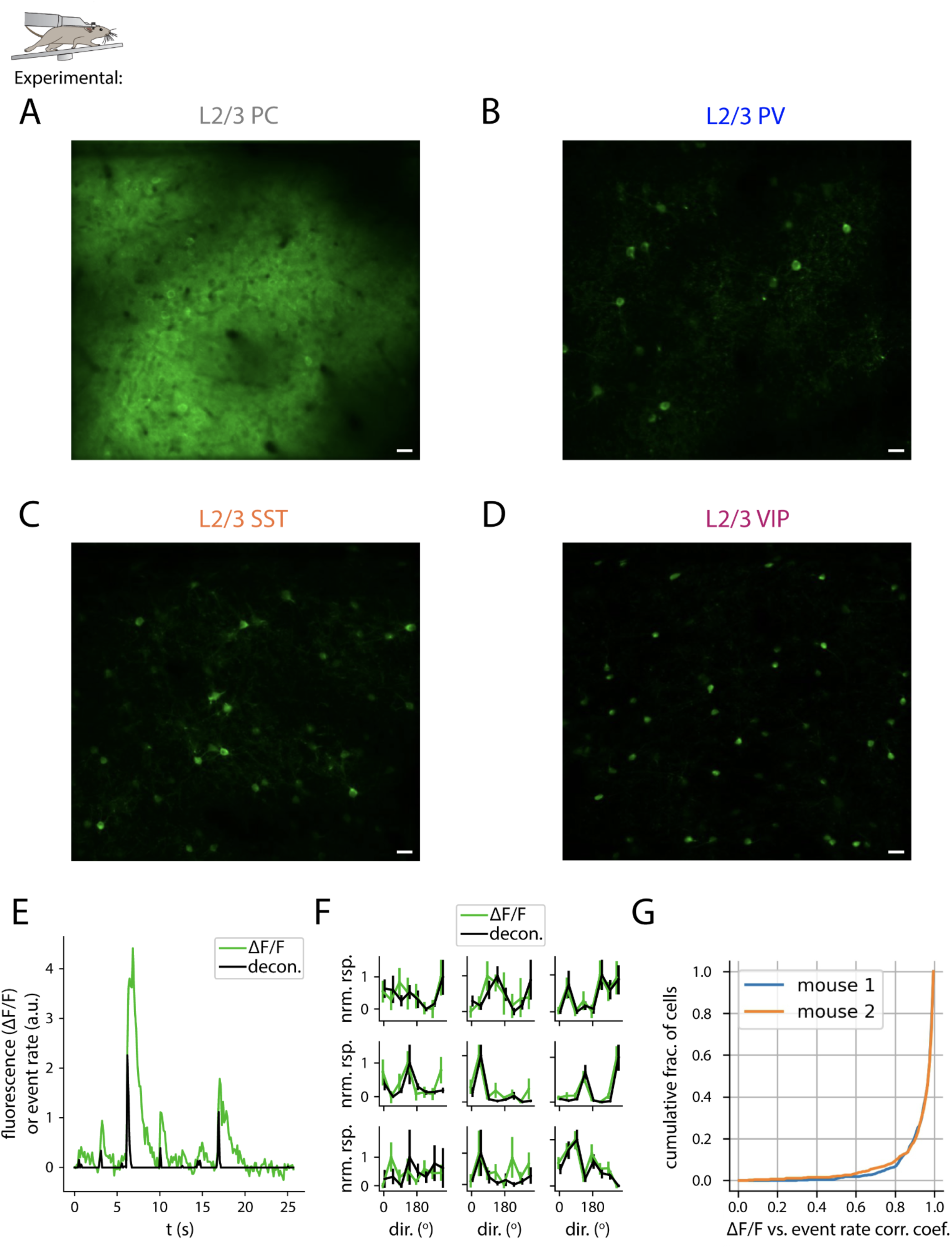
Example fields of view, and deconvolution analysis controls. A) An example imaging plane of a Camk2a-tTA; tetO-GCaMP6s animal, used for recording L2/3 PCs. Computed by averaging motion-corrected frames across an experiment. B-D) Same as A), for PV-Cre; TITL2-GCaMP6s, SST-Cre; TITL2-GCaMP6s, and VIP-Cre; TITL2-GCaMP6s, respectively. E) Example fluorescence (ΔF/F) trace with deconvolved event rate, from a L2/3 PC (same neuron as in fig. 1C). F) In a subset of control experiments, a longer inter-stimulus interval was used, so that evoked calcium transients did not overlap between successive trials. Orientation tuning in nine neurons with reliably estimated ΔF/F tuning curves (R>0.5 between halves of the data, see Methods) is plotted, as measured using ΔF/F and deconvolved event rate. Values are normalized for each to lie between 0 and 1. Error bars represent bootstrap SEM equivalents. G) Cumulative distribution of Pearson’s R between ΔF/F and deconvolved tuning curves for two experiments (analyzing only neurons with reliably estimated ΔF/F tuning curves, as above, *n* = 488/1131, 466/1060, respectively). Error bars indicate bootstrap SEM equivalent.

## Methods

### Experimental model details

All experiments were performed on mice between 1.5 to 14 months of age. CaMKII-tTA mice (RRID:IMSR_JAX:003010) crossed to tetO-GCaMP6s mice (RRID:IMSR_JAX:024742) were used when imaging L2/3 pyramidal cells. Both lines had been outcrossed to the ICR line (Charles River) for several generations. These mice were on a mixed background between outcrossed tetO-GCaMP and camk2-tTA on the C57/B6 background. For L4 PC experiments (resp. L2/3 SST, L2/3 VIP, and L2/3 PV experiments), Scnn1a-Tg3-Cre (resp. Sst-IRES-Cre, Vip-IRES-Cre, and Pv-IRES-Cre) mice were crossed to Ai162(TIT2L-GC6s-ICL-tTA2)-D mice (RRID:IMSR_JAX:031562). For VIP optogenetic and calcium imaging experiments, Vip-IRES-Cre or Vip-IRES-Cre crossed to Sst-IRES-Flp were used. For optogenetic and electrophysiology experiments, Vip-IRES-Cre or Sst-IRES-Cre animals were used. Both female and male animals were used, and maintained on a 12:12 reversed light:dark cycle. All procedures were approved by the Animal Care and Use Committee of UC Berkeley.

### Preparation for *in vivo* two-photon imaging

Headplate attachment, habituation to running on a circular treadmill, and cranial window installation were performed as described previously (Lyall et al., 2021). Briefly, anesthesia was induced with 5% isoflurane and maintained at 1-3% during surgery. Respiratory rate and response to toe/tail pinching was monitored throughout surgery to ensure adequate anesthetic depth. 0.05 mg/kg of buprenorphine was administered subcutaneously for post-operative analgesia, and 2 mg/kg of dexamethasone as an anti-inflammatory. The scalp was disinfected with 70% alcohol and 5% iodine. The skin and fascia above the sensory cortices were removed and Vetbond (3M) was applied to the skull surface and wound margins. A custom titanium headplate was fixed to the skull with dental cement (Metabond). A cranial window was installed to provide for optical access to the cortex. A biopsy punch was used to create a 3.5mm diameter craniotomy over the left primary visual cortex. A window plug consisting of two 3 mm diameter coverslips glued to the bottom of a single 5 mm diameter coverslip using Norland Optical Adhesive #71 was placed over the craniotomy and sealed permanently using dental cement (Orthojet or Metabond). The dental cement was coated in a layer of black oxide to mitigate light leakage during subsequent experiments. Mice were provided at least two days to recover.

For VIP silencing experiments, neonatal Vip-IRES-Cre; Sst-IRES-Flp (P4–6) were briefly cryo-anesthetized and placed in a head mold. Transcranial injection of ∼45nl of undiluted AAV9-CAG-DIO-eNpHR3.0-mRuby3 (UPenn Vector Core) was performed using a Drummond Nanoject injector at three locations in left V1 using a glass pipette beveled to fine tip (∼30–60μm). With respect to the lambda suture coordinates for V1 were 0.0 mm AP, 2.2 mm L and injection was as superficial as possible under the skull.

For VIP activation experiments, Vip-IRES-Cre mice were injected with AAV9-syn-GCaMP6s (UPenn Vector Core) and rAAV9-syn-DIO-ChrimsonR-tdT (UNC Vector Core) virus in left V1. For VIP silencing experiments, Vip-IRES-Cre; Sst-IRES-Flp mice were injected with AAV9-syn-GCaMP6s (UPenn Vector Core) virus in left V1. Briefly, they were anesthetized and administered buprenorphine as described above. A dental drill (Foredom) was used to create a small burr hole 2.75 mm lateral to bregma. Then a WPI UltraMicroPump3 injector was used to inject 300-400 nL of the virus at a rate of 50 nL/min. Post-injection, the needle was left in the brain for 5 minutes to allow the viral solution to absorb into the tissue. Injected mice were provided 2-3 weeks with intermittent head-fixation over the circular treadmill to allow the infected neurons to ramp up expression of GCaMP6s.

### Visual stimulus presentation, *in vivo* imaging, and pupil tracking

For visual stimulus presentation, the monitor was placed 13-15 cm from the eye. Animals were habituated to visual stimulation on the setup for at least two sessions prior to imaging. Before and/or after each experiment, receptive fields were mapped using 10° drifting grating patches. These patches had spatial frequency of 0.08 cycles per degree, and a temporal frequency of 1 Hz, with direction cycling through 0-360° over the course of 1.5 seconds. They appeared at randomly interleaved locations tiling a 40° x 40° visual degree grid, sampled at 5° intervals, with 1.5 second inter-stimulus intervals. In subsequent size-contrast experiments, the visual stimulus consisted of square wave drifting gratings, with directions tiling 0-360° at 45° intervals, with a spatial frequency of 0.08 cycles/ °, and a temporal frequency of 1 Hz (calcium imaging experiments without optogenetics) or 2 Hz (calcium imaging and optogenetic experiments). Visual stimulus presentation lasted one second, followed by a one second inter-stimulus interval. Patch configurations, orientations, and sizes were pseudo-randomly interleaved, and stimuli were generated and presented using the Psychophysics Toolbox (Brainard, 1997). Each distinct visual stimulus was displayed for 5-10 repetitions.

Mice were head-fixed on a freely spinning running wheel under a Nixon 16x- magnification water immersion objective and imaged with a two-photon resonant scanning microscope (Neurolabware) within a light tight box. The imaging FOV was 430 by 670 um, with four planes spaced 37.5 µm apart imaged sequentially using an electrotunable lens (Optotune), sampling each plane at an effective frame rate of 7.72 Hz. For L2/3 imaging, imaging depth was 100 – 300 µm, and for L4 imaging, depth was 350 – 500 µm deep. Electrical tape was applied between the objective and the mouse’s headplate to block monitor light from entering the microscope.

Locomotion was monitored using a rotary encoder, with trials of mean absolute run velocity > 1 cm/sec classified as “locomoting” and < 1 cm/sec classified as “non-locomoting”. Eye movements were imaged using a Basler Ace aCA1300-200um camera, with a hot mirror (Edmund Optics) placed between the eye and the monitor reflecting infrared light for eye imaging while transmitting visible light for visual stimulation. Infrared illumination was provided by the two-photon imaging laser, transmitted through the pupils, as well as a panel of 850nm LEDs (CMVision). Pupil location and diameter were tracked using custom MATLAB code.

### Imaging and optogenetic stimulation

Optogenetic stimulation through the objective was performed with a 617 nm LED (Thorlabs), filtered through a 632/22 nm single-band bandpass filter (Semrock), at an amplitude through the objective of either 6 mW (VIP silencing experiments) or 1.5 mW (VIP activation experiments). Electrical tape between the mouse’s head and the objective served to mitigate direct visual stimulation by the optogenetic light. The PMT was not gated, but protected by a strict shortpass filter. Optogenetic illumination began 0.25 seconds prior to visual stimulus delivery, and ended 0.25 seconds after. Small square wave optogenetic artifacts visible on the PMT were subtracted in post-processing.

### Calcium imaging analysis

Motion correction and ROI segmentation was performed using Suite2p (Pachitariu et al., 2017). Neuropil subtraction was applied as described in (Lyall et al., 2021). ΔF/F traces were calculated as (F – F_0_)/F_0_ with baseline F_0_ computed over a sliding 20^th^ percentile filter of width 3000 frames. Because the inter-stimulus interval was reduced in V1 recordings to permit more stimuli to be displayed, calcium transients overlapped between successive trials. Therefore, we deconvolved calcium traces for this data using OASIS with L1 sparsity penalty (Friedrich et al., 2017), using ΔF/F traces as input.

To confirm that the deconvolution procedure did not distort tuning curve measurements, we performed two experiments with longer inter-stimulus intervals (3 seconds rather than 1 second), varying only stimulus direction, at fixed 15° size and 100% contrast, for 20-30 trials per condition. Because evoked calcium transients did not overlap between successive trials in this condition, tuning curves computed using ΔF/F and deconvolved event rate should agree quantitatively (fig. S17f). To restrict our analysis to neurons for which our estimate of the tuning curve with ΔF/F was robust (i.e., neurons that were visually responsive and orientation-tuned) we randomly split trials into two halves, computed ΔF/F tuning curves for each half, and computed Pearson’s correlation coefficients between the two independent ΔF/F tuning curve estimates. We determined that a neuron’s ΔF/F tuning curve measurement was robust if the two estimates were correlated with R>0.5 (*n* = 488/1131, 466/1060 for the two experiments). For these neurons, we then computed tuning curves based on all trials, using ΔF/F and deconvolved event rates. Indeed, we found that for both mice, ΔF/F and deconvolved estimates of tuning curves were correlated with R>0.9 in ∼80% of neurons for which ΔF/F tuning curve measurement was robust.

### Retinotopic map estimation

Responses to receptive field mapping stimuli were fit using 2D gaussians. The underlying retinotopic map was estimated using linear regression of cell centers (in microns of cortical space) vs. receptive field centers (in degrees of visual space). For subsequent analyses, neurons were separated based on their location on the estimated retinotopic map, relative to the center of the stimulus representation, as indicated in the main text. For each FOV, the retinotopic map orientation relative to the rostral-caudal and medial-lateral axes, and cortical magnification, was confirmed to be consistent with those of V1.

### Tuning curve slope estimation

To compute the local slope of PC event rate/mean with respect to size (or contrast), we divided the difference in responses to adjacent sizes (or contrasts) by the difference in size (or contrast). For intermediate sizes (or contrasts), the slope estimate for a given size (or contrast) corresponded to the average of the two difference terms involving that size (or contrast). On the other hand, for extremal sizes (or contrasts)—meaning the lowest or highest tested—the slope estimate corresponded to the single difference term involving that size (or contrast). Note that although positive contrasts were plotted on a logarithmic scale, linear contrast was used for this estimate.

### Modulation index and slope estimation

Surround modulation index (SMI) was computed as (response to 60° size – response to 0% contrast)/(response to preferred size – response to 0% contrast). Contrast sensitivity index (CSI) was computed as (response to 6% contrast – response to 0% contrast)/(response to preferred contrast – response to 0% contrast). PC SMI (or CSI) was first computed for each recorded neuron individually before averaging to arrive at the within-animal average. To compute the global slope of PC SMI (or CSI) with respect to contrast (or size), we performed linear regression of SMI (or CSI) with respect to the contrast values (or size values) shown.

### Simultaneous imaging of VIP and SST activity using Cre-Flp

We used VIP-Cre SST-Flp mice, injected with GCaMP6s expressed in all neurons. SST cells expressed cytosolic tdTomato in a Flp-dependent manner, while VIP cells expressed Cre-dependent mRuby3 fused to eNpHR3.0. Using calcium imaging, we identified SST cells using their bright, cytosolic red signal, compared with the dimmer, membrane bound mRuby3 in VIP cells; they could be distinguished both by membrane localization and brightness of the red fluorophore (fig. S6).

### Recurrent network model

For a detailed expanation, see Supplementary Note 1. Briefly, we modeled cortical space using two spatial domains. The first represented neurons within ten visual degrees of retinotopic space of the stimulus representation’s center; the second, neurons beyond 10 degrees. Activity in layer 4 pyramidal cells (L4 PCs) in each spatial domain was treated as input to the four recurrently connected layer 2/3 (L2/3) populations in each pixel. The activity of L2/3 populations evolved in time according to:

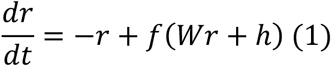

where ***r*** is an 8 (=2 spatial domains x 4 cell types)-dimensional vector of firing rates, ***h*** is a vector of external (i.e., primarily L4) inputs, and *f* is a pointwise expansive nonlinearity (note that the time units in this equation are membrane time constants, assumed to be uniform across cell types). The full weight matrix *W* describing synaptic connections between the four L2/3 cell types in each of the two spatial domains was modeled as the product of a cell type-dependent term, describing the connection strength of cell type *i* onto cell type *j*, multiplied by a term describing the integration across space, of the postsynaptic population *j*. Thus, if *w_ji_* is the synaptic weight from cell type *i* to cell type *j* in the same spatial location, and *c_j_* is the spatial integration factor for cell type *j*, then the synaptic weight from cell type *i* at one spatial location to cell type *j* at the other location is *c_j_w_ji_*. Thus, the full 8×8 matrix of recurrent synaptic connections was described by a 4×4 matrix (the *w*’s), plus a 4-dimensional vector (the *c*’s). A similar procedure was used to generate the full 2×8 matrix of feedforward synaptic connections from L4 PCs. A four-dimensional vector gave the bias current for each L2/3 population. The linear combination representing synaptic input currents was then passed through an expansive nonlinearity. The shape of this nonlinearity was identical for each cell type:

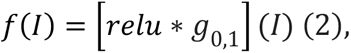

where *g_μ,σ2_* denotes a gaussian with mean *μ* and variance *σ^2^*. Evolution of firing rates in time was simulated according to equation (1) after addition of a small perturbation to ensure stability of the network solution, and for the purpose of this work, we examined time-averaged (“steady state”) behavior over the window of 5-10 membrane time constants after perturbation.

Because ***r*** appears on both the left and right sides of equation (1), we say that solutions for ***r*** must in general be “self-consistent.” For each cell population and each stimulus condition, a stimulus-dependent residual current parameter was introduced, added to the synaptic input and bias currents, so that perfect self-consistency was not required at each step of the optimization. However, squared error penalties on these residual currents ensured that the final network parameters achieved nearly complete self-consistency (see main text). The model fits were randomly initialized, and optimized using L-BFGS-B. The cost function incorporated neuron-level calcium imaging data (fig. S5a), such that model fits were penalized based on their distance from the distribution of single neuron tuning curves. Several synaptic connections (L4 PC→SST, SST⊣SST, and VIP⊣PC) were constrained to be 0, and the PC→PC weight was constrained to be large, in the sense that the product *w_PC->PC_ φ_PC_* > 1, in units where the synaptic leak current is 1, for every stimulus condition (with *φ_PC_* as defined below).

For steady state solutions to equation (1), the change in firing rate *r* expected from a small change in input current *I* can be computed as the product

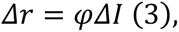

where *φ* is the slope of the firing rate nonlinearity around the steady state solution ***r***. Because *I* is itself a function of ***r***, we were able to simulate the response of a cell population *i* to small current injections to population *j* by computing a “response matrix” *R* as in (Del Molino et al., 2017):

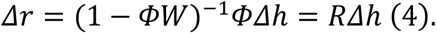

This amounts to linearizing ***r*** about a steady state solution to equation (1). Here, ***Δh*** is a vector of optogenetically induced currents, and is a diagonal matrix whose entries are the values of for each cell type. The matrix *R* yields the theoretical prediction for the effect of small current injections to a given cell population *j*, on any cell population *i*. This matrix changes as a function of stimulus via changes in ***r***, and thus for each cell type.

We leveraged these recurrent network model fits to predict the effects of manipulations on network activity, some experimentally realizable with current technology, and some not. First, we simulated the deletion of certain synaptic connections. For these simulations, we allowed the network to evolve to a new steady state according to equation (1), after setting certain elements of the 4×4 L2/3 synaptic weight matrix to 0, with all other parameters held constant. Second, we simulated the injection of inhibitory (negative) or excitatory (positive) current to specific cell types, modeled as a stimulus-dependent DC offset to the synaptic input current for that cell type. Again, we allowed the network to evolve to a new steady state according to equation (1). In both of these cases, we computed both the perturbed steady state firing rates ***r***, as well as the perturbed response matrix *R*, resulting from the perturbed values for *Φ* and *W*.

For simulations of optogenetic perturbations (silencing and activation) of VIP cells, we tuned the magnitude of current injection for each model fit based on the average of transition currents (see main text and fig. 6) across sizes at 100% contrast. Changes to L4 inputs to the network in response to VIP perturbations were modeled using linear fits to plots of Δ activity vs. activity (firing rate or deconvolved event rate) for recordings of L4 in response to the respective perturbations (see fig. S9).

### Quantification and statistical analysis

Statistically significant differences between conditions were determined using standard nonparametric tests in the SciPy library, including the Wilcoxon signed-rank test and Mann-Whitney *U* test. Analyses were performed on each ROI’s deconvolved event rate for each trial. A single trial’s response was calculated as the average deconvolved event rate during the entire 1 second of stimulation. 95% confidence intervals and SEM equivalents (68% confidence intervals) were generated via bootstrap. For validation of the deconvolution procedure, trials were split randomly into two halves, and tuning curves were computed on each half of trials. Neurons whose tuning curves computed on both halves of the data using ΔF/F agreed with R>0.5 were retained. For these neurons, tuning curves were computed across all trials using both ΔF/F and deconvolved event rate (fig. S17f,g).

### Data Availability

The data that support the findings of this study are available from the corresponding author upon reasonable request.

## Contributions

D.P.M., J.V., A.P., K.D.M., and H.A. conceived the study. D.P.M. performed the calcium imaging experiments. J.V. performed the extracellular electrophysiology experiments. D.P.M. designed and built the recurrent network model, and designed and carried out *in silico* experiments. D.P.M. and H.A. wrote the paper, with input from all other authors.

## Acknowledgements

The authors acknowledge the GENIE Project, Janelia Farm Research Campus, and the Howard Hughes Medical Institute for the GCaMP6 viruses. Thanks to Karthika Gopakumar and Janine Beyer for technical support, and to Mei Li and the Vision Science Core Gene Delivery Module for preparation of AAVs. Thanks to Ruben Coen-Cagli and the members of the Adesnik Lab, particularly Hayley Bounds, Mora Ogando, and Hyeyoung Shin for helpful comments. Thanks also to Charles G. Frye, Sang Min Han, and Benjamin E. Smith. D.P.M. was supported by an NSF Graduate Research Fellowship. A.P. would like to acknowledge the support of the Swartz Foundation Fellowship for Theory in Neuroscience 2019-4 and 2020-6. H.A. is a New York Stem Cell Foundation-Robertson Investigator. This work was supported by The New York Stem Cell Foundation. This work was supported by NEI grant R01EY023756-01 and U19NS97613.

