## Supplementary Note for "Antagonistic inhibitory subnetworks control cooperation and competition across cortical space"

The recurrent network model fit to the data sought to explain tuning curves of recorded neurons using recurrently connected populations of linear-nonlinear model neurons. Five separate populations were modeled: in the input layer, layer 4 pyramidal cells (L4 PCs), which provided only feedforward input to the modeled populations; and four recurrently connected layer 2/3 (L2/3) cell types: PCs, parvalbumin expressing (PV) cells, somatostatin expressing (SST) cells, and vasointestinal peptide expressing (VIP) cells. For each of these populations, we modeled two spatial pixels, with one corresponding to neurons within ten degrees of retinotopic space of the stimulus representation's center, and the other to neurons beyond ten degrees. In all, then, there were $N_{pop} = 8$ recurrently connected L2/3 populations modeled (four cell types by two spatial pixels) and 2 layer 4 populations providing feedforward input (one cell type by two spatial pixels).

### 1 Network Dynamics and Cost Function

#### 1.1 Firing Rate Nonlinearity

Activity patterns in response to a given stimulus were modeled as solutions to a system of equations describing a population of rate neurons asymptotically approaching some firing rate $f(\mathbf{I})$, where $f$ is an expansive nonlinearity and $\mathbf{I}$ is a vector of synaptic currents, with first order kinetics.

$$\frac{d\mathbf{r}}{dt} = -\mathbf{r} + f(\mathbf{I}) = 0 \tag{1}$$

(*2*).

Here, for a model describing $N_{pop} = 4$ recurrently connected cell populations, $\mathbf{r}$ is a $N_{pop}$-dimensional vector where the $i$th entry gives the firing rate of cell type $i$. $\mathbf{r}$ is defined separately for each of $N_{stim} = 36$ stimulus conditions. For the expansive pointwise nonlinearity $f$, we use

$$f_i(x) = F(x, \sigma_0^2), \tag{2}$$

where after (*5*),

$$\begin{aligned} F(\mu, \sigma^2) &= \frac{\mu}{2}\left(1 + \text{erf}(u)\right) + \sqrt{\frac{\sigma^2}{2\pi}}\exp(u^2), \\ u &= \frac{\mu}{\sqrt{2\sigma^2}}. \end{aligned} \tag{3}$$

This is equivalent to a rectified linear unit where the input is convolved with a gaussian of variance $\sigma_0^2$. $\sigma_0^2$ is fixed to 1 for all modeled populations.

#### 1.2 Synaptic Connections

The synaptic currents $\mathbf{I_0}$ coming from the modeled cell types are described by

$$\mathbf{I_0}(\mathbf{r}, \mathbf{h}) = U\mathbf{r} + V\mathbf{h}, \tag{4}$$

where $U$ is a $N_{pop} \times N_{pop}$ matrix describing recurrent weights among the modeled neurons, $V$ is a matrix describing weights from the input layer to the modeled population, and $\mathbf{h}$ is a $N_{inp} = 3$-dimensional vector consisting of the activity in the two input layer populations and a constant (=1) offset term for spontaneous input.

Synaptic connections were constrained to be negative where the postsynaptic cell type was inhibitory, and positive where it was excitatory. Several weights were constrained to be 0 based on prior studies that reported them to be small or absent: SST $\leftarrow$ L4 PC, SST $\leftarrow$ SST, and L2/3 PC $\leftarrow$ VIP.

In order to reduce the number of free parameters and improve interpretability, we modeled the weight matrices, $U, V$, as factorizable into a cell type component and a spatial component. For example, for the connection weight from cell type $j$, spatial pixel $x_2$ onto cell type $i$, spatial pixel $x_1$:

$$
\begin{gathered}
U_{i,x_1 \leftarrow j,x_2} = \tilde{U}_{i \leftarrow j} K_{x,i}(x_1, x_2), \\
K_{x,i}(x_1, x_2) = \begin{cases} 1, x_1 = x_2 \\ k_{x,i}, x_1 \neq x_2, \end{cases}
\end{gathered} \tag{5}
$$

and $\tilde{U}_{i \leftarrow j}$ describes the connection strength from population $j$ to population $i$ within a single spatial pixel. Note that the relative strength of connection across spatial pixels depends only on the postsynaptic population $i$. If $k_{x,i}$ is large, then population $i$ integrates strongly across space.

#### 1.3 Residual Currents

In general, when requiring perfect self-consistency of the recurrent model throughout the fitting procedure, the loss landscape is highly non-convex. To improve convergence, we found it to be empirically useful to introduce additional parameters $\mathbf{I}$, where we enforce

$$
\mathbf{r} = f(\mathbf{I}) \tag{6}
$$

exactly, and

$$
\mathbf{I} \approx U\mathbf{r} + V\mathbf{h} (= \mathbf{I_0}) \tag{7}
$$

with a squared error penalty. Like the entries of $\mathbf{r}$, there is one entry of $\mathbf{I}$ for each cell type, and it is defined separately for each stimulus condition. The error

$$
\Delta\mathbf{I} = \mathbf{I} - \mathbf{I_0} = \mathbf{I} - U\mathbf{r} - V\mathbf{h} \tag{8}
$$

amounts to a residual input current for each cell type and stimulus condition, external to the recurrently connected populations modeled. A possible biological interpretation of these residuals is as currents coming from other differently tuned populations not modeled here (for example long-range input from other cortical areas, or translaminar input from cells in other layers).

#### 1.4 Dynamics Simulation

Solutions to equation 1 can be either stable or unstable. In order to avoid unstable network solutions, we initialized the network at a starting point defined by $\mathbf{r_0} = f(\mathbf{I}) + \boldsymbol{\epsilon}$, with $\boldsymbol{\epsilon}$ a small, random perturbation drawn from a spherical gaussian distribution of variance $\sigma_\epsilon^2$. We then allowed the network to evolve in time (discretized into time steps of duration $\Delta t$ in units of the simulated membrane time constant) according to

$$\begin{aligned} \frac{d\mathbf{r}}{dt} &= -\mathbf{r} + f(U\mathbf{r} + V\mathbf{h} + \Delta\mathbf{I}; \boldsymbol{\sigma}_0^2) \\ \Delta\mathbf{I} &= \mathbf{I} - Uf(\mathbf{I}; \boldsymbol{\sigma}_0^2) - V\mathbf{h} \end{aligned} \tag{9}$$

for $N$ time steps of burn-in, before averaging the subsequent $N$ time steps of $\mathbf{r}$'s evolution to compute $\langle \mathbf{r} \rangle (W, \mathbf{h}, \mathbf{I}, \boldsymbol{\sigma}_0^2 = 1; \boldsymbol{\epsilon})$.

#### 1.5 Cost Function

We fit this model to the first and second moments of the data distribution. We first approximate the distribution of $N_{stim}$-dimensional tuning curves by an $N_{stim}$-D gaussian with covariance given by the first $k = 5$ principal components of the tuning curve covariance matrix, which we call $\mathbf{q}_{data}$. We assume spherical gaussian measurement noise with variance $\sigma_m^2$, and thus the modeled tuning curve distribution is $g_{\mathbf{r},\sigma_m^2}$, denoting a spherically symmetric gaussian of variance $\sigma_m^2$, centered on $\mathbf{r}$. The likelihood of observing this data given a model tuning curve distribution becomes simply $-D_{KL}(\mathbf{q}_{data} || g_{\mathbf{r},\sigma_m^2}) + const.$ A similar cost function was used for the input layer firing rates $\mathbf{h}$, where we fit the first $k$ principal components of the layer 4 PC tuning curve distribution $\mathbf{p}_{data}$ using a spherical gaussian $\mathbf{p}_{model} = g_{\mathbf{h},\sigma_m^2}$. Because $\mathbf{h}$ is independent of other network parameters, we fit the (non-constant) components of $\mathbf{h}$ directly in the optimization.

Some features of model dynamics were also constrained in the fitting procedure. In order to understand the recurrent dynamics predicted by the model, we simulated network responses to infinitesimal current injections in selected cell types, using an $N_{pop} \times N_{pop}$ "response matrix" $R$ as described in (*3*):

$$\Delta \mathbf{r} = (\mathbb{1} - \Phi U)^{-1} \Phi \Delta \mathbf{I} = R \Delta \mathbf{I}, \tag{10}$$

where $\Phi$ denotes a $N_{pop} \times N_{pop}$ diagonal matrix with entries $\Phi_{ii} = f_i'(I_i)$, and $\Delta \mathbf{I}$ denotes a $N_{pop}$-dimensional vector of current injections. This analysis amounts to linearizing equation 1 about a fixed point. Given a small current injection in population $j$, the change in firing rate of population $i$ is proportional to $R_{ij}$. The simulated responses to perturbations depended on network firing rates via the slopes of the pointwise nonlinearities $\varphi_i$.

In keeping with the results of previous optogenetic studies, we constrained the response of center PCs to infinitesimal activation of center SST cells ($R_{PC,SST}$) to be negative (inhibitory) around the steady state for each stimulus condition. To accomplish this, we incorporated a cost term $f_B(-R_{PC,SST})$, where we defined a log barrier function:

$$f_B(u; G) = \begin{cases} -\log u, & u > 0 \\ G - Gu, & u < 0 \end{cases} \tag{11}$$

where $G$ is a large constant.

Using a similar barrier function, we constrained the network to be in an inhibitory stabilized network (ISN) regime across all stimulus conditions, meaning that the PC subnetwork was unstable in the absence of the inhbitory cell types. This is the case when $\varphi_{PC} U_{PC \leftarrow PC} > 1$. Thus, we incorporated the additional term in the cost function $f_B(\varphi_{PC} U_{PC \leftarrow PC} - 1; G)$.

Finally, we incorporated two regularization terms: first, we penalized changes in $\mathbf{I}$ between adjacent sizes and contrasts using a squared total variation (TV) penalty. That is, parameterizing the values $I_i$ associated with cell type $i$ by size $s$ and contrast $c$, we had

$$\begin{aligned} tv(I_i) &= \sum_{s,c} \left( (I_{i;s,c+1} - I_{i;s,c})^2 + (I_{i;s+1,c} - I_{i;s,c})^2 \right) \\ TV(\mathbf{I}) &= \sum_i tv(I_i) \end{aligned} \tag{12}$$

Second, we penalized the elements of $U, V$ using an $L_2$ penalty (i.e., the squared Frobenius norm of each matrix).

The total cost function, then, is written as the negative likelihood of the data given the model parameters, plus the additional penalties described:

$$L(U, V, \mathbf{h}, \mathbf{I}, \boldsymbol{\sigma}_\mathbf{0}^\mathbf{2} = 1; \boldsymbol{\epsilon}) = \beta_r D_{KL}(\mathbf{q}_{data} || g_{\langle \mathbf{r} \rangle, \sigma_m^2}) + \beta_h D_{KL}(\mathbf{p}_{data} || g_{\mathbf{h}, \sigma_m^2}) + \beta_I ||\mathbf{I} - U f(\mathbf{I}; \boldsymbol{\sigma}_0^2) - V\mathbf{h}||^2$$
$$+ \beta_R f_B(-R_{PC,SST}; G) + \beta_{ISN} f_B(\varphi_{PC} U_{PC \leftarrow PC} - 1; G) + \beta_{TV} TV(\mathbf{I}) + \beta_{L2} \left(||U||^2 + ||V||^2\right) \quad (13)$$

The $\beta$-terms, representing the relative weights of terms in the cost function, were tuned empirically by hand:

| | |
|---|---|
| $\beta_r$ | 3 |
| $\beta_h$ | 3 |
| $\beta_z$ | 3 |
| $\beta_R$ | $1 \times 10^{-3}$ |
| $\beta_{ISN}$ | 0.1 |
| $\beta_{TV}$ | 0.1 |
| $\beta_{L2}$ | 0.1 |

with remaining constants:

| | |
|---|---|
| $\Delta t$ | 0.1 |
| $N$ | 100 |
| $G$ | $1 \times 10^6$ |
| $\sigma_m^2$ | 1 |
| $\sigma_\epsilon^2$ | $2.5 \times 10^{-4}$ |

.

### 2 Data Preprocessing

Responses of each neuron to each stimulus were computed using event rate deconvolved from $\Delta F/F$ as described in the methods. Responses were averaged over the eight directions shown, and normalized such that they summed to 1 across stimuli for each neuron. Each neuron's responses were reshaped into an $(N_{size} = 6) \times (N_{contrast} = 6)$ array, and smoothed with a boxcar filter of width 2. For each cell type $i$, responses were then reshaped into an $N_{neuron,i} \times N_{stim}$ array. The mean across neurons was used for the first moment of the "data distribution" $p_i$ or $q_i$ as defined above. This mean was subtracted, and SVD performed on the resulting array. The first $k = 5$ singular values and corresponding dimensions were then used to parameterize the second moments of the "data distribution" across stimulus conditions. In this way, a network solution close by the distribution of measured single neuron tuning curves was less severely penalized than one far from this distribution, even if the distance from the mean were similar.

### 3 Initialization

For random initialization (as used in the first round of optimization; see below), $\mathbf{h}$ was initialized to exactly match measured L4 PC activity, and $\mathbf{I}$ was initialized by inverting equation (6), with $\sigma_{\mathbf{0}}^{\mathbf{2}} = 1$ for all cell types. Inhibitory synaptic connections were initialized from a uniform distribution between -1 and 0, and excitatory connections from a uniform distribution between 0 and 1; as an exception, to ensure the network started in an ISN regime, connections among PCs and PV cells were initialized to:

| | ←PC | ←PV |
|---|---|---|
| PC← | 3 | -5 |
| PV← | 5 | -5 |

For initialization from a previous model fit (as used in subsequent rounds of optimization), all parameters were initialized to their previously fit values, plus random gaussian noise of variance $\sigma^2 = 0.01$.

### 4 Optimization

The model parameters could be categorized into two groups. On the one hand, $U$ and $V$ were low-dimensional, but related to simulated neural activity in subtle ways. One might expect the landscape of the cost function $L$ with respect to these parameters to be bumpy; small changes in $U$ might strongly impact network stability, causing large changes in neural activity. On the other hand, $\mathbf{h}$ and $\mathbf{I}$ were high-dimensional, but related to neural activity in straightforward ways (i.e. simply via the cell-intrinsic firing rate nonlinearity). For this reason, we alternated between optimizing $U$ and $V$, and optimizing $\mathbf{h}$ and $\mathbf{I}$.

For this, we first computed the gradient of $L$ with respect to the parameters of interest using the Autograd Python package (*4*). Note that because computing $\langle \mathbf{r} \rangle$ requires simulating many time steps of the network's time evolution, its gradient amounts to a complicated expression computed using backpropagation through time. Using the cost function and its gradient, we then alternatingly minimized the cost with respect to $\mathbf{h}$ and $\mathbf{I}$ using the conjugate gradient method, and with respect to $U$ and $V$ using L-BFGS-B (*1*), until convergence.

We performed three rounds of progressive optimization: round 1, with random initialization; round 2, initialized from 90th percentile fits of round 1 plus additive noise of variance $\sigma^2 = 0.01$; round 3, initialized from 70th percentile fits of round 2 plus additive noise.

### 5 Simulated perturbations

#### 5.1 Optogenetic perturbation

To model infinitesimal current injections around a steady state, we computed the response matrix as in equation 10. To model finite current injections to a defined L2/3 cell type $i$, we added cell-type specific elements $\Delta I'_i$, of defined magnitude across all stimuli, to the synaptic input current:

$$\mathbf{I}(\mathbf{r}, \mathbf{h}) = U\mathbf{r} + V\mathbf{h} + \Delta\mathbf{I} + \Delta\mathbf{I}' \tag{14}$$

and allowed the network to evolve to a new steady state for each stimulus condition. To model perturbations to L4 PCs, we simply increased or decreased $\mathbf{h}$ by a fixed amount across all stimuli. Note that in all these cases, the cell type- and stimulus-specific residual currents $\Delta\mathbf{I}$ were assumed to be unaffected by the perturbation. To model finite optogenetic perturbations to VIP cells for direct comparison to experimental data, we fixed L4 PC activity to measured values in response to VIP silencing or activation, and additionally simulated finite current injections to VIP cells as described above.

##### 5.2 Synaptic weight deletion

To model the effect of deleting specific connections from the network, we fixed specific elements of the weight matrix $U$ to be 0, and allowed the network to evolve to a new steady state for each stimulus condition, again assuming residual currents $\Delta\mathbf{I}$ to be unaffected. In some cases, we subsequently performed analyses described in section 5.1 using the new steady state activity levels.

2. Peter Dayan and Laurence F Abbott. Theoretical neuroscience: computational and mathematical modeling of neural systems. 2001.

3. Luis Carlos Garcia Del Molino, Guangyu Robert Yang, Jorge F Mejias, and Xiao-Jing Wang. Paradoxical response reversal of top-down modulation in cortical circuits with three interneuron types. *Elife*, 6:e29742, 2017.

4. Dougal Maclaurin, David Duvenaud, and Ryan P Adams. Autograd: Effortless gradients in numpy. In *ICML 2015 AutoML Workshop*, volume 238, page 5, 2015.

5. Kenneth D Miller and Todd W Troyer. Neural noise can explain expansive, power-law nonlinearities in neural response functions. *Journal of neurophysiology*, 87(2):653–659, 2002.
